# Fine-mapping candidate neuropsychiatric regulatory variants using cell type-aware comparative genomics

**DOI:** 10.64898/2026.06.17.732994

**Authors:** BaDoi N. Phan, Alyssa J. Lawler, Jing He, Ashley R. Brown, Irene M. Kaplow, Amanda Kowalczyk, Chaitanya Srinivasan, Grant A. Fox, Rajee Ganesan, Ziheng Chen, Daniel Schäffer, William R. Stauffer, Andreas R. Pfenning

**Affiliations:** Computational Biology Department, School of Computer Science, Carnegie Mellon University; Pittsburgh, PA, USA; Medical Scientist Training Program, School of Medicine, University of Pittsburgh; Pittsburgh, PA, USA; Department of Biological Sciences, Mellon College of Science, Carnegie Mellon University; Pittsburgh, PA, USA; Broad Institute of Harvard and MIT; Cambridge, MA, USA; Neurobiology Department, School of Medicine, University of Pittsburgh; Pittsburgh, PA, USA; Neuroscience Institute, Carnegie Mellon University; Pittsburgh, PA, USA; Allen Institute; Seattle, WA, USA

## Abstract

Measures of nucleotide sequence conservation across species are useful for identifying functional genomic loci, but can fail when regulatory function is maintained, often in a cell type-specific manner, even when sequence is not. We introduce CTACIT, the Cell Type-Aware Conservation Inference Toolkit, to identify trait-associated regulatory variants. CTACIT integrates sequence conservation scores with cell type-specific open chromatin data collected from a few mammalian species to impute function for hundreds more. Applying CTACIT to neuropsychiatric trait loci identifies higher heritability enrichment and more fine-mapped variants than nucleotide conservation and human chromatin data alone. Our *in vivo* reporter assays validate predictions for enhancers with risk variants near the *DRD2* schizophrenia risk locus. By integrating genome conservation and multi-species open chromatin data, CTACIT prioritizes variants within regions of conserved regulatory function for *in vivo* characterization and addresses a major challenge in translating disease associations to mechanistic understanding.

**One-Sentence Summary:** Regulatory functions conserved across mammals underlying caudate cell type evolution reveal functions of human risk variants.

## Main Text

As the number of subjects involved in genome-wide association studies (GWAS) of neuropsychiatric traits increases, a greater number of associated loci are detected [1]. While neuropsychiatric traits share overlapping features, each trait has a unique pathophysiology, prevalence, personal and societal cost, so understanding the precise targets of genetic risk for neuropsychiatric traits will lead to better understanding of each human condition. Across all of these traits, a common feature is that only 4.2% of associated genetic variants are in coding regions [2]. The rest fall outside known protein-coding genes, often in regulatory elements that significantly contribute to the heritability of common traits and disorders [3–5]. To interpret the impact of regulatory SNPs, DNA sequence conservation metrics across primates, mammals, and vertebrates have provided powerful lenses for inferring the risk impact of genetic variation within the human genome [2,5–13]. Across different metrics of genome sequence conservation and varying degrees of power, genetic variation at conserved DNA increases genetic liability for human traits [13]. However, genome constraint approaches cannot resolve which tissues or cell types are implicated in the trait or disease process [14]. This presents challenges for identifying the cell types affected by risk variants in the brain, which has extreme cell type diversity and the role each plays in the pathophysiology of neuropsychiatric traits is still under investigation.

A complementary approach that measures the cell type-specific activity of putative *cis*-regulatory elements using assays like transposase-accessible chromatin with sequencing (ATAC-seq) has proved increasingly useful for interpreting the cell type targets of human traits [13,15–21]. In the past decade, studies profiled cell type open chromatin from a single species to nominate candidate causal cell types affected in human traits. Many works have shown these cell type-specific open chromatin contribute to large proportion of GWAS SNP heritability across a broad spectrum of human traits [21,22]. Studies in non-human model species find that cell type open chromatin mapped to the human genome is highly enriched for human trait liability and is concordant with open chromatin profiled in human cell types [23]. A previous study found that putative human cell type enhancer and promoters regions paired with the genome sequence conservation metric GERP showed higher heritability enrichment across 41-independent human disease and traits [13]. The authors similarly found that putative enhancers and promoters measured in histone ChIP-seq in liver had higher heritability enrichment when those regions had similar histone marks in the livers of other mammals. These studies demonstrate that putative human enhancers and promoters that are in conserved DNA and have measured conserved function across species are valuable to interpret complex human traits.

Christmas et al. showed that DNA sequence conservation in the non-coding human genome is indicative of function in unique cell type or tissue contexts un-profiled by large consortium efforts such as ENCODE. Current ongoing efforts are profiling cell type contexts of the genome in human and other model species. Few of these consortia efforts evaluate how cell type open chromatin may lie on a spectrum of cross-species conservation and how this spectrum would contribute to genetic disease risk. This represents an opportunity to investigate the complex roles of cell type open chromatin conservation within the genome and develop methods to scale multi-species atlases using machine learning and comparative genomics approaches.

In this work, we expand on work related works by Hujoel et al. and others to characterize the conserved open chromatin contributions of diverse neural cell types in the caudate nucleus to identify trait heritability in a panel of neuropsychiatric traits. We first assemble a cross-species caudate cell type open chromatin dataset in human, rhesus macaque, and mouse. When we intersect human cell type open chromatin profiles with new measures of mammalian genome sequence conservation, a 240-mammal PhyloP scores and 43-primate PhastCons score [24] we reveal that neuropsychiatric trait heritability is both cell type-specific and increased in the open chromatin of higher conserved genomic regions. We also introduce new usage for the machine-learning based cell type-aware conservation inference toolkit (CTACIT) that takes regions from genomes of 240 mammals to predict which human cell type open chromatin are still active in which mammalian species better than intersecting human cell type open chromatin and genome conservation scores alone. We use CTACIT-predicted open chromatin conservation across species to fine-map candidate causal variants to conserved, active cell type open chromatin regions and validate the conserved cell type-specific activity with *in vivo* reporter assays of enhancers near the *DRD2* locus containing highly pleiotropic fine-mapped SNPs. In summary, neuropsychiatric traits across multiple domains have complex cell type-specific patterns of trait heritability enrichment and genetic variation at cell type open chromatin that retain this function across mammalian species reflect higher genetic risk than those cell type open chromatin that lose cell type function across mammalian species.

## Results

To provide a foundation to investigate comparative cell type epigenomic analyses of neural cell types with neuropsychiatric disorders, we assembled a cross species atlas of chromatin accessibility of the caudate nucleus, a brain region with well-established cell type homologies [16,25] that has been heavily implicated in psychiatric disorders, addiction behavior, and sleep behavior [16,26–29]. We performed single nucleus assays for transposase-accessible chromatin (snATAC-seq) experiments in rhesus macaque (**Figure S1A-B**) and combined this with published snATAC-seq from human and mouse [15,16,19]. We found 8 caudate cell types at a resolution with matched orthologs across humans, rhesus macaques, and mice (**Figure 1A**, **S1C-F**, *Supplemental Text, Cross-species caudate cell type ortholog identification*). The neuronal subtypes consist of interneurons, the majority of which are *PVALB*^+^, canonical medium spiny neurons (MSNs) marked by dopamine receptors D1 and D2 [25,30], and a distinct population of neurons that shares features with D1/D2-hybrid medium spiny neurons, also known as eccentric medium spiny neurons, previously reported in primates, rodents, and bats single cell transcriptomic studies [19,31–34].

**Figure 1:**
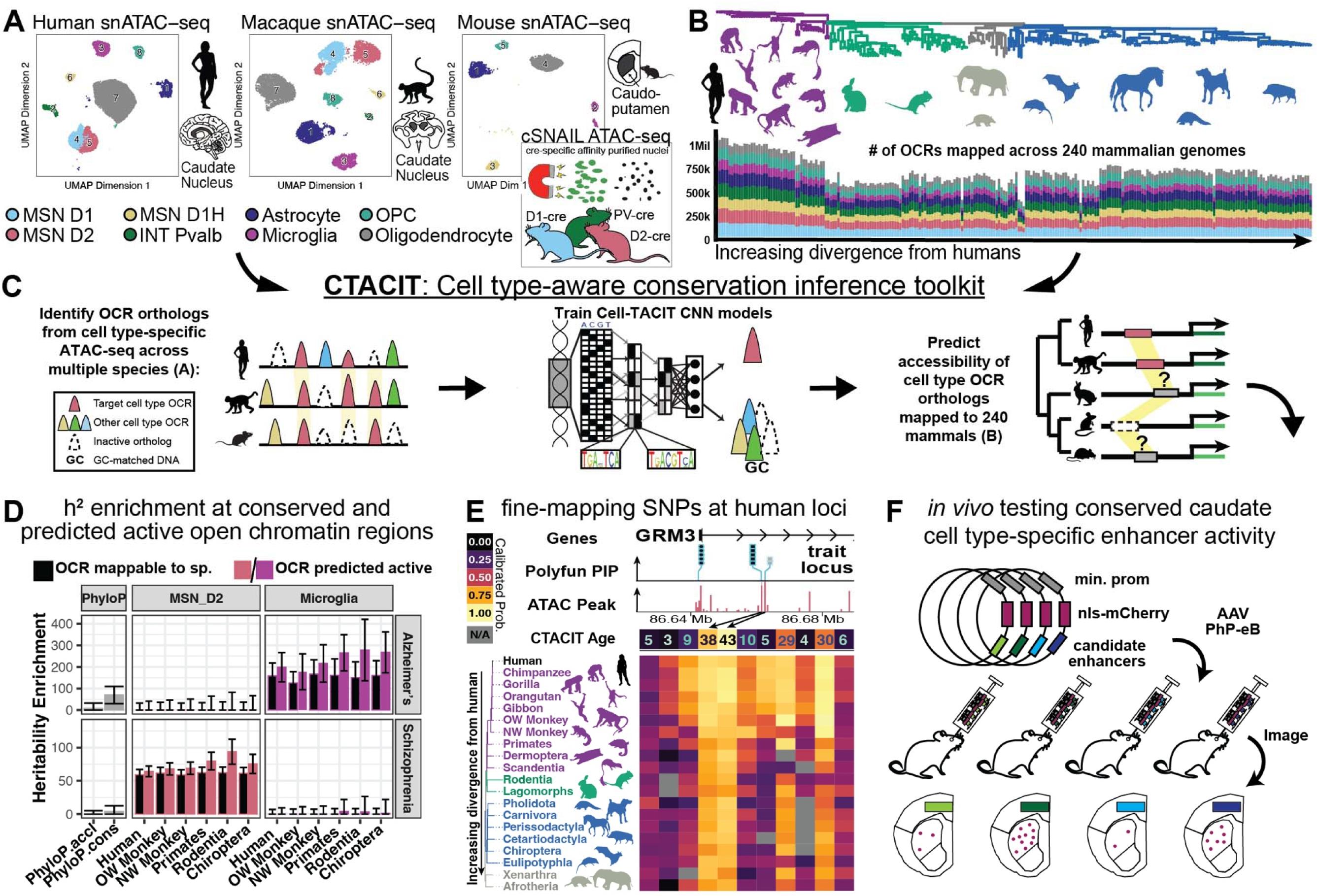
Interpreting genetic variants of complex neurological disorder traits with machine learning models inferring conserved functional genomes of caudate cell types across species. **(A)** Eight cell types identified across the primate caudate nucleus and mouse caudoputamen. Reduced dimensionality UMAP projection plots of single nucleus assay for transposase-accessible open chromatin (snATAC-seq) from rhesus macaque performed in this study alongside reprocessed snATAC-seq data from human, and mouse. *Cre*-dependent Sun1GFP Nuclear-Anchored Independent Labeling (cSNAIL) ATAC-seq conducted in transgenic mouse strains, sub panel. **(B)** The inferred molecular Zoonomia phylogeny aligned to a stacked histogram showing the number of human caudate cell type open chromatin regions (OCRs) that are mappable across placental mammals. The phylogenetic tree is rooted with *Homo sapiens* as the outgroup. Tree branches are colored by the species membership in distinct grand orders and magna orders with representative animal silhouettes from phyloPic.org. **(C)** Cell Type-Aware conservation inference toolkit (CTACIT) pipeline to build convolutional neural network (CNN) models of cell type-specific regulatory activity using DNA sequence underlying OCRs and use models to predict open chromatin at OCR orthologs mapped across species. **(D)** Simplified genome-wide heritability enrichment barplot using CTACIT predicted conserved open chromatin across placental mammals. This example shows D2 MSN and microglia OCRs mappable across species, respective CTACIT predictions, and mammalian accelerated or constrained regions (phyloP.accel and phyloP.cons) with Alzheimer’s Disease (AD) and schizophrenia GWAS. These panels are plotted in full in Figure 2. **(E)** Simplified track plot of the *GRM3* gene locus with fine-mapped SNPs interpreted by leveraging CTACIT predicted conserved open chromatin across placental mammals. Composite subplots display SNP and OCR tracks and corresponding heatmap shows cross-species OCR CTACIT predicted activity. Blue track: lollipop plots of fine-mapped SNPs from the Neuroticism GWAS. The lollipop feature height reflects the maximum PIP of the fine-mapped SNPs, and features are jittered to better visualize clusters of fine-mapped SNPs across a large genomic region. Red track: human D2 MSN OCRs tracks with heights reflecting the CTACIT Age of each OCR. Arrows align cell type OCR tracks to corresponding columns in the heatmap. Heatmap entries are the average CTACIT calibrated scores of species within the same clade/divergence group from humans. OCRs not mappable to a clade are plotted in gray. **(F)** Schematic of testing CTACIT predictions of conserved cell type-specific gene-regulatory activity with an *in vivo* reporter assay using adeno-associated virus (AAV) and microscopy. MSN D1, canonical medium spiny neurons with D1 receptors; MSN D2, canonical MSNs with D2 receptors; MSN D1H, D1/D2 hybrid MSNs; INT, interneurons that largely *PVALB+*; OPC, oligodendrocyte precursor cells; h^2^, narrow sense SNP heritability; phyloP.accel, genomic regions accelerated in mammals; phyloP.cons, genomic regions conserved in mammals; OW Monkey, old-world monkeys/catarrhine clade; NW Monkey, new-world monkeys/platyrrhine clade; Primates, strepsirrhine clade; Rodentia, rodents clade; Chiroptera, bats clade; Sp., species. Mb: million bases.

Having established matched cell type OCRs across species, we then were able to ask whether these annotations capture trait heritability beyond what is explained by nucleotide-level sequence conservation alone. We applied stratified-linkage-disequilibrium score regression (S-LDSC) on common frequency SNPs with open chromatin region of caudate cell types, hereby referred to as cell type OCRs, in 52 neuropsychiatric traits spanning neurodegeneration, psychiatric disorders, substance abuse disorders, sleep and neurological traits, and 12 non-brain traits to assess how cell type epigenomic features of the genome compare with genome sequence conservation (**Figure 1D, Data S4**). Across our brain and non-brain traits, the heritability enrichment of conserved genomic regions measured by a Zoonomia 240 mammals PhyloP (score > 3.51, [2]), or Zoonomia 43 primates PhastCons score (score > 0.994, [2]) tend not to be enriched in neuropsychiatric or have lower heritability enrichments than human caudate cell type open chromatin (Figure 1D). We took a similar approach to Sullivan et al. [2] and computed how the simple average of these genome sequence conservation scores in human cell type open chromatin would stratify trait heritability enrichment.

We find that conditional independent LD score regression effect sizes, τ*, conditioned on the baseline annotations and a background set of cell type OCRs (Methods), are larger in shared human-macaque or human-mouse cell type OCRs than OCRs from any one species alone (**Data S5,** *Supplemental Text, Pairwise cross-species LDSC regression*). This suggests that the conservation of cell type-specific open chromatin between human and other mammals can help to finemap loci identified through GWAS for neuropsychiatric disorders.

We first estimated SNP-heritability using nucleotide conservation metrics, specifically phyloP [11] computed from 240 placental mammal genomes. These phyloP scores showed unprecedented power to interpret human trait-associated loci [2] but had inconsistent heritability enrichment for neuropsychiatric traits (**Figure 1D**, left, phyloP.cons from 240 mammals, Alzheimer’s Disease heritability enrichment = 69.7; schizophrenia HE = 3.82) [24]; because smaller annotations tend to inflate enrichment, these magnitudes are best read as relative rankings (Limitations). In the case of schizophrenia, others have speculated that evolutionary annotations are not capable of substantially explaining the heritability of schizophrenia due to the human-specificity of the phenotype of implicated loci [35–38] despite success from analyses from mouse brain single-cell transcriptomic profiles to identify putative cell types underlying schizophrenia [27,28,39]. The framework of CTACIT provides that opportunity to compare two alternative models for the evolution of regulatory elements associated with schizophrenia and other neuropsychiatric traits: do the regulatory functions of these regions emerge anew in the human lineage, or do they have deeply conserved function that is often missed by traditional methods?

Despite several hundred million years of divergence, some genomic elements retain cell type-specific gene regulation across Metazoa by shuffling and reforming critical transcription factor-binding sites [40], suggesting that borrowing information across species could help make up for the absence of extensive empirical data on gene regulatory function. Since conservation of regulatory function does not always require nucleotide-level sequence conservation [14,40,41], machine learning models are useful for inferring functional conservation from nuanced sequence patterns. We previously demonstrated that convolutional neural networks learn transcription factor binding sites related to cell type-specificity [16] and accurately identify orthologous DNA sequences across species that retain tissue-specific open chromatin activity [14].

We present Cell Type-Aware Conservation Inference Toolkit (CTACIT), a machine learning framework that learns the conserved functional activity of non-coding genomic sequence from cross-species matched cell type OCR and imputes regulatory activity across hundreds of placental mammals included in a reference-free genome alignment (**Figure 1A-C**). By cell type OCR we mean a reproducible open chromatin region identified within a given caudate cell type; specificity relative to other cell types is handled separately in model training through the cell type negative set. In [42], we demonstrate our ability to predict differences in tissue- and cell type-specific open chromatin at orthologous genomic regions of other species and use those predictions to identify regulatory loci undergoing convergent evolution similarly to our prior work [42]. Here, we introduce a new application to marry comparative genomics and cell type-specific experimental evidence to infer the impacts of non-coding genetic variation in neuropsychiatric human traits. Throughout, we refer to the per-species calibrated probability that an OCR ortholog is active as the CTACIT Score, the binary active/inactive calls derived from these scores in each clade as CTACIT predictions or annotations, and the divergence-weighted summary of predicted activity across the placental phylogeny as the CTACIT Age (defined below).

The striatum is known in neurobiology to be a hub of many psychiatric disorders, addiction behavior, and sleep behavior. Previous studies using tissue level heritability enrichment find many neuropsychiatric traits have enrichment around human caudate genes [5,7], we explored OCRs from caudate cell types across multiple species on a panel of neuropsychiatric conditions. We mapped human OCRs across 240 Zoonomia genomes to find conserved cell type regulatory grammar, i.e., combinations of transcription factor binding sites that drive expression (**Figure 1B**) [43,44]. The OCRs of each cell type encompass many cell type-specific transcription factor binding motifs across species (**Figure S2, Data S1)**.

We collected a matched dataset of cell type OCRs from caudoputamen or caudate of mouse and rhesus macaque, respectively, to assess cell type epigenomic conservation compared to human. We selected rhesus macaque and mouse because matched cell type open chromatin from a primate and a rodent span roughly 90 million years of divergence while retaining identifiable caudate tissue and cell type homologies, the divergence range over which CTACIT needs to learn conserved regulatory codes. We mapped the mouse and rhesus cell type open chromatin to human and each other’s genome and compared whether cell type-specific open chromatin regions were conserved across species. Despite the 90 million years of evolutionary distance, caudate cell type-specific OCRs retain cell type-specific function. The subset of human caudate cell type OCRs mappable to rhesus macaque or mouse genomes overlap OCRs measured active in those species (**Figure S3)**. Previous methods to integration of multi-species single cell epigenomic data use computed gene-based metrics and then 1-1 gene orthology. These data show caudate neural cell types retain distinct regulatory codes within the Euarchontoglires genomes, which based on our previous works [14,42], indicate that CTACIT models would likely infer cross-species open chromatin status from orthologous DNA sequences. We trained CTACIT models for each neural cell type and demonstrated their accuracy at predicting cell type-specific open chromatin status of orthologous DNA across species (**Figure 1C**, **Figure S4-S8**, **Data S2-3**, *Supplemental Text, CTACIT performance evaluation*). Model training occurs over positive examples from tens of thousands of cell type OCRs and folds more in negative OCRs from off target cell types and random GC-match genomic non-enhancer regions per species. Our model evaluations demonstrate, as does the prior work, [14,42] that CTACIT can be trained with three species from cell type OCRs. We empirically find our model performance has high auROC for cell type OCRs across species and cell types across varying evaluation datasets (**Figure S6**).

Using CTACIT predictions of caudate OCRs, we created conserved cell type open chromatin annotation tracks for each clade of mammals increasingly diverged from humans (**Data S6**). We show in both brain- and non-brain related traits that cell type OCRs with orthologs predicted to be active in distant species had higher heritability enrichment than other human cell type OCRs (**Figure 2**, colored bars; **Figure S11-26** “CTACIT Score’’). We found an increase in heritability enrichment in the cell type of likely etiology, such as Alzheimer’s Disease (AD) with microglia (HE_AD-Micro_ = 177 - 278) [45,46], schizophrenia with D2 MSNs (HE_SCZ-D2_ = 64.1 - 93.8) [39], and SmokingInitiation with D1 MSNs (HE_SmokInit-D1_ = 19.5 - 43.1) [47]. Since it may be more difficult to map OCRs to species more distantly related to humans, this trend may have been driven by biases in evolutionary distance instead of biologically meaningful differences across species. We therefore confirmed that heritability enrichments are not higher due to mappability to distant species (**Figure 2**, black bars, HE_AD-Microglia_ = 126 - 166; HE_SCZ-D2_ = 58.8 - 62.2; HE_SmokInit-D1_ = 20.8 - 24.2) [24]. Any mappable OCR to distant mammalian species has heritability enrichments substantially higher than enrichments calculated when using single-nucleotide PhyloP conservation (**Figure 2**, left), however this could be due to smaller annotation set size. Overall, the heritability enrichments from CTACIT annotations suggest that the conserved regulatory code of distinct cell types, rather than nucleotide-level genome sequence conservation, holds considerably more genetic heritability for neuropsychiatric traits.

**Figure 2:**
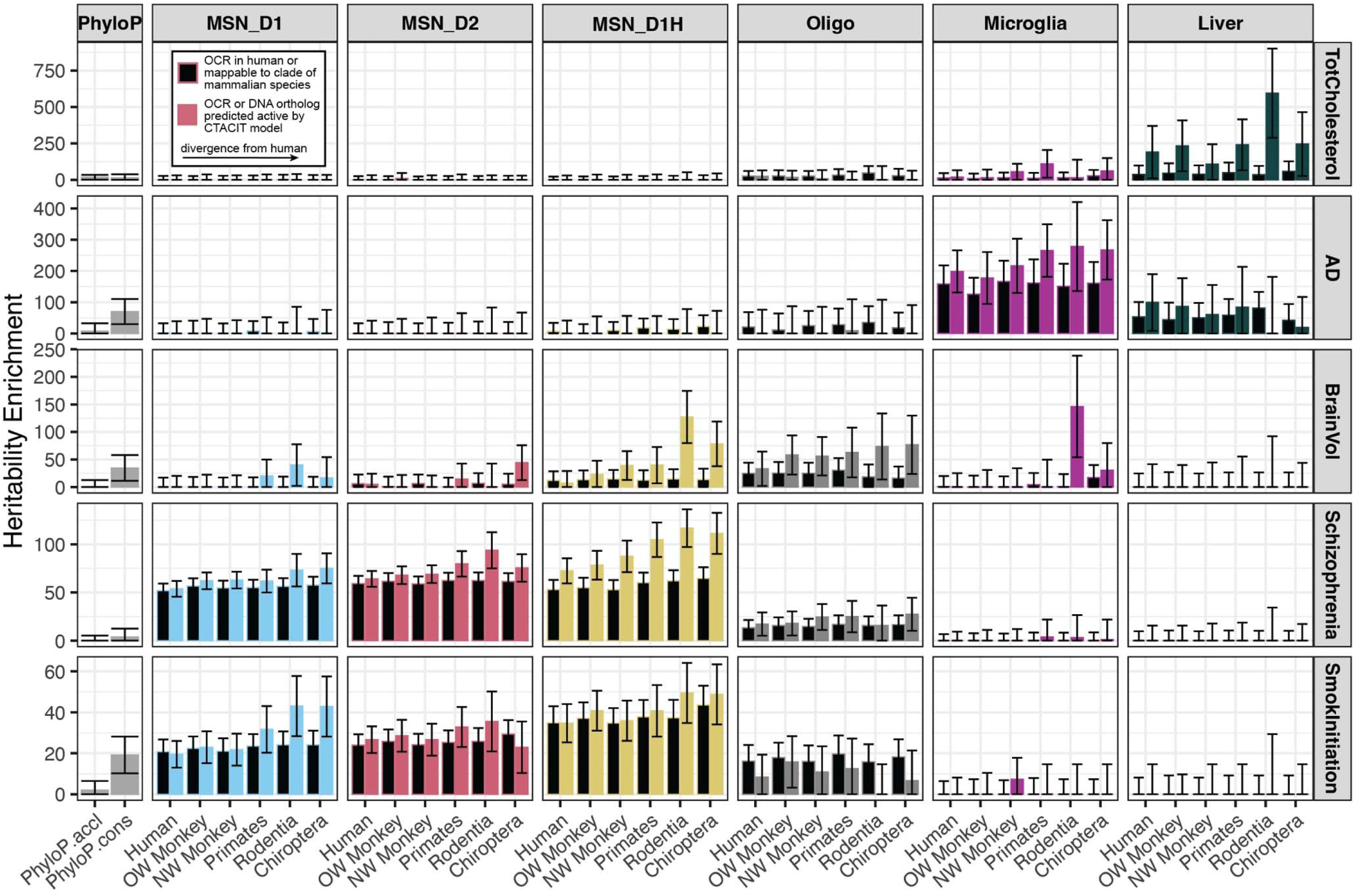
Cell type-aware machine learning predicts regulatory evolutionary conservation to interpret cell type specificity neuropsychiatric traits. Barplots of estimated heritability enrichment (±standard error) of genome-wide association studies in cell type open chromatin regions (OCRs) that are mappable to distant mammalian clades (black bars), predicted active by CTACIT models (colored bars), and regions of mammalian nucleotide conservation or acceleration (PhyloP, left). Traits from top to bottom: total blood cholesterol (TotCholesterol), Alzheimer’s Disease (AD), brain volume (BrainVol), schizophrenia, and current smoker (SmokInitiation). OW Monkey: Catarrhini, species diverged from humans at the branch for old-world monkeys, NW Monkey: Platyrrhini, species split from humans at the branch for new-world monkeys; Primates: Strepsirrhini, species split from humans at the branch for primates. CTACIT Models from MSN_Ds1, D1 medium spiny neurons; MSN_D2, D2 medium spiny neurons; MSN_D1H, D1/D2 hybrid medium spiny neurons; phyloP.accel, genomic regions accelerated in mammals; OW Monkey, other old-world monkeys/catarrhine clade; NW Monkey, new-world monkeys/platyrrhine clade; Primates, other primates/strepsirrhine clade; Rodentia, rodents clade; Chiroptera, bats clade.

We summarized the predicted conserved activity across placental mammals of each non-coding, distal open chromatin region into the CTACIT Age, a metric capturing the million years divergence from humans weighted by predicted activity (**Figure 1E**). In essence, the CTACIT Age measures the millions of years that a human OCR has persisted to retain activity in a cell type of another species. We conducted an orthogonal set of heritability enrichment analyses excluding coding regions and promoters, which have different levels of evolutionary constraints, and binning open chromatin regions into equal CTACIT Age quartiles to control for annotation size. Despite the lower power for genome-wide enrichment analyses when using fewer OCRs per bin, we find that neuropsychiatric traits have higher heritability enrichments in “older” Age quartiles. Such traits include self-reported depression in D2 MSNs, age of smoking initiation in D1 MSNs, and cross-psychiatric disorders in D1 Hybrid MSNs (**Figures S11-26**, “CTACIT Age” columns). The full set of heritability enrichment analyses comparing various cross-species snATAC-seq, Zoonomia PhyloP, and CTACIT annotations are reported in the Supplement (**Figure S11-26, Data S7**).

We examined the value of CTACIT Age to identify functional genetic variation by comparing to two expression quantitative trait loci (eQTL) datasets [48,49]. We first looked at Age quartiles that overlap eQTLs from the caudate nucleus from the GTEx Consortium (**Methods**). We found that caudate GTEx eQTLs were uniformly enriched in the “youngest” Age quartiles of all cell types but mostly depleted from “older” Age quartiles (**Figure 3A**, logistic regression Bonferroni p-value < 0.05, **Data S8**). Since CTACIT models were at the resolution of neural cell types, the lack of enrichment in “older” Age quartiles may be due to differences between bulk tissue and single cell gene expression. We then compared eQTLs (FDR < 0.05) from single–cell RNA-seq of the human prefrontal cortex [49] where four major glial cell types have one-to-one correspondence with our models. The matched CTACIT models had the top enrichment for cell type eQTLs (**Figure 3B**, Bonferroni p-value < 0.05, **Data S8**). This effect was notable for microglia and oligodendrocyte eQTLs. Astrocyte eQTLs had lower enrichment for astrocyte Age quartiles, possibly due to astrocytes having brain region-specific gene expression, especially between the prefrontal cortex and the caudate nucleus [50,51]. Interestingly, “older” Age quartiles in matched cell types have decreasing enrichment for eQTLs–suggesting that predicted “younger” regulatory regions have less time for negative selection and enjoy variable effects on gene regulation as presented by [52]. For cell type eQTLs that do not have a matching CTACIT model, “older” Age quartiles are not enriched for eQTLs (**Figure S28A**, Bonferroni p-value < 0.05). While imperfect, comparing to brain and cell type eQTLs demonstrates that CTACIT Age is specific at ranking genetic variants with regulatory function.

**Figure 3:**
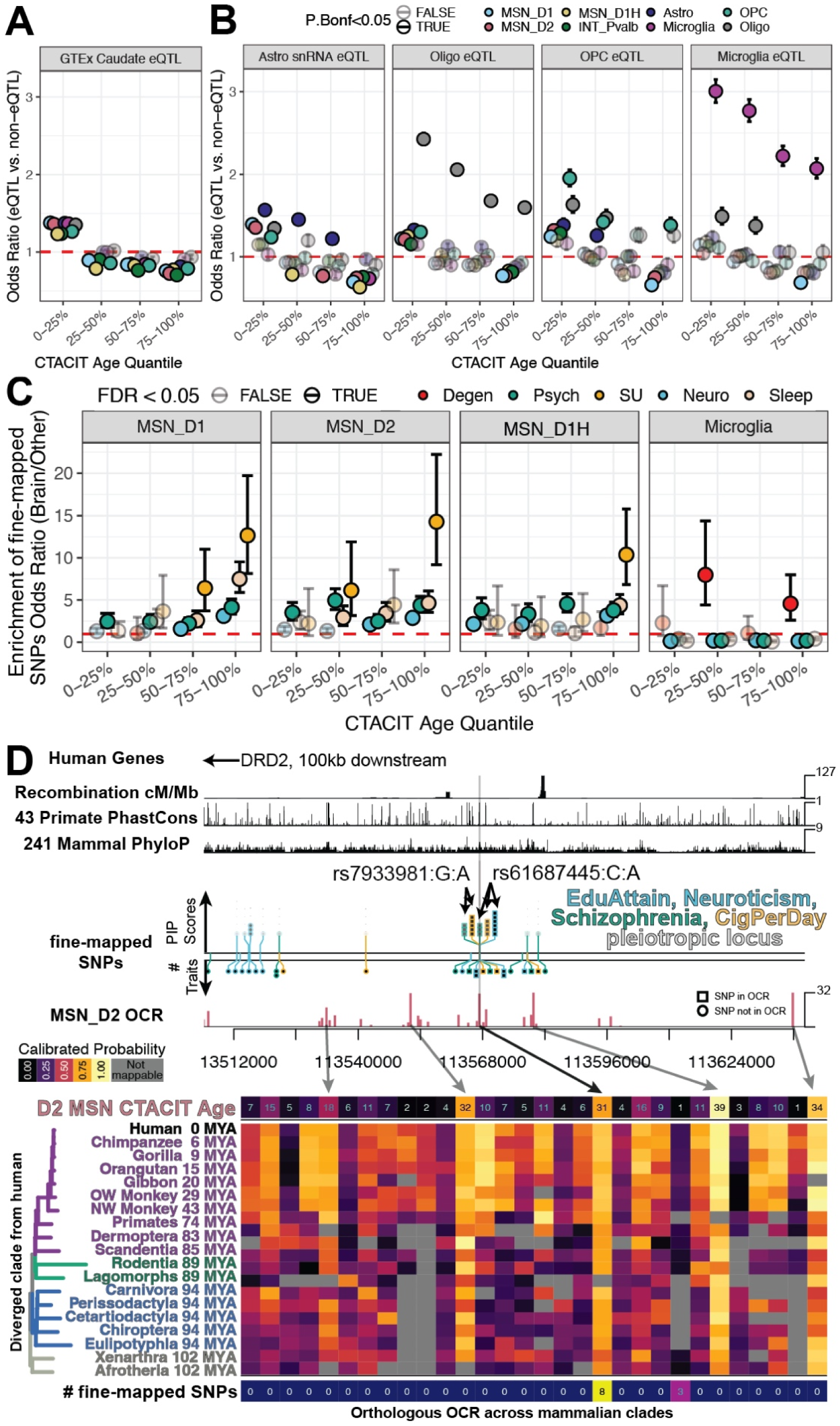
Annotating cell type eQTLs and fine-mapped SNPs with CTACIT Age estimates conserved cell type regulatory activity across mammalian species. **(A)** Stripchart plotting the odds ratio (±standard error of the mean) that significant eQTLs from bulk caudate nucleus RNA-seq are enriched in caudate cell type open chromatin regions stratified by the CTACIT Age. The top Age quantiles are regions predicted to be “older” and retain activity in distant placental mammals. Each point is colored by the corresponding CTACIT model. Solid colored points indicate significant enrichment at Bonferroni corrected P-value < 0.05. **(B)** Enrichment stripchart eQTLs from single-cell prefrontal cortex cell types with CTACIT Age quartiles. These prefrontal cortex cell types have one-to-one correspondence with a CTACIT cell type model. Cell types without one-to-one correspondence are shown in **Figure S28A**. **(C)** Enrichment stripchart for fine-mapped neuropsychiatric trait SNPs with PIP > 0.10 in CTACIT Age quartiles for the top caudate neural cell types. Each point is colored by enrichment for SNPs fine-mapped in neurodegenerative traits (Degen), psychiatric traits (Psych), substance use traits (SU), neurological traits (Neuro), or sleep traits (Sleep). Other CTACIT cell types are shown in **Figure S28B**. Solid points are significant enrichment at FDR corrected P-value < 0.05. **(D)** Locus plot 100kb downstream from the *DRD2* gene with SNPs and CTACIT annotations for D2 MSN models identifies candidate regulatory elements with fine-mapped SNPs from education attainment, neuroticism, schizophrenia, and cigarettes per day GWAS. CTACIT Age (top row) heatmap for D2 MSNs summarizes the predicted activity of each open chromatin region in the track plot across increasingly distant clades of placental mammals (middle rows). The phylogenetic tree annotates the millions of years ago (MYA) that the common ancestor of each group of placental mammals and humans has diverged. Two D2 MSN open chromatin regions contain fine-mapped SNPs in this locus (bottom row).

We used the CTACIT Age to prioritize SNPs from the subset of neuropsychiatric traits with moderately- and well-powered GWAS. We performed functionally informed fine-mapping using our cross-species caudate cell type OCRs and updated annotations of primate and mammalian genome sequence constraint (**Figure S27, Data S9-10,** *Supplemental Text, PolyFun Fine-mapping*) [2,53]. Fine-mapped neuropsychiatric trait SNPs relative to other annotations or non-brain traits are more likely to be enriched in “older” CTACIT Age quartiles (**Figure 3C**, FDR < 0.05, **Data S8**). “Older” medium spiny neuron Age quartiles had the highest enrichments for SNPs associated with psychiatric, sleep, substance abuse, and general neurological traits. In addition, two microglia Age quartiles were enriched for neurodegenerative disorder-associated SNPs. Fine-mapped neuropsychiatric trait SNPs were not enriched in other neural cell types, likely indicating they are not the etiological targets of risk variants (**Figure S28B,** FDR > 0.05). Furthermore, the opposing trends of enrichment in CTACIT Age regions between fine-mapped GWAS SNPs and eQTLs may be due to differences in selection pressure on genetic variation contributing to human disorders and unassociated variants [52,54].

We focused on the *DRD2* locus because it is pleiotropic across several neuropsychiatric traits and because statistical fine-mapping, nucleotide conservation, and public epigenomic annotations gave conflicting evidence about which variant is causal, the setting where cell type-aware prediction is most informative. A cluster of pleiotropic SNPs near the *DRD2* locus were fine-mapped as candidate causal SNPs for multiple traits [39,47,55,56]. Dopamine signaling and changes in dopamine receptor D2 availability in the caudate nucleus have long been associated with schizophrenia, addiction traits, and depression [57–62], with recent findings showing that mouse MSN marker genes were enriched for schizophrenia risk SNPs [27,28]. These SNPs overlap many D2 MSN OCRs, some predicted to remain active across placental mammals and some predicted to lose activity in species diverged from humans (**Figure 3D**). The top two fine-mapped SNPs, rs7933981 and rs61687445, overlap the same open chromatin region, which is accessible across MSNs (**Figure 4A**, Candidate A). A nearby SNP, rs7928017, is in perfect linkage disequilibrium with the top variants and overlaps another MSN-accessible OCR (**Figure 4A**, Candidate B). All three SNPs are eQTLs for *DRD2* in the postmortem human brain [48].

**Figure 4:**
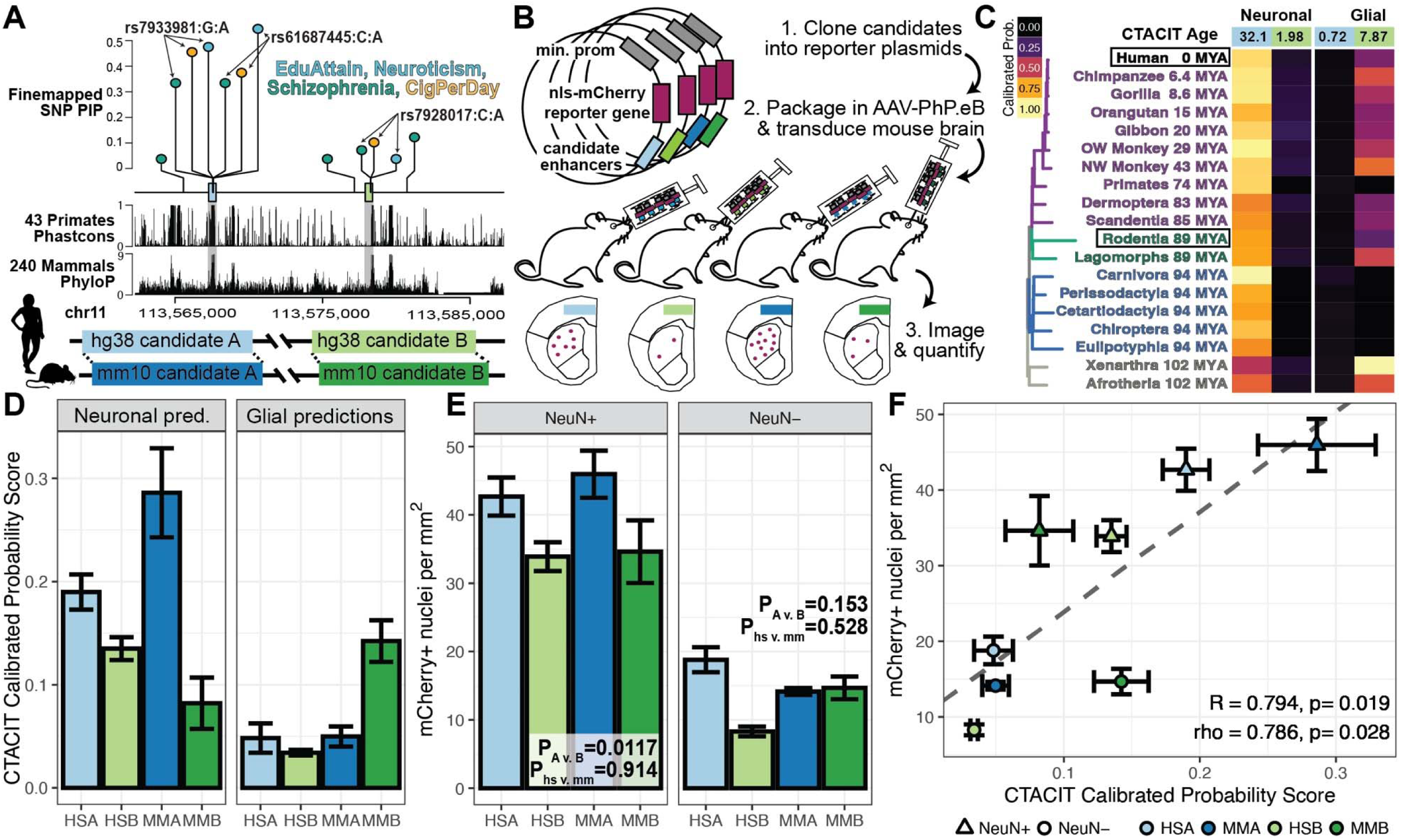
CTACIT accurately predicts conserved cell type-specific regulatory activity in *in vivo* reporter assays of candidate enhancers containing fine-mapped neuropsychiatric variants. **(A)** Zoomed in lollipop plots of Figure 3D showing fine-mapped pleiotropic single nucleotide polymorphisms (SNPs) within two candidate enhancers A and B upstream of the *DRD2* locus. Below the lollipop plots are primate PhastCons and mammal PhyloP nucleotide conservation genome tracks. The schematic showing the orthologs of human candidate enhancers (light blue and light green) in the mouse genome (dark blue and dark green) is shown on the bottom. **(B)** Schematic showing adeno-associated virus (AAV) PhP.eB-based *in vivo* reporter assays to test conserved cell type regulatory activity of the human and mouse ortholog candidate enhancer sequences in the mouse striatum. **(C)** Heatmap of the average CTACIT Scores (calibrated predicted probability), weighted by the proportion of neurons or glia, for the two candidate enhancers in humans and species increasingly diverged from humans. The scores predicted used the full candidate sequences that were mappable across 240 placental mammals. The CTACIT Age across mammals is reported in top blue and green boxes for candidate enhancer A and B, respectively. **(D)** Barplot of the average CTACIT Score (±standard error of the mean, SEM, N = 4), weighted by the proportion of neurons and glia. These scores are predicted using the shorter sequences used to clone the reporter assay plasmids. **(E)** Barplot of the average number of mCherry+ nuclei per mm^2^ (±SEM, N=3) that were co-positive for the neuronal marker NeuN (left) or negative for NeuN (right). **(F)** Scatter plot of average CTACIT Score (±SEM) by number of mCherry+ nuclei(±SEM) and trend line. EduAttan; GWAS for educational attainment; CigPerDay; GWAS for number of cigarettes smoked per day; PIP, fine mapping posterior inclusion probability; R, Pearson’s correlation; rho, Spearman’s correlation.

The phyloP conservation scores at these SNPs indicate that these nucleotides are conserved across placental mammals at the single-nucleotide level (candidate A phyloP_rs7933981_ = 7.0 and phyloP_rs61687445_ = 3.01; candidate B phyloP_rs7928017_ = 2.34) [11], and only the first two SNPs are conserved in primates using the PhastCons method, which accounts for conservation of neighboring nucleotides (**Figure 4A**, PhastCons_rs7933981_ = PhastCons_rs61687445_ = 1.0, PhastCon_rs7928017_ = 0) [10]. Multiple public functional annotations available through the UCSC genome browser indicate that candidate B has more regulatory activity than Candidate A (**Figure S29**). When statistical fine-mapping, genome conservation, and epigenomic annotations provide equivocal or conflicting evidence of SNP impact, we propose that CTACIT can shed light on cell type-specific regulatory function.

In contrast to existing functional evidence, we calculated a higher neuronal CTACIT Age for candidate A relative to B (**Figure 4C**). Both the human and mouse orthologs of candidate A have higher predicted neuronal activity than B (**Figure 4D**, **Data S11**, human sequence A, HSA, and mouse sequence A, MMA, vs. HSB and MMB). We compared our predictions with bulk ATAC-seq of the caudate nucleus in Egyptian fruit bats conducted by [63]. We measured multiple reproducible open chromatin regions at the fruit bat ortholog of candidate A yet none at the ortholog of candidate B. Even though CTACIT models were trained on data from two primate species and mouse, they correctly predicted conserved open chromatin activity at this locus in a bat species, 94 million years diverged from humans.

To test cell type-specific gene regulatory function of the two candidate regulatory elements upstream of the *DRD2* locus, we performed adeno-associated virus-based *in vivo* reporter assays of the two candidate enhancers (**Figure 1F, 4B, S30**). We co-stained transduced mouse brains for the reporter mCherry and the neuronal marker NeuN and found that both candidate sequences were able to express the reporter gene mCherry; namely, they act as functional enhancers (**Figure 4E, S31**). We detected more mCherry+ nuclei in injections of candidate A relative to candidate B (linear mixed effect t-test p_A vs. B_= 0.012, N=3 per reporter, **Data S12**). Whether we used the human or mouse ortholog of the enhancers did not significantly affect the detected level of neuronal mCherry+ nuclei (linear mixed effect t-test p_human vs mouse_ = 0.914, **Data S12**). We noticed that the predicted trends in neurons and glia correspond to the measured mCherry+ nuclei counts: CTACIT had predicted weak glial gene regulatory activation and stronger neuronal activation, and the correlation between CTACIT predictions and the mCherry+ nuclei measurements is strong (Pearson’s R = 0.794, p = 0.019, **Figure 4F**).

Our *in vivo* reporter experiments validate the CTACIT predictions of cell type-specific enhancer activity, not the GWAS trait associations themselves: they demonstrate that CTACIT quantitatively predicts the conserved gene regulatory activity of both human and mouse orthologs of a candidate sequence in the correct cell type-specific expression patterns. Both tested human sequences have varying degrees of cell type-specificity, so the fine-mapped SNPs within these enhancers may affect *DRD2* transcription in caudate cells at either or both enhancers. In addition to the *DRD2* locus, we highlight other loci with fine-mapped SNPs with predicted cell type-specific gene regulatory impacts (**Figure S32-34**, *Supplemental Text, Additional fine-mapped loci*).

## Discussion

The functional interpretation of non-coding variants remains a central problem in human genetics. Measurements such as nucleotide conservation metrics and open chromatin annotations each capture only one axis of the information that distinguishes a causal regulatory variant from a passenger due to linkage disequilibrium. We show that the conservation of cell type-specific regulatory function across mammals is a third axis, and that it adds prioritization power the other two miss. By integrating sequence conservation scores with cell type-resolved open chromatin from a few mammalian species, our machine learning tool CTACIT imputes regulatory activity across hundreds more species from the Zoonomia consortium and assigns each human OCR a measure of how long its cell type-specific function has persisted. Applied to a panel of neuropsychiatric traits, caudate cell type CTACIT models yields higher heritability enrichment and more fine-mapped variants than nucleotide conservation and human chromatin alone, and we validate its predictions for enhancers near the *DRD2* psychiatric disease risk locus with *in vivo* reporter assays. Cross-species functional conservation, measured at the resolution of individual cell types, can therefore serve as a prioritization layer for non-coding variant interpretation. We quantify the added prioritization these annotations provide rather than claim a formal statistical interaction between cell type specificity and conservation, which our current heritability models cannot separate (Limitations).

The case of schizophrenia, where deficits affect higher cognition and which is currently defined as human-specific [36,38,64], highlights the power of CTACIT to interpret the genetic basis of complex neuropsychiatric traits. Although some schizophrenia variants have nucleotide-level conservation, more tend to be found at loci where open chromatin is predicted to be highly conserved across placental mammals in specific caudate neuron types. Our findings support the biological model where regulatory elements containing trait-associated variants can tolerate turnover at individual nucleotide positions while maintaining regulatory activity in a particular cell type across distant species. This model reframes a long-standing puzzle: evolutionary annotations have been thought incapable of explaining schizophrenia heritability because the phenotype is human-specific [36–38], yet the regulatory elements that carry its risk variants can be deeply conserved in function even where individual nucleotides are not.

The accuracy of predictions across the placental phylogeny, in species up to roughly 100 million years diverged from the three used in training, may seem surprising, but it follows from how regulatory sequence is conserved. Some genomic elements retain cell type-specific gene regulation across Metazoa by shuffling and reforming critical transcription factor-binding sites [40], so the cell type-specific regulatory grammar a model learns is not tied to a particular nucleotide arrangement. Because CTACIT learns this grammar rather than memorizing exact sequences, a model trained on human, macaque, and mouse OCRs can recognize the same combinations of binding sites when they reappear in a rearranged form in a distant genome. We previously demonstrated that convolutional neural networks trained on a few species accurately identify orthologous sequences that retain tissue- and cell type-specific open chromatin activity across mammals [14,42] and the correct prediction of conserved activity at the DRD2 candidate enhancer in Egyptian fruit bat, 94 million years diverged from humans, shows the same generalization here. CTACIT joins a growing set of sequence-based models that predict regulatory activity and its conservation across species [41,42,65], but rather than predicting expression or accessibility within a single genome, it imputes cell type-resolved regulatory activity at orthologous loci across the placental phylogeny and tie that conservation directly to complex trait heritability.

CTACIT Age is one of several ways to summarize cross-species regulatory conservation, alongside simpler measures such as whether an OCR is mappable to a distant clade (e.g. fruit bats) or directly overlaps an OCR measured in another species (e.g. house mouse). These measures are complementary rather than competing. Mappability and pairwise OCR overlap capture conservation that is already supported by sequence alignment and direct measurement, and conditional effect sizes are larger for OCRs shared between human and another species than for OCRs from any one species alone (**Data S5**). CTACIT Age extends this logic to the full placental phylogeny by weighting predicted activity in each clade by its divergence time, reaching species where no chromatin data exist. A head-to-head benchmark establishing which summary best prioritizes causal variants is a natural next step that we leave to future work.

The opposing enrichment patterns of eQTLs and fine-mapped GWAS SNPs across CTACIT Age quartiles support the interpretation that the metric ranks functional variants. Caudate and matched cell type eQTLs were enriched in the “youngest” Age quartiles and depleted from “older” ones, while fine-mapped neuropsychiatric SNPs showed the reverse, with the highest enrichments in “older” medium spiny neuron and microglia quartiles (**Figure 3A-C**). Predicted “younger” regulatory regions have had less time under negative selection and so tolerate variants with variable effects on gene regulation[52], whereas the deeply conserved “older” regions are where trait-associated variation concentrates. That eQTLs and disease variants separate along the same axis in opposite directions [52,54] indicates CTACIT Age is capturing a real difference in selection pressure rather than an artifact of annotation construction.

## Limitations

The heritability enrichments we report are best read as relative rankings rather than absolute magnitudes. Smaller annotations tend to have larger enrichments, and the high values for predicted-active OCRs in distant clades partly reflect annotation size. Some of these annotations approach the size threshold at which S-LDSC heritability estimates become less reliable[66], so future analyses that size-match the CTACIT, PhyloP, and mappability annotations will sharpen the comparison of enrichment magnitudes (**Figure 2**, **Figure S11-26**).

Our OCR annotations are related to both nucleotide conservation and cell type specificity, and our heritability models combine these two properties. As a result, we cannot attribute the gain over existing models to either property alone, whether the improvement comes from OCR-level conservation capturing disease heritability beyond nucleotide conservation, from cell type-specific OCRs capturing cell type-specific heritability beyond other cell type-specific annotations, or from both. A heritability model that partitions per-SNP heritability into separate conservation, cell type-specificity, and OCR terms would isolate the independent contribution of the OCR annotations, and optimizing a single best-performing OCR annotation against models built from nucleotide conservation and cell type-differential accessibility would establish where these annotations add the most.

These analyses focus on common-frequency SNPs and extending them to low-frequency variants and to independent-trait sets would clarify how enrichments behave across genetically correlated traits. CTACIT Age offers room for similar refinement as a prioritization tool. Because older SNPs tend to have higher linkage disequilibrium, Age quartiles may be partly confounded by LD, and pairing CTACIT Age with fine-mapped eQTLs from methods such as PolyFun or SuSIE[53,67] would more directly resolve which variants drive the signal. A genome-wide benchmark of fine-mapping with and without the OCR annotations would further quantify how much they improve the number of fine-mapped loci and the size of their credible sets.

We made predictions across the placental mammalian phylogeny, but our models were trained using only primates and mouse, so the regulatory code they learned may not be conserved in all species where we made predictions. Confirming conservation beyond Euarchontoglires requires cell type regulatory genomics data from at least one mammal outside this clade. Finally, our *in vivo* reporter assays trade throughput for cell type resolution and measure each candidate enhancer in an episomal context, so future work pairing orthogonal CRISPR-knockout or CRISPR-inactivation with cell type targeting can extend these validations to more loci in the native genome[68,69]. Further limitations are noted in the *Supplemental Text, Limitations* section.

CTACIT enables future work using chromatin accessibility data from matched cell types in multiple species to prioritize human trait-associated variants for *in vivo* characterization and untangle their roles in complex tissues and traits. By integrating genome conservation and multi-species open chromatin, CTACIT prioritizes non-coding variants within regions of conserved regulatory function, predicts the cell type in which that function acts, and supports *in vivo* cell type-specific enhancer validation, as we demonstrate for enhancers near the *DRD2* locus. Future work could expand these *in vivo* experiments to identify interactions between human risk variants, gene regulatory mechanisms, and the specific neural circuits underlying behavior in rodents, primates, or non-traditional model organisms. Some experiments can be paired with established paradigms such as intravenous drug self-administration for substance use behaviors or multi-arm bandit gambling tasks for reward and risk-taking behaviors, dissecting the gene-to-behavior relationships ultimately key to understanding the human genetics of complex traits.

## Acknowledgments

We thank members of the Pfenning lab, Diane P. Genereux, Ruby A. Holland, Michael Leone, Patrick F. Sullivan, Samuel J. Dienel, and members of the Zoonomia Consortium for discussions and feedback. We thank Jacquelyn Breter for taking excellent care of animals. We thank Martin Jinye Zhang for useful comments on the manuscript. One or more of the authors of this paper self-identifies as a member of the LGBTQ+ community.

## Funding

National Institutes of Health NIDA grant DP1DA046585 (ARP)

National Institutes of Health NIDA grant F30DA053020 (BNP)

National Institutes of Health NIMH grant UG3MH120094 (WRS)

Carnegie Mellon University Computational Biology Department Lane Fellowship (IMK)

Carnegie Mellon University Neuroscience Institute (AK)

## Author contributions

Conceptualization, Writing – original draft: BNP, ARP Methodology: BNP

Visualization: BNP, AK, AJL Software: BNP, DES

Investigation: BNP, JH, ARB, GAF, CS, IMK, MEW Formal Analysis: BNP, AJL, AK

Resources: ARP, AJL, ARB, JH, WRS

Data Curation: BNP, AJL, ARB, IMK, MEW Writing – review & editing: all authors Supervision: ARP

Funding acquisition: ARP, WRS

## Data and materials availability

Code for analyses and cross-species convolutional neural network models of caudate cell types used in this study are available at https://github.com/pfenninglab/Zoonomia_CTACIT_fine-mapping. Open chromatin regions from human, macaque, and mouse are uploaded within the github repository. Rhesus macaque snATAC-seq sequencing reads are deposited to GEO. Other intermediate files will be available by request to the corresponding author.

## Supplementary Materials

Materials and Methods

Supplementary Text

Figures S1 to S34

References (59–158)

Data S1 to S12

## References

1. Sullivan PF, Agrawal A, Bulik CM, Andreassen OA, Børglum AD, Breen G, et al. Psychiatric Genomics: An Update and an Agenda. Am J Psychiatry. 2018;175:15.

2. Sullivan PF, Meadows JRS, Gazal S, Phan BN, Li X, Genereux DP, et al. Leveraging base-pair mammalian constraint to understand genetic variation and human disease. Science. 2023;380:eabn2937.

3. Consortium RE, Kundaje A, Meuleman W, Ernst J, Bilenky M, Yen A, et al. Integrative analysis of 111 reference human epigenomes. Nature. 2015;518:317–30.

4. Meuleman W, Muratov A, Rynes E, Halow J, Lee K, Bates D, et al. Index and biological spectrum of human DNase I hypersensitive sites. Nature. 2020;584:244–51.

5. Finucane HK, Bulik-Sullivan B, Gusev A, Trynka G, Reshef Y, Loh P-R, et al. Partitioning heritability by functional annotation using genome-wide association summary statistics. Nat Genet. 2015;47:1228.

6. Lindblad-Toh K, Garber M, Zuk O, Lin MF, Parker BJ, Washietl S, et al. A high-resolution map of human evolutionary constraint using 29 mammals. Nature. 2011;478:476–82.

7. Gazal S, Finucane HK, Furlotte NA, Loh P-R, Palamara PF, Liu X, et al. Linkage disequilibrium dependent architecture of human complex traits reveals action of negative selection [Internet]. Available from: 10.1101/082024

8. Gazal S, Loh P-R, Finucane HK, Ganna A, Schoech A, Sunyaev S, et al. Functional architecture of low-frequency variants highlights strength of negative selection across coding and non-coding annotations. Nat Genet. 2018;50:1600–7.

9. Davydov EV, Goode DL, Sirota M, Cooper GM, Sidow A, Batzoglou S. Identifying a high fraction of the human genome to be under selective constraint using GERP++. PLoS Comput Biol. 2010;6:e1001025.

10. Siepel A, Bejerano G, Pedersen JS, Hinrichs AS, Hou M, Rosenbloom K, et al. Evolutionarily conserved elements in vertebrate, insect, worm, and yeast genomes. Genome Res. 2005;15:1034–50.

11. Pollard KS, Hubisz MJ, Rosenbloom KR, Siepel A. Detection of nonneutral substitution rates on mammalian phylogenies. Genome Res [Internet]. 2010 [cited 2021 Mar 11];20. Available from: https://pubmed.ncbi.nlm.nih.gov/19858363/

12. Burch KS, Hou K, Ding Y, Wang Y, Gazal S, Shi H, et al. Partitioning gene-level contributions to complex-trait heritability by allele frequency identifies disease-relevant genes [Internet]. Available from: 10.1101/2021.08.17.456722

13. Hujoel MLA, Gazal S, Hormozdiari F, van de Geijn B, Price AL. Disease heritability enrichment of regulatory elements is concentrated in elements with ancient sequence age and conserved function across species [Internet]. Available from: 10.1101/420166

14. Kaplow IM, Schäffer DE, Wirthlin ME, Lawler AJ, Brown AR, Kleyman M, et al. Inferring mammalian tissue-specific regulatory conservation by predicting tissue-specific differences in open chromatin. BMC Genomics [Internet]. 2022 [cited 2022 July 5];23. Available from: https://pubmed.ncbi.nlm.nih.gov/35410163/

15. Li YE, Preissl S, Hou X, Zhang Z, Zhang K, Qiu Y, et al. An atlas of gene regulatory elements in adult mouse cerebrum. Nature. 2021;598:129–36.

16. Srinivasan C, Phan BN, Lawler AJ, Ramamurthy E, Kleyman M, Brown AR, et al. Addiction-associated genetic variants implicate brain cell type- and region-specific cis-regulatory elements in addiction neurobiology. J Neurosci [Internet]. 2021; Available from: 10.1523/JNEUROSCI.2534-20.2021

17. Pan Z, Yao Y, Yin H, Cai Z, Wang Y, Bai L, et al. Pig genome functional annotation enhances the biological interpretation of complex traits and human disease. Nat Commun. 2021;12:5848.

18. Cusanovich DA, Hill AJ, Aghamirzaie D, Daza RM, Pliner HA, Berletch JB, et al. A Single-Cell Atlas of In Vivo Mammalian Chromatin Accessibility. Cell. 2018;174:1309–1324.e18.

19. Corces MR, Shcherbina A, Kundu S, Gloudemans MJ, Frésard L, Granja JM, et al. Single-cell epigenomic analyses implicate candidate causal variants at inherited risk loci for Alzheimer’s and Parkinson’s diseases. Nat Genet. 2020;52:1158–68.

20. Dey KK, van de Geijn B, Kim SS, Hormozdiari F, Kelley DR, Price AL. Evaluating the informativeness of deep learning annotations for human complex diseases. Nat Commun. 2020;11:4703.

21. van de Geijn B, Finucane H, Gazal S, Hormozdiari F, Amariuta T, Liu X, et al. Annotations capturing cell type-specific TF binding explain a large fraction of disease heritability. Hum Mol Genet. 2020;29:1057–67.

22. Amariuta T, Ishigaki K, Sugishita H, Ohta T, Koido M, Dey KK, et al. Improving the trans-ancestry portability of polygenic risk scores by prioritizing variants in predicted cell-type-specific regulatory elements. Nat Genet. 2020;52:1346–54.

23. Hook PW, McCallion AS. Leveraging mouse chromatin data for heritability enrichment informs common disease architecture and reveals cortical layer contributions to schizophrenia. Genome Res. 2020;30:528–39.

24. Christmas MJ, Kaplow IM, Genereux DP, Dong MX, Hughes GM, Li X, et al. Evolutionary constraint and innovation across hundreds of placental mammals. Science. 2023;380:eabn3943.

25. Grillner S, Robertson B, Stephenson-Jones M. The evolutionary origin of the vertebrate basal ganglia and its role in action selection. J Physiol. 2013;591:5425–31.

26. Fullard JF, Hauberg ME, Bendl J, Egervari G, Cirnaru M-D, Reach SM, et al. An atlas of chromatin accessibility in the adult human brain. Genome Res. 2018;28:1243–52.

27. Skene NG, Bryois J, Bakken TE, Breen G, Crowley JJ, Gaspar HA, et al. Genetic identification of brain cell types underlying schizophrenia. Nat Genet. 2018;50:825–33.

28. Bryois J, Skene NG, Hansen TF, Kogelman LJA, Watson HJ, Liu Z, et al. Genetic identification of cell types underlying brain complex traits yields insights into the etiology of Parkinson’s disease. Nat Genet. 2020;52:482–93.

29. Jansen PR, Watanabe K, Stringer S, Skene N, Bryois J, Hammerschlag AR, et al. Genome-wide analysis of insomnia in 1,331,010 individuals identifies new risk loci and functional pathways. Nat Genet. 2019;51:394–403.

30. Grillner S, Robertson B. The Basal Ganglia Over 500 Million Years. Curr Biol. 2016;26:R1088–100.

31. Gokce O, Stanley GM, Treutlein B, Neff NF, Camp JG, Malenka RC, et al. Cellular Taxonomy of the Mouse Striatum as Revealed by Single-Cell RNA-Seq. Cell Rep. 2016;16:1126–37.

32. Stanley G, Gokce O, Malenka RC, Südhof TC, Quake SR. Continuous and Discrete Neuron Types of the Adult Murine Striatum. Neuron. 2020;105:688–699.e8.

33. He J, Kleyman M, Chen J, Alikaya A, Rothenhoefer KM, Ozturk BE, et al. Transcriptional and anatomical diversity of medium spiny neurons in the primate striatum. Curr Biol [Internet]. 2021; Available from: 10.1016/j.cub.2021.10.015

34. Buyukkahraman G, Caglayan E, Hörpel SG, Zhang Y, van Tussenbroek IA, Orozco CG, et al. Comparative analysis of the cellular landscape in mammalian striatum. Nat Commun [Internet]. 2026; Available from: 10.1038/s41467-026-73305-8

35. Huo Y, Li S, Liu J, Li X, Luo X-J. Functional genomics reveal gene regulatory mechanisms underlying schizophrenia risk. Nat Commun. 2019;10:670.

36. Bhattacharyya U, Bhatia T, Deshpande SN, Thelma BK. Genetic variations in evolutionary accelerated regions disrupt cognition in schizophrenia. Psychiatry Res. 2022;314:114586.

37. Srinivasan S, Bettella F, Hassani S, Wang Y, Witoelar A, Schork AJ, et al. Probing the Association between Early Evolutionary Markers and Schizophrenia. PLoS One. 2017;12:e0169227.

38. Banerjee N, Polushina T, Bettella F, Giddaluru S, Steen VM, Andreassen OA, et al. Recently evolved human-specific methylated regions are enriched in schizophrenia signals. BMC Evol Biol. 2018;18:63.

39. Consortium TSWG of TPG, The Schizophrenia Working Group of the Psychiatric Genomics Consortium, Ripke S, Walters JTR, O’Donovan MC. Mapping genomic loci prioritises genes and implicates synaptic biology in schizophrenia. Nature [Internet]. 2022; Available from: 10.1038/s41586-022-04434-5

40. Wong ES, Zheng D, Tan SZ, Bower NL, Garside V, Vanwalleghem G, et al. Deep conservation of the enhancer regulatory code in animals. Science [Internet]. 2020;370. Available from: 10.1126/science.aax8137

41. Chen L, Fish AE, Capra JA. Prediction of gene regulatory enhancers across species reveals evolutionarily conserved sequence properties. PLoS Comput Biol. 2018;14:e1006484.

42. Kaplow IM, Lawler AJ, Schäffer DE, Srinivasan C, Sestili HH, Wirthlin ME, et al. Relating enhancer genetic variation across mammals to complex phenotypes using machine learning. Science. 2023;Apr 28;380:eabm7993.

43. Genereux DP, Seres A, Armstrong J, Johnson J, Marinescu VD, Murén E, et al. A comparative genomics multitool for scientific discovery and conservation. Nature. 2020;587:240–5.

44. Armstrong J, Hickey G, Diekhans M, Fiddes IT, Novak AM, Deran A, et al. Progressive Cactus is a multiple-genome aligner for the thousand-genome era. Nature. 2020;587:246–51.

45. Kunkle BW, Grenier-Boley B, Sims R, Bis JC, Damotte V, Naj AC, et al. Genetic meta-analysis of diagnosed Alzheimer’s disease identifies new risk loci and implicates Aβ, tau, immunity and lipid processing. Nat Genet. 2019;51:414–30.

46. Jansen IE, Savage JE, Watanabe K, Bryois J, Williams DM, Steinberg S, et al. Genome-wide meta-analysis identifies new loci and functional pathways influencing Alzheimer’s disease risk. Nat Genet. 2019;51:404–13.

47. Liu M, Jiang Y, Wedow R, Li Y, Brazel DM, Chen F, et al. Association studies of up to 1.2 million individuals yield new insights into the genetic etiology of tobacco and alcohol use. Nat Genet. 2019;51:237–44.

48. Consortium TG, The GTEx Consortium. The GTEx Consortium atlas of genetic regulatory effects across human tissues [Internet]. Science. 2020. p. 1318–30. Available from: 10.1126/science.aaz1776

49. Bryois J, Calini D, Macnair W, Foo L, Urich E, Ortmann W, et al. Cell-type-specific cis-eQTLs in eight human brain cell types identify novel risk genes for psychiatric and neurological disorders. Nat Neurosci. 2022;25:1104–12.

50. Herrero-Navarro Á, Puche-Aroca L, Moreno-Juan V, Sempere-Ferràndez A, Espinosa A, Susín R, et al. Astrocytes and neurons share region-specific transcriptional signatures that confer regional identity to neuronal reprogramming [Internet]. Science Advances. 2021. Available from: 10.1126/sciadv.abe8978

51. Chai H, Diaz-Castro B, Shigetomi E, Monte E, Octeau JC, Yu X, et al. Neural Circuit-Specialized Astrocytes: Transcriptomic, Proteomic, Morphological, and Functional Evidence. Neuron. 2017;95:531–549.e9.

52. Mostafavi H, Spence JP, Naqvi S, Pritchard JK. Systematic differences in discovery of genetic effects on gene expression and complex traits. Nat Genet. 2023;55:1866–75.

53. Weissbrod O, Hormozdiari F, Benner C, Cui R, Ulirsch J, Gazal S, et al. Functionally informed fine-mapping and polygenic localization of complex trait heritability. Nat Genet. 2020;52:1355–63.

54. Glassberg EC, Gao Z, Harpak A, Lan X, Pritchard JK. Evidence for Weak Selective Constraint on Human Gene Expression. Genetics. 2019;211:757–72.

55. Lee JJ, Wedow R, Okbay A, Kong E, Maghzian O, Zacher M, et al. Gene discovery and polygenic prediction from a genome-wide association study of educational attainment in 1.1 million individuals. Nat Genet. 2018;50:1112–21.

56. Nagel M, Jansen PR, Stringer S, Watanabe K, de Leeuw CA, Bryois J, et al. Meta-analysis of genome-wide association studies for neuroticism in 449,484 individuals identifies novel genetic loci and pathways. Nat Genet. 2018;50:920–7.

57. Benjamin KJM, Chen Q, Jaffe AE, Stolz JM, Collado-Torres L, Huuki-Myers LA, et al. Analysis of the caudate nucleus transcriptome in individuals with schizophrenia highlights effects of antipsychotics and new risk genes. Nat Neurosci. 2022;25:1559–68.

58. Zhou H, Kember RL, Deak JD, Xu H, Toikumo S, Yuan K, et al. Multi-ancestry study of the genetics of problematic alcohol use in over 1 million individuals. Nat Med. 2023;29:3184–92.

59. Hatoum AS, Colbert SMC, Johnson EC, Huggett SB, Deak JD, Pathak GA, et al. Multivariate genome-wide association meta-analysis of over 1 million subjects identifies loci underlying multiple substance use disorders. Nat Ment Health. 2023;1:210–23.

60. Major Depressive Disorder Working Group of the Psychiatric Genomics Consortium. Electronic address, Major Depressive Disorder Working Group of the Psychiatric Genomics Consortium. Trans-ancestry genome-wide study of depression identifies 697 associations implicating cell types and pharmacotherapies. Cell. 2025;188:640–652.e9.

61. Hirvonen J, van Erp TGM, Huttunen J, Aalto S, Någren K, Huttunen M, et al. Increased caudate dopamine D2 receptor availability as a genetic marker for schizophrenia. Arch Gen Psychiatry. 2005;62:371–8.

62. Abi-Dargham A, Rodenhiser J, Printz D, Zea-Ponce Y, Gil R, Kegeles LS, et al. Increased baseline occupancy of D2 receptors by dopamine in schizophrenia. Proc Natl Acad Sci U S A. 2000;97:8104–9.

63. Wirthlin ME, Schmid TA, Elie JE, Zhang X, Kowalczyk A, Redlich R, et al. Vocal learning-associated convergent evolution in mammalian proteins and regulatory elements. Science. 2024;383:eabn3263.

64. Mathalon DH, Ford JM. Neurobiology of schizophrenia: search for the elusive correlation with symptoms. Front Hum Neurosci. 2012;6:136.

65. Avsec Ž, Agarwal V, Visentin D, Ledsam JR, Grabska-Barwinska A, Taylor KR, et al. Effective gene expression prediction from sequence by integrating long-range interactions. Nat Methods. 2021;18:1196–203.

66. Tashman K, Cui R, O’Connor L, Neale B, Finucane H. Significance testing for small annotations in stratified LD-Score regression [Internet]. medRxiv. medRxiv; 2021. Available from: 10.1101/2021.03.13.21249938

67. Wang G, Sarkar A, Carbonetto P, Stephens M. A simple new approach to variable selection in regression, with application to genetic fine mapping. J R Stat Soc Series B Stat Methodol. 2020;82:1273–300.

68. Gasperini M, Hill AJ, McFaline-Figueroa JL, Martin B, Kim S, Zhang MD, et al. A Genome-wide Framework for Mapping Gene Regulation via Cellular Genetic Screens. Cell. 2019;176:377–390.e19.

69. Klann TS, Black JB, Chellappan M, Safi A, Song L, Hilton IB, et al. CRISPR-Cas9 epigenome editing enables high-throughput screening for functional regulatory elements in the human genome. Nat Biotechnol. 2017;35:561–8.

