## Supplemental Text for "Fine-mapping candidate neuropsychiatric regulatory variants using cell type-aware comparative genomics"

**Affiliations:**

^4^Current Affiliation: Broad Institute of Harvard and MIT; Cambridge, MA, USA.

^7^Current Affiliation: Allen Institute; Seattle, WA, USA.

**This PDF file includes:**

Materials and Methods

Supplementary Text

Figures S1 to S34

References (59–158)

Data S1 to S12

Materials & Methods

### Animal Care & Use

All non-human primate animal procedures were in accordance with the National Institutes of Health Guide for the Care and Use of Laboratory Animals and approved by the University of Pittsburgh’s Institutional Animal Care and Use Committee (IACUC) (Protocol ID, 19024431). Rhesus macaque monkeys were single- or pair-housed with a 12h-12h light-dark cycle. Monkey P was a 5-year-old female (5.4kg).

All mouse animal procedures were in accordance with the National Institutes of Health Guide for the Care and Use of Laboratory Animals and approved by Carnegie Mellon University’s IACUC (Protocol ID, 201600003). The mice used for reporter assay experiments were C57BL/6J wild type mice or D1-*cre* and A2a-*cre* mice maintained on a C57BL/6J background from the GENSAT Consortium (Jackson Laboratory, Bar Harbor, ME). All mice were 2-5 months old at the time of the sacrifice. Mice experiments included representation from both sexes. All animals were housed with a 12 hour light cycle, and experiments were performed at the same time of day relative to lights on. Animals for the data primary to this study received no treatments other than the retro-orbital AAV injections.

### Non-Human Primate Single Nuclei ATAC-seq

To maximize nuclei viability for single nuclei ATAC-seq (below), we followed a harvesting protocol described in [1]. Briefly, we sedated Monkey P with ketamine and maintained general anesthesia with isoflurane. We transported the monkey to a surgery suite and placed it in a stereotaxic frame (Kopf Instruments). We removed the calvarium and then perfused the monkey with 3-4 liters of ice cold artificial cerebrospinal fluid (ACSF; 124 mM NaCl, 5 mM KCl, 2 mM MgSO4, 2 mM CaCl2, 23 mM NaHCO3, 3 mM NaH2PO4, 10 mM glucose; pH 7.4, osmolarity 290–300 mOsm) oxygenated with 95 % O2:5 % CO2. We then opened the dura and removed the brain. We sliced the brain into slabs on a custom 3-D printed matrix along the rostral-caudal axis. We dissected the caudate nucleus under a dissection microscope.

We isolated nuclei according to the 10X Genomics “Demonstrated Protocol: Nuclei Isolation for Single Cell ATAC Sequencing”. Briefly, we cut brain tissue into small pieces and triturated the tissue in NbActiv1 buffer (BrainBits) using wide- and regular-bore pipette tips sequentially to dissociate cells. We filtered the cells through a 30 um MACS SmartStrainer (130-098-458) into an Eppendorf tube. After centrifuging, we used a lysis buffer (1mM Tris-HCl, 1mM NaCl, 0.3 mM MgCl_2_, 0.01% Tween-20, 0.01% Nonidet P40 Substitute, 0.001%. Digitonin, 0.1% BSA) to lyse the cells for 5 min. We washed the lysed cells once with 1 ml chilled wash buffer (1 mM Tris-HCl, 1mM NaCl, 0.3 mM MgCl_2_, 0.1% BSA, 0.01% Tween-20) and resuspended the nuclei pellet with Diluted Nuclei Buffer (10X Genomics, PN-2000153). We filtered the nuclei through a strainer and counted the nuclei.

We used 10x Chromium Single Cell ATAC Library & Gel Bead Kit (10X Genomics, Cat# PN-1000110) for monkey P. We transposed the nuclei and generated libraries according to 10x Genomics protocol and briefly incubated the nuclei suspensions in a transposition mix that includes transposase, which preferentially fragments the DNA in open chromatin. We generated Gel beads-in-emulsion (GEMs) after running through a 10x Genomics Chromium controller. We barcoded open chromatin within GEMs in a Bio-Rad PCR machine (Cat# C1000). We purified DNA with Dynabeads (10X Genomics, PN-2000048) and SPRIselect reagent (Beckman Coulter, Cat# B23318) after breaking the emulsion with a recovery agent (10X Genomics, 220016). Then, we amplified and indexed the samples by sample index PCR and purified them with SPRIselect reagent. We analyzed quality of the libraries using Agilent Bioanalyzer 2100 and quantified the libraries by qPCR using a KAPA Library Quantification Kit (KAPA Biosystems, Cat# KK4824). We pooled together libraries and loaded them onto NovaSeq 6000 S4 Flow Cell. We sequenced N=2 technical replicates of the caudate nucleus from monkey P to the depth of 150,000 x 150 bp paired-end reads per nuclei (Figure S1A, B).

### Rhesus Macaque snATAC-seq read processing

We de-multiplexed sequencing reads with dex-fastq from the SnapTools package (<https://github.com/r3fang/SnapTools>) [2], trimmed them for the transposase adapter sequence “CTGTCTCTTATACACATCT” on the 3’ end of reads with cutadapt [3], and aligned them to the rheMac10 genome with bowtie2 with the “--sensitive --no-discordant --no-mixed --minins 25 --maxins 1000” parameters to map fragments between 25bp and 1000bp long [4,5]. After aligning the reads, we removed PCR duplicates with samtools [6] and created arrow files for downstream analyses with the ArchR package [7]. We followed the approach by Jing, Kleyman *et al*. to map the higher quality GRCh38.p13 human RefSeq gene annotations onto the rheMac10 genome with the liftOff tool [8,9] and used this to create a custom rheMac10 ArchR gene and genome annotation as outlined within the ArchR manual. Using this liftOff annotation, we were able to compute ArchR gene activity scores around the orthologous rhesus macaque regions of human genes, enabling us to obtain scores for substantially more genes and complete gene bodies than we would have been able to obtain using base rheMac10 gene annotations. Within ArchR, we removed inferred doublet nuclei from each sample with cutEnrich = .5 and filterRatio = 1. Then we performed iterative latent-semantic index (LSI) clustering with 4 iterations and 30,000-150,000 variable features followed by Harmony batch correction [10], UMAP visualization, and Louvain clustering, where we used default settings for all methods. We identified snATAC-seq clusters by co-clustering with Seurat v3’s canonical correlation analysis (CCA) on ArchR gene activity scores [7,11] with previously published single nucleus RNA-seq gene expression of the rhesus macaque striatum [9]. We called peaks across technical replicates with macs2 [12], identified reproducible peaks between replicates for each cell type, created a consensus peakset across cell types, and created a peak by cell “PeakMatrix” through ArchR [7].

### Reporter Assay Molecular Cloning

We created the template *in vivo* reporter plasmid construct, pAAV-Hsp68-pNLS/mCh-3rNLS/ReporterAssayEmpty (pAAV-ReporterAssayEmpty), with VectorBuilder Cloning Services (VectorBuilder Inc, Chicago, IL). The template plasmid contains inverted terminal repeats (ITRs) standard for AAV production, Hsp68 minimal promoter, synthetic intron [13], mCherry with two SV40 Large T antigen nuclear localization signals (PKKKRKVED) on the N- and C- terminals [14], and the Chloramphenicol resistance gene as a place-holder for the candidate enhancer between two pairs of multiple cloning sites. After receiving the plasmid, we also added three RNA nuclear localization motifs (AGCCC) outside the coding sequence and before the SV40 poly-adenylation signal [15] (**Figure S33**).

To clone in the candidate enhancers, we synthesized the human enhancer sequence by IDT as gBlocks™ with common 5’ (GAGTCTGAACCTGTGTGCTA) and 3’ (TCAGAAGATCGGAAGAGCATCGTAG) sequences for downstream PCR amplification (Integrated DNA Technologies). We used PCR to amplify synthesized inserts using the KAPA HiFi HotStart ReadyMix protocol (Roche, KK2601) with primers to add BsmBI restriction enzyme sites compatible with Golden Gate cloning (F: GGCTACCGTCTCACATGGAGTCTGAACCTGTGTGCTA, R: GGCTACCGTCTCGACCACTACGATGCTCTTCCGATCT). We linearized the pAAV-ReporterAssayEmpty plasmid using the Q5 HotStart HiFi ReadyMix protocol (NEB, M0494) with primers to both add BsmBI restriction enzyme sites and excise the chloramphenicol place-holder (F: GGCTACCGTCTCGTGGTACCTAGCATCTCAC, R: GGCTACCGTCTCCCATGTTTCTCCATTTGGCGC). We cloned the enhancers into the pAAV-RAe plasmids with the NEB Golden Gate Assembly Kit (NEB, E1602) following manufacturer instructions. We validated cloning accuracy with Sanger sequencing.

### AAV production and delivery

We packaged reporter assay constructs into individual AAV preps packaged with the AAV-PhP.eB capsid that crosses the blood-brain barrier and has high transduction of neural cell types as previously described [16,17]. Briefly, AAV was produced in AAVpro(R) 293T cells (Takara, Kyoto, Japan; #632273) by co-transfection of the genome pAAV Reporter Assay plasmid, an AAV helper plasmid, and pUCmini-iCAP-PHP.eB. pUCmini-iCAP-PHP.eB was a gift from Viviana Gradinaru (<http://n2t.net/addgene:103005>; RRID: Addgene 103005) [18]. We precipitated the AAV particles with Polyethylene Glycol (PEG 8000, Sigma-Aldrich, St. Louis, MO; cat. P2139-500G) from the cell media, lysed the cells with Salt Active Nuclease enzyme (SAN, ArcticZymes, Sykehusveien, Norway, cat. 70900-201), and purified AAV on an iodixanol gradient (OptiPrep, Sigma-Aldrich, cat. D1556-250ML) with ultracentrifugation for 2.5 hours at 350,000 x g at 18°C. We filtered and concentrated the virus in PBS+F-68 (PBS, Pluronic(™) F-68 Non-ionic Surfactant (100x) 0.001%) using Amicon Ultra-15 centrifugation filters (Millipore, Burlington, MA; #UFC905024) and resuspended in PBS. We measured the viral titer for each with the AAVpro(R) Titration Kit (Takara, #6233), diluted to the same concentration of 2.32 x 10^9^ vector genomes (vg) / µL, and stored in single-use aliquots at −80 °C until injection.

We used three cages of 4 adult age- and sex-matched littermates for each retro-orbital virus injection. We anesthetized each animal with 2-3% isoflurane until we did not observe any pedal withdrawal reflex and maintained anesthesia with 1-2% isoflurane during the procedure. Then, we injected each reporter assay AAV (1.16 x 10^11^ vg in 50 µL) with a viral transduction control AAV expressing nuclear-localizing GFP (4 x 10^11^ vg in 50 µL) of into the retro-orbital cavity and treated the eye with 0.5% Proparacaine Hydrochloride Ophthalmic Solution. We monitored the animals while the virus incubated for 2 weeks before brains were harvested.

### Immunofluorescence and Microscopy

We fixed animals with 4% paraformaldehyde (PFA) transcardial perfusion and incubated the brains in 4% PFA overnight after dissection. We cut 50-60 µm thick coronal slices with a vibrating microtome (Leica VT1000 S). We performed immuno-fluorescence staining of transduced mouse brains to assay for reporter gene presence in caudoputamen neurons, which overwhelmingly are canonical D1 or D2 MSNs in equal proportions [19] and are where the rodent caudoputamen is the orthologous region to primate caudate nucleus. We co-stained free-floating sections with anti-mCherry primary antibody (Cell Signaling; E5D8F) to amplify signal with AlexaFluor 594 secondary antibody (Cell Signaling Technology, Danvers, MA; #8889) as well as anti-NeuN primary antibody (ABcam; ab279297) to stain for neurons with AlexaFluor 405 (Thermo Fisher; A48268). We took images of the caudoputamen blinded to experimental conditions with a brightfield epi-fluorescence microscope under 20x objective (Nikon, Eclipse Ti2) with two to three 3x3 tiled images per animal capturing different coronal views of the dorsal striatum in both the left or right hemispheres. Each image had 0.365 micron per pixel in the x- and y- directions, and we reduced background noise using the NIS-Elements denoising software.

### Human caudate snATAC-seq preprocessing

We downloaded human snATAC-seq reads aligned to the hg38 genome from GSE147672 [20]. We created a human caudate snATAC-seq ArchR project (N=3 biological replicates) and assigned glial and neuronal subtype labels from the publication’s table SupplementaryDataSet2_scATAC-QC-And-Cluster-Residence_v1.xlsx. We subclustered the main MSN cluster to resolve D1 and D2 MSN subtypes by increasing the number of variable features during iterative LSI and otherwise used the Corces *et al.* [20] caudate cell type labels: D1 MSN, D2 MSN, striatonigral (SN) MSN, *PVALB*+ interneurons, astrocytes, oligodendrocytes, oligodendrocyte precursors (OPC), and microglia.

### Mouse caudoputamen cell types reprocessing

We downloaded the de-multiplexed sequencing reads for the caudoputamen (regions 4D and 5E) samples of mouse snATAC-seq from the NeMO archive (N=4 biological replicates, <https://nemoarchive.org>) [21]. We aligned de-multiplexed reads with the same method that we used for rhesus macaque, aligned reads with bowtie2 to the mm10 mouse genome, and created arrow files and ArchR projects for the mouse. We assigned the neuronal and glial subtypes from the cell barcodes available through the CATlas CellBrowser webportal (<http://catlas.org>). For D1 MSNs, D2 MSNs, and interneurons, most of which had high *PVALB* gene activity scores in primate snATAC-seq, we instead used the corresponding, previously published high-quality cre-dependent Sun1GFP nuclear-anchored independent labeled (cSNAIL) ATAC-seq datasets from mouse caudoputamen, which we downloaded from Srinivasan, Phan *et al.* GSE161374 [16].

### Bulk M1 cortex and liver ATAC-seq data processing and peak ortholog mapping

We compared the above cell type ATAC-seq profiles with cross-species matched bulk ATAC-seq profiles. For these data, we used the processed house mouse motor cortex ATAC-seq data from [16], the processed rhesus macaque and Norway rat motor cortex ATAC-seq data from [22] that were originally presented in [23], and the processed Egyptian fruit bat motor cortex ATAC-seq data from GSE187366 (access token: exsheqcazjsnxsx) [24,25]. For rhesus macaque and Egyptian fruit bat, we used the subset of orofacial motor cortex peaks that overlapped hand/wing motor cortex peaks, as described in the [24]. We used the processed liver ATAC-seq data from [22], where the house mouse data was originally presented in [22], the rhesus macaque and Norway rat data that were originally presented in [23], and the cow and pig data that were originally presented in [26]. For these datasets, we used the open chromatin regions that were filtered to be likely enhancers in [22] and [24]. We used the orthologs in the Zoonomia genomes obtained using the Zoonomia v2 CACTUS [27] that were obtained in [24].

### Cross-species peak ortholog mapping

For all snATAC-seq and cSNAIL-ATAC-seq cell type peaks from the primate caudate nucleus and mouse caudoputamen, we kept peaks in the narrowPeak format to retain the summit information for each peak, as critical transcription factor motifs have been shown to be enriched near open chromatin peak summits [28]. For ArchR snATAC-peaks, we achieved this by taking the summits of reproducible peaks in each cell type and extending them 250bp in both directions. We named each peak uniquely with “refGenome:chr:start:stop:summit” to track it and its unique, usable orthologs when mapping across species. We mapped peaks across Zoonomia genomes using the Zoonomia version 2 CACTUS alignment [29] (<https://cglgenomics.ucsc.edu/data/cactus/>) with halLiftover [30] and identified unique, usable orthologs with HALPER’s orthologFind.py script with parameters -max_frac 1.5 -min_len 50 -protect_dist 10 -narrowPeak (<https://github.com/pfenninglab/halLiftover-postprocessing>) [28]. We mapped human snATAC-seq peaks for 8 cell types across the 240 genomes in the Zoonomia alignment. We mapped the house mouse and rhesus macaque snATAC-seq and cSNAIL ATAC-seq peaks to the human genome. Prior to applying halLiftover and HALPER for the rhesus macaque peaks, we used the liftOver tool [31] to map rheMac10 coordinates to coordinates in rheMac8, which is the genome in the 241 Zoonomia alignment. We mapped both the peaks (chr:start-stop) and the summits (chr:summit), keeping only peaks where a summit was mappable within its corresponding peak.

To evaluate whether the regulatory codes in the identified caudate cell types were sufficiently conserved across between primates and mouse to train CTACIT models that could achieve high lineage-specific OCR accuracy, we compared the enrichments of TF motifs occurring in mouse OCRs to those enriched for occurring in human OCRs, as in [24]. We then ran MEME-ChIP (*131*) on the OCRs identified in each cell type from each respective species, and we used the respective -db to the CIS-BP version 2.00 databases (*86*) for *Homo sapiens*, *Macaca mulatta*, and *Mus musculus*. We used default settings along with the parameters -spamo-skip, and -fimo-skip. After that, we compared the motif enrichments of each cell type across species.

### LD score regression in hg38 coordinates

We performed linkage disequilibrium score regression (LDSC) analyses within the hg38 coordinates by applying liftOver across the published bed files baseline annotation (v1.1) and baselineLD annotations (v2.2) for cell type-specific LDSC and partitioned heritability enrichment LDSC analyses, respectively, as recommended by original authors (<https://alkesgroup.broadinstitute.org/LDSCORE/>, <https://github.com/bulik/ldsc/wiki/LD-Score-Estimation-Tutorial>) [32,33]. We used the 1000G SNPs mapped to hg38 coordinates across super-populations (ftp://[ftp.1000genomes.ebi.ac.uk/vol1/ftp/release/20130502/supporting/GRCh38_positions](http://ftp.1000genomes.ebi.ac.uk/vol1/ftp/release/20130502/supporting/GRCh38_positions)) [34] and computed LD scores as previously outlined using the --ld-wind-kb 1000 parameter. Using these LD scores, we computed genetic correlations between traits [35], LD scores for additional annotations in hg38 coordinates, and partitioned heritability analyses as previously described [32,33].

For LDSC cell type-specific analyses (<https://github.com/bulik/ldsc/wiki/Cell-type-specific-analyses>), we sought to estimate the independent contribution of an annotation after regressing out a background annotation and the baseline (v1.1) annotations. This coefficient is further standardized to the number of SNPs in the annotation and the per-SNP estimated narrow-sense heritability, the *τ** as described by [32,36]. This allowed us to compare both the significant contribution of a cell type annotation to a trait as well as the relative effect sizes between traits with varying power and heritability. We compared *τ** in human cell type OCRs to cell type OCRs from other species across significant LDSC traits with a standard linear regression model for each cell type, coeff_diff ~ model_species/coeff_mean + peaktype, and the numerical results are reported in **Data S5**. The regression variables are defined as follows:

coeff_diff: *τ** from using other species information minus human OCR *τ**

coeff_mean: mean *τ** from using other species information and human OCR *τ**

model_species: categorical levels rheMac10 or mm10

peaktype: categorical levels Model_Org -> Hg, Hg -> Model_Org, or OCR_Ortholog.

For LDSC partitioned heritability LDSC analyses (<https://github.com/bulik/ldsc/wiki/Partitioned-Heritability>), we sought to estimate the genome-wide enrichment of heritability of SNPs within an annotation compared to outside the annotation normalized to how many SNPs fall within the annotation and the total heritability of the trait. We computed heritability enrichment of human cell type OCRs mappable or mappable and predicted active in increasingly distant mammalian orders.

### GWAS for neurological and psychiatric disorders

We collected summary statistics of 64 genome-wide association studies (GWAS) spanning neurological and psychiatric traits related to the brain and cognitive functions along with control traits known to be involved with other tissues and organs (**Data S4**). These traits have been grouped grossly into six clinical domains of effect: neurodegenerative (Degen) [37–41], psychiatric disorders from diagnosis or with related measures (Psych) [42–50], substance abuse behavior (SU) [51–55], substance abuse disorder by diagnosis or with related measures of disordered use (SUD) [51,55–58], neurological trait not otherwise specified (Neuro) [59–67], sleep-related traits (Sleep) [68–70], metabolism traits from blood biochemistry (Metabolism) [71–73], or other traits (Other) [61,74–78]. Three GWAS (Cannabis use disorder, opioid dependence vs. opioid exposure, and post-traumatic stress disorder) analyzed in this study had been conducted in African-ancestry or admixed populations.

Complex traits have varying degrees of SNP heritability (h^2^), and genome-wide association studies to current date have varying degrees of power to detect trait-associated loci. Furthermore, the complexity and interrelatedness of complex traits prove a challenge for attributing overlap or non-overlap of cell type-specific measurements of related GWAS. We provide two auxiliary analyses to ascertain the relationship between the selected traits: estimates of the GWAS power based on confidence of the h^2^ estimation by LDSC and by genetic correlation, r_g_, (<https://github.com/bulik/ldsc/wiki/Heritability-and-Genetic-Correlation>).

### CTACIT Model training

We trained Cell Type-Aware Conservation Inference Toolkit (CTACIT) convolutional neural network models (CNN) in the caudate nucleus using cell type OCRs from snATAC-seq or cell type-sorted ATAC-seq from multiple species described below. We applied theCNN architecture as previously described in Srinivasan, Phan *et al.* (5 convolutional layers, 1 dense layer, and a sigmoid activation) and trained CNNs with a one-cycle policy (OCP) cyclic learning rate scheme [16]. Using that architecture, we specifically did a grid-search to find that the following parameters produced robust models according to cross-validation performance: batch size=1000, L2 regularization=1e-10, and dropout=0.25. Similarly, we found that the following OCP learning parameters produced the most robust models: epoch=23, cycles=2.35, learning rates varying between 1e-2 to 1e-1, and momentum varying between 0.85 to 0.99. We trained 5-fold cross-validated models using the same chromosome hold-out scheme as in our previous work [16], where OCRs on the test set (chr1 and chr2) are never seen by any models or evaluations for hyper-parameter selection, and models for each fold do not see OCRs on its own validation chromosome hold-out sets: fold1: chr[6, 13, 21], fold2: chr[7, 14, 18], fold3: chr[11, 17, X], fold4: chr[9, 12], and fold5: chr[8, 10]. In the cross-species setting, the peak assignment to test, validation, and training sets are determined by the hg38 chromosome if the peak is mappable to human, or the hg38 chromosome of the mappable peaks in synteny with the unmappable peak. To identify peaks in synteny with an unmappable peak, we sorted the peaks by the source mouse or macaque chromosome and coordinates, converted the results to a data table, and filled in the missing hg38 chromosome to peaks not mappable to hg38 by the previous or next non-missing hg38 entry by running dplyr::fill(direction = “updown”). We trained models using 1 species, human (HgOnly), to evaluate whether a model trained on one species can make accurate predictions in species not used in training [22,79] or using all 3 species, human-rhesus-mouse (HgRmMm) to evaluate the gain in model performance using cell type-matched, cross-species open chromatin data. The 3-species HgRmMm training paradigm established the final caudate CTACIT models because these models had as good or better performance than the models trained using only 1 species, a result further supported by multiple previous studies [22,80].

We trained models using the following negative sets to identify the negative set with the best balance in overall performance and specific performance at detecting cell type OCR specificity and changes in OCR conservation. All models used positive sets from the reproducible OCR set detected in their cell types across species, where we removed OCRs overlapping exons and within 20,000bp from a TSS to thoroughly exclude promoter elements, as described in [24] because promoters and enhancers within the same cell type have been shown to be bound by only partially overlapping groups of transcription factors [81–83]. The following negative sets were used to train 5-fold cross-validated models, and the nonCellEnhLargeGC negative set is the final negative set for caudate CTACIT models.

1. nonEnhNeg: We obtained orthologs of the positive set OCRs mapped to other species that did not overlap any reproducible or non-reproducible OCR of the cell type used in model training [22]. In the human-rhesus-mouse 3-species setting, the nonEnhNeg set is 40-60% the size of the positive set. This negative set aims to push models to learn cases when cell type OCR activity is lost although genome sequences can still be aligned.
2. largeGC: We obtained the large set of GC-matched negatives to the positive set within each species using the biasaway tool with the following parameters: biasaway c --nfold 10 --deviation 2.6 --step 50 --seed 1 --winlen 100 [84]. While 10 times GC-matched negatives are generated, we further filter them to exclude exonic and promoter regions and regions overlapping any reproducible or non-reproducible OCR of the target cell type, resulting in 4-5 times as many GC-matched negatives as positives. This negative set aims to account for non-specific biases due to GC-content of or other functional activity of the genome unrelated to cell type OCR activity.
3. nonEnhLargeGC: We obtained the union of negatives from nonEnhNeg and largeGC. This negative set was 4.5-5.5x larger than the positive set.
4. nonCellEnhLargeGC: We obtained the union of negatives from nonEnhLargeGC and OCRs within each species from other cell types (nonCellNeg). These additional negatives come from the positive sets of the other 7 cell types that do not overlap any reproducible or non-reproducible OCR of the target cell type, which is about 4-5 times the positive set. This negative set combines the first two and also aims to train models to learn cell type-specific OCRs rather than tissue- or cell type-invariant OCRs. When combined, the nonCellEnhLargeGC negative set is 9-11x larger than the positive set.

### CTACIT Model evaluation

We evaluated CTACIT models across folds for overall performance on the validation set using the area under the receiver-operator curve (auROC), area under the Precision-Recall curve (auPRC), area under the Precision-Recall curve subtracting by the ratio of positives to negatives (P:N) to account for various P:N ratios across models (auPRC.adj), f1 score (f1_score), false positive rate (fpr), true positive rate (tpr), and negative predictive value (npv). We also evaluated these models for cell type-specific prediction of OCRs using tpr and true negative rate (tnr), where we evaluated results for each model-cell type combination separately. In this case, tpr and tnr are calculated using the nonCellNeg validation set. Similarly, we evaluated our models’ abilities to predict whether open chromatin is conserved, in which we calculated tnrs using the nonEnhNeg validation set [22]. For metrics that required categories (e.g fpr), rather than scores (e.g. auROC), we used the threshold > 0.5 to classify a DNA sequence as open and <=0.5 as closed. Lastly, we accessed the average CTACIT predictions of cell type OCRs on average when mapped to increasing distant species, the phylogeny-matched correlations as described in [22]. The negative sets vary in size, which will affect the evaluation metrics affected by class imbalance. For the reported evaluation metrics, the following values would be the baseline (random), auROC = 0.5, auPRC.adj = 0, f1_score = 0.5, tpr = 0.5, and tnr = 0.5.

### Cross-species peak cell type annotations and predictions

We predicted open chromatin at orthologs of human caudate cell type OCRs that were mappable [28] to each Zoonomia genome. This creates a peak-by-species matrix for each cell type with CTACIT predictions, where there is a row for each human OCR for a cell type and a column for each Zoonomia genome. We entered NAs for entries in species that did not have a unique, usable DNA ortholog and recorded the CTACIT prediction if there was a unique, usable DNA ortholog. For each species, we obtained the median TimeTree estimate of million years ago (MYA) diverged from humans (e.g. rhesus macaque is 28.8 MYA, house mouse is 89 MYA) [85,86] (**Data S6**, tab: Zoonomia_Genome_Information, column: “Time Since Split from Human (TimeTree median)”). We averaged across predicted OCR ortholog open chromatin in species at the same MYA divergence from humans and within the same taxonomic order for Boreroeutherian species or taxonomic clade for Afrotherians and Xenarthrans (e.g. all predictions mappable to Chiroptera). This created 20 groupings from human to the most distant clades, which represent the predicted evolutionary history of each OCR as far as it has a mappable ortholog (**Data S6**, tab: Zoo_meta_clades, column: “group_meta”). For each cell type, we classified whether each human OCR is predicted to be accessible in each of the 20 groupings by taking the average for each group and defining predicted accessible as having a prediction of at greater than 0.5. We used these binary annotations for CTACIT LDSC analyses (**Figure 1D**, **Figure 2, Figure S11-26**).

As introduced in Srinivasan, Phan *et al.*[16], we calibrated the raw CTACIT model score by the percentage of the validation set that had that score or lower to scale the strength of CNN nonlinear sigmoid activation across model folds and cell types. This was accomplished by taking the CTACIT score of the validation positive set to create an empirical cumulative distribution (ecdf R function) to scale all new predictions by the fraction of validation positives with a lower CTACIT model score. This produced calibrated scores near 0 for the validation negative set and expandrf the dynamic range for prediction values in the positive set. We applied this calibration to the continuous score for the average across the species groupings for CTACIT predictions (**Figure 1E**, **Figure 3D, Figure S30-32**) and individual species predictions (**Figure 1F**, **Figure 4C,D**). For heatmaps of CTACIT predictions, we used the following Newick tree created by collapsing the Zoonomia species tree from Foley *et al.* [87] (Zoonomia_ChrX_lessGC40_241species_30Consensus.tree) to select species per clade groups, and we used the binomial names to obtain the each species’s silhoeutte from <http://phylopic.org> to plot alongside mammalian phylogeny trees created with ggtree and ggtreeExtra R packages [88,89]: ((Loxodonta_africana:0.1204058581,Tolypeutes_matacus:0.1229192742):0.01145639324,(((Oryctolagus_cuniculus:0.1608919201,Rattus_norvegicus:0.2965409366):0.006031885449,(Tupaia_chinensis:0.1385286196,(Galeopterus_variegatus:0.08750990202,(Eulemur_flavifrons:0.07870638791,(Callithrix_jacchus:0.05610238745,(Macaca_mulatta:0.02760672357,(Nomascus_leucogenys:0.01503824207,(Pongo_abelii:0.01184875793,(Gorilla_gorilla:0.006121144092,(Pan_troglodytes:0.004607178674,Homo_sapiens:0.004410717579):0.00190302907):0.005963570605):0.002256803519):0.00775016763):0.01406957514):0.05984338905):0.01269656686):0.002845408602):0.002347302752):0.01705378614,(Uropsilus_gracilis:0.207400545,(Rousettus_aegyptiacus:0.1237216561,(Sus_scrofa:0.1368474811,(Equus_caballus:0.07947626741,(Canis_lupus_familiaris:0.1226195774,Manis_pentadactyla:0.1145549768):0.006550057057):0.002145230337):0.002284646667):0.005799996479):0.02130276027):0.01145639324);.

### CTACIT Age to prioritized fine-mapped GWAS loci

We created a metric called CTACIT Age that quantifies the evolutionary persistence of open chromatin conservation for a human cell type OCR. Specifically, CTACIT Age for a cell type OCR is the sum of the MYA of each 20 species grouping weighted by the average calibrated predicted CTACIT score of each group (**Figure 1E**). OCRs that do not have any mappable ortholog to a group do not have CTACIT scores. We chose to call this calculation composed of CTACIT scores across clades the CTACIT Age because it is derived from species MY divergence from humans and to differentiate it from scores of a species or scores averaged over species of a clade. We assigned fine-mapped SNPs a CTACIT Age for each overlapping cell type OCR that is distal with a less strict cutoff >3kb from a TSS and non-exonic based on gene annotations of the human genome. CTACIT Age calculations are most appropriate for distal open chromatin elements. To convert CTACIT Age into quantiles for eQTL and LDSC analyses, we ordered non-exonic, distal cell type open chromatin regions by the Age value and divided the regions into equal quartiles. We had also tried quintiles, but this created too few genome-wide regions per quantile bin to estimate heritability by LDSC or other enrichment analyses.

### eQTL Dataset Processing & Analyses

To identify enrichment of genetic variants associated with gene regulation in CTACIT annotations, we used the full summary statistics of two eQTL datasets. We downloaded the first dataset, eQTLs from bulk caudate RNA-seq from the GTEx consortium v8 [90], from the GTEx google bucket at the link gs://gtex-resources/GTEx_Analysis_v8_QTLs/GTEx_Analysis_v8_eQTL_all_associations/Brain_Caudate_basal_ganglia.v8.allpairs.parquet. We downloaded the second dataset, eQTLs from single-cell single nucleus-RNA-seq of the prefrontal cortex [91], from Zenodo link <https://zenodo.org/record/6104982#.YtLARy-B1hE>. We calculated the FDR by applying the Benjamini-Hochberg method [92] to the summary statistics nominal p-values of each dataset to identify variants that were significantly correlated with allele-specific gene expression either in bulk caudate nucleus or prefrontal cortical cell types after multiple hypothesis correction. Some variants were tested multiple times with different neighboring genes, so we selected the most significant eQTL p-value for each eQTL. We annotated the variant hg38 location to be in one of the CTACIT Age quartiles or outside these annotations. We computed enrichment odds ratio that a variant is a significant eQTL (FDR < 0.05) and overlaps a CTACIT Age quartile (vs. not being in a CTACIT Age annotation) using a generalized linear model with a binomial link `signif_FDR ~ quantile` with the speedglm R-package for >10 million variants [93]. The enrichment analyses are well-powered by the >10 million of eQTL variants tested, so we used the Bonferroni correction at alpha = 0.05 to determine if eQTLs tend to be in CTACIT Age quartiles. We did the Bonferroni correction over 4 CTACIT Age quartiles by 8 CTACIT cell type annotations by 8 single-cell eQTL sets or 1 bulk caudate eQTL set (**Data S8**). For plotting, we transformed the estimated log odds ± standard error to the regular odds ratio (**Figure 3A-B, Figure 28A**).

### Polyfun fine-mapping

We applied functionally informed fine-mapping (polyfun), an approach for prioritizing candidate-causal variants from GWAS by accounting for LD and partition heritability (h^2^) at the SNP level across any set of genomic annotations [94], to the 43 traits from the above list of 64 traits with significant population overlap with the UK Biobank (**Data S4**, column “runPolyfun”). To do this, we pre-computed polyfun LD matrices on >18 million imputed SNPs [94]. We augmented the polyfun baselineLF v2.2 annotations with improved annotations of genome constraint using accelerated and conserved mammalian phyloP (top 0-4%, 8 annotations), primate PhastCons (top 0-5%, 5 annotations), conserved and accelerated phyloP at the FDR 5% threshold (PhyloP ≥ 2.270 and ≤ -2.270, 2 annotations), phastCons covering the top 3.5% of SNPs (PhastCons ≥ 0.961, 1 annotation) [95], and 6 ENCODE3 cCRE annotations from promoters, distal, and proximal enhancers [96], and we excluded 12 older primate and mammalian PhastCons annotations. Lastly, we included annotations for caudate cell type OCRs measured in humans, rhesus macaque (used human orthologs), and house mouse (used human orthologs) for a total of 24 regulatory genomic and functional conservation caudate cell type annotations. We created these annotations by lifting over bed files of each annotation from hg38 to hg19 [31] and computing LD scores using the precomputed UKBB LD matrices [94]. We excluded the few SNPs not found in the UKBB LD matrices from analyses (--allow-missing). We performed all polyfun analyses using the SuSIE as the underlying fine-mapper for a maximum of 10 candidate causal SNPs within a 1Mb locus (--method susie --max-num-causal 10) [97]. We fine-mapped any locus containing a least a SNP with GWAS p-value below a cutoff selected based on (--pvalue-cutoff) the power of each GWAS; the cutoffs were p < 5x10^-7^ for traits with few genome-wide significant loci (Alzheimer’s Disease, OCD, PTSD, OpioidExposure, AlcoholDependence, OpioidDependence, and Dozing) and p < 5x10^-8^ otherwise [97]. All polyfun runs, except Obsessive-Compulsive Disorder (OCD) and Opioid Exposure (OE_vs_UX), returned fine-mapped SNPs with non-negligible posterior probabilities of being predicted causal SNPs (PIP > 0.10), where the number of fine-mapped causal SNPs correlated with the power of each GWAS. We reported fine-mapped SNPs in **Data S9** alongside the SNP’s phyloP score. We annotated fine-mapped SNPs that overlap a non-coding and non-promoter cell type OCR with the CTACIT Age of that OCR along with the fraction of bases in the OCR that has a phyloP score ≥ 2.270 (“consFrac”) (**Data S10, Figure S27**).

We compared the fine-mapped variants associated with human traits to the CTACIT annotations. We combined our tested fine-mapped neuropsychiatric traits with similarly fine-mapped traits conducted in [95]. We excluded the fine-mapped SNPs from 'BMI,' 'Intelligence score,' 'Neuroticism,'and 'Morning Person,' traits already in our set of 43-fine mapped traits, to create a “non-brain” background set of SNPs. Because some traits were from under-powered GWAS to find few fine-mapped SNPs (e.g. Anorexia), while other traits have well-powered studies to find more fine-mapped SNPs (e.g. BMI), we grouped SNPs together by their trait groups to power enrichment analyses across 6 trait domains (“Degen,” “Psych,” “SU,” “Neuro,” “Sleep,” “non-brain,” as defined in **Data S4**). Variants may be pleiotropic–fine-mapped from multiple traits within a trait group, so we selected the highest PIP of each variant and filtered out variants with marginal probability of being fine-mapped variants as previously done (PIP > 0.10) [98]. We annotated the fine-mapped variants by the hg38 location to be in one of the CTACIT Age quartiles or outside these annotations and similarly computed enrichment odds ratio that a variant is a non-marginal fine-mapped brain variant (vs. “non-brain”) and overlaps a CTACIT Age quartile (vs. not being in a CTACIT Age annotation) using a generalized linear model with a binomial link `trait_group ~ quantile.` We had fewer fine-mapped SNPs across trait groups and CTACIT quartiles compared to >10 million eQTLs, so we used the FDR correction at alpha = 0.05 to determine if fine-mapped SNPs tend to be in CTACIT Age quartiles. We did the FDR correction over 4 CTACIT Age quartiles by 8 CTACIT cell type annotations by 5 trait groups (**Data S8**). For plotting, we transformed the estimated log odds ± standard error to the regular odds ratio. (**Figure 3C, Figure 28B**).

### Selection of conserved, active cell type enhancers

To test the prediction of conserved striatal cell type enhancer activity across species, we selected two human MSN candidate enhancers and the corresponding mouse orthologs to test *in vivo* with a reporter assay plasmid, similar to previous strategies for validating candidate cell type-specific enhancers [99]. We selected two human candidate enhancers upstream of the *DRD2* locus to validate CTACIT model predictions of conserved cell type enhancers according to the following criteria: 1) human sequences predicted active by CTACIT D1 and D2 models, 2) the mouse orthologous region are predicted active or inactive by CTACIT models, 3) have open or closed chromatin in D1 and D2 canonical medium spiny neurons in human and mouse that are consistent with CTACIT model predictions, and 4) contain fine-mapped SNPs from neuropsychiatric trait GWAS with PIP > 0.10. The first human sequence, called HSA (hg38 chr11:113567227-113567453), is an open chromatin peak summit-centered 227 bp region that contains fine-mapped SNPs rs7933981 allele G and rs61687445 allele C reported in **Figure 3B**. The mouse ortholog, called MMA (mm10 chr9:49210951-49211177), overlaps mouse D1 and D2 open chromatin region and is predicted by CTACIT models to be strongly active in mouse D1 and D2 MSNs. The second sequence, called HSB (hg38 chr11:113577834-113578060), contains rs7928017 allele C. The mouse ortholog, called MMB (mm10 chr9:49196535-49196761), does not overlap any mouse D1 or D2 open chromatin region and is not predicted by D1 and D2 CTACIT models to have conserved activity.

### Reporter Assay Sequence Predictions

We predicted the open chromatin of the orthologs of human candidate enhancer sequence in clades of placental mammals diverged from humans (**Figure 4C**) and of the shorter, central 227 bp of each open chromatin region that was used to clone into the reporter plasmids (**Figure 4D**). We padded the shorter sequences with N’s to be the 501bp required to predict open chromatin with CTACIT models. To get predictions at the level of neurons and glia for comparing to NeuN-stained mouse brains, we computed the weighted average across each CTACIT model prediction by the proportion of that cell type in the human caudate nucleus, estimated from the proportion of cell types in the *Corces et al.* snATAC-seq across 3 subjects [20]. The estimated proportions within neurons are interneurons = 7.67%, D1 MSN = 42.2%, D2 MSN = 44.9%, and D1H MSN = 5.3%. The estimated proportions within glia are astrocytes = 6.97%, microglia = 10.4%, oligodendrocytes = 76.2%, and oligodendrocyte precursors 6.46%. To calculate the weighted average standard error, we used the wt.var()function in the Hmisc R package, which calculates the weighted variance that we converted to the weighted standard error with N=4 CTACIT models for either the average neuron or glia predictions.

### Image Analyses and Statistics

We counted nuclei in each fluorescent channel using the python package DeepCell pre-trained Mesmer nuclear detection with default parameters aside from the micro-per-pixel correction factor [100] and post-processed the segmented nuclei with scikit-image [101]. We excluded nuclei towards the perimeter of the tiled image not from the mouse caudoputamen. We filtered the segmentation by shape factors that are likely artifacts rather than biological nuclei with the regionprops_table function from scikit-image v0.19.3 [102], diameter between 5 µm and 25 µm, and segmentation solidity greater than 92.5%. We assigned mCherry- and GFP-detected nuclei as NeuN+ if they had more than 5% overlap with NeuN nuclei. We summarized the segmentation across biological replicates to determine the mean mCherry nuclei per mm^2^ and standard error of the mean as plotted in Figure 4E.

We analyzed the number of mCherry detected nuclei per mm^2^ as linear mixed effect model accounting for cohort batch effects, viral transduction control GFP nuclei, and repeated imaging from each animal using the R package lme4 [103]. We applied statistical model to account enhancer of origin and species of origin: mCherry ~ Enhancer + Species + Cage + GFP + (1 | Animal_ID).

### Statistical corrections for multiple comparisons

In the *τ** LD score regression analyses, we report the p-values that cell type annotations have a non-zero conditionally independent effect. The coefficients estimate the annotation’s contribution to heritability conditionally independent from the baseline annotations and a background set of cell type OCRs and open chromatin regions. The *τ** p-value is adjusted for FDR correction across all combinations of 64 traits and 8 cell types measured in 7 cell type OCR sets, relevant to **Figure S4**, **Data S5**.

In the heritability enrichment LD score regression analyses, we report the p-values that cell type annotations have non-zero heritability enrichment. The enrichment estimates the proportion of heritability in the SNPs in the annotation over the number of SNPs in the annotation and the total heritability of the trait. This enrichment p-value is adjusted for FDR correction across all combinations of 64 traits and 8 cell types measured in 26 annotations sets (CTACIT predictions in species, PhyloP annotations, and genome mappability), relevant to **Figure 2**, **S11-26**, **Data S7**. Statistical significance is determined by FDR lower than an alpha of 0.05.

In the eQTL enrichment logistic regression analyses, we report the p-values that the log odds of significant eQTLs in CTACIT Age quartile bins has a coefficient that is non-zero. The log odds p-value is adjusted for Bonferroni correction across all combinations of 1 caudate eQTL region or 8 single cell eQTL cell types by 8 CTACIT cell type models, relevant to **Figure 3A-C**, **S28**, **Data S8**.

Supplementary Text

### Cross-species caudate cell type ortholog identification

We observed that using more highly variable features used to compute the PCA dimensionality reduction during ArchR’s iterative LSI clustering (varFeatures) enabled us to find discriminating features for separating D1 and D2 medium spiny neuron (MSN) clusters in caudate samples. (Using fewer variable features during iterative LSI clustering separated glia from neurons, producing only 1 MSN cluster.) From the clustering results that used more variable features, we were clearly able to discern the same 8 caudate subtypes in human and rhesus snATAC-seq. We aligned the He and Kleyman *et al.* D1/2 hybrid subtype, which corresponds to Corces *et al.* striatonigral MSN subtype, using marker gene activity scores (e.g. *RXFP1*, *FOXP2*, and *DRD1*, **Figure S1C-D**) and the high-dimensional co-clustering the ArchR cross-cell type “PeakMatrix'' subsetted to peaks with unique, usable OCR orthologs between human and rhesus macaque, described below [7,9,20]. All other macaque and human caudate cell types also corresponded to a unique rhesus macaque cell type cluster. These orthogonal computational approaches suggest that caudate cell types are well-conserved between humans and rhesus macaques.

When we performed the sub-clustering for mouse snATAC-seq as for human and rhesus macaques, we were unable to resolve D1 vs. D2 subtypes. Neither the aggregate cell type track plots for D1 and D2 MSN subtypes around marker genes such as *Drd1* and *Drd2* nor the per-cell gene activity scores in these genes corroborated the published CATlas snATAC-seq labels. We suspect the lower per-nuclei fragment depth (<10,000 unique fragments per nuclei) resulted in low sampling of accessible chromatin in D1- and D2-specific OCRs. However, we were able to identify the CATlas-labeled neuronal “MXD'' L2-cluster subtype (with high *Tac1*, *Foxp2*, and *Chrm4* gene activity scores found restricted to the mouse striatum) as the orthologous cell types to the primate D1H MSN cell type by marker gene accessibility (**Figure S1F**). From this CATlas snATAC-seq dataset of the mouse caudoputamen, we were able to identify the striatonigral MSN, astrocytes, oligodendrocytes, oligodendrocyte precursors, and microglia cell types. We called reproducible peaks as above for these 5 cell types. We paired these snATAC cell type OCR profiles with high quality cSNAIL ATAC-seq from validated D1-*cre*, A2a-*cre* (D2), and Pvalb-*cre* transgenic mouse lines (**Figure S1E**) [16].

We mapped the human, rhesus macaque, and mouse cell type open chromatin regions (OCRs) across species to demonstrate that the cell types share a common regulatory code. Roughly 90% of the OCRs are mappable within primates and 60% between human and more distantly related mammalian species, including mice (**Figure 1B**). We computed the motifs underlying the accessible peaks, including the MSN D1H subtype, we found a large number of cell type-specific motifs (found in fewer than 4 cell types) discovered within each species that were also identified in another species **(Figure S2**). The OCRs mappable to rhesus macaque and mouse genomes have similar proportions of promoters and distal elements, including those peaks that are shared between human-rhesus macaque and human-mouse (**Figure S3**). These data together at single cell and cell type-specific ATAC-seq strongly suggest that we are able to identify orthologous cell types across species, the cell type-specific marker peaks are shared across species, and the transcription factor motifs that make up the cell type-specific regulatory code are conserved.

### CTACIT performance evaluation

We trained CTACIT models for each caudate cell type (**Figure 1C**, **Methods**) [24] using only human sequences to determine if a model trained using data from only one species could make accurate predictions in species not used in training [22,79]. Our positive set was our human OCRs, and our negative set was the human sequences from the final negative set nonCellEnhLargeGC. These models, which refer to as HgOnly models, had decent accuracy for predicting open chromatin (auROC = 0.854 ± 0.0580, tpr = 0.751 ± 0.0916, mean ± st. dev.). Since these models were evaluated using an 9.46-to-1 negative-to-positive ratio, we also evaluated the area under the precision-recall curve (auPRC), F1 score, and false positive rate (fpr) and found that those were generally high considering how imbalanced the dataset was (auPRC.adj = 0.420 ± 0.157, f1_score = 0.829 ±, 0.0578) and had low false positive rates (fpr = 0.204 ± 0.0811). These models also achieved performance that was substantially better than expected by chance for predicting cell type-specific open chromatin.

Since previous studies have suggested that models trained on data for multiple species provide more accurate open chromatin predictions than models trained using data from only 1 species [22,80], we then used our data from human, macaque, and mouse to train CTACIT models for each caudate cell type (**Figure 1C**, Methods) [24], using OCRs from human, macaque, and mouse against the final negative set nonCellEnhLargeGC. Across the 8 cell types, the HgRmMm models had even higher performance for predicting open chromatin than the HgOnly models did (auROC = 0.880 ± 0.0528, tpr = 0.750 ± 0.0889). Here, HgRmMm models were evaluated using an 11.1-to-1 negative-to-positive ratio–150k validation negatives per cell type, 15k validation positives per cell type) and found that performance was also generally high for such an imbalanced dataset (auPRC.adj = 0.465 ± 0.177, f1_score = 0.864 ± 0.0338, fpr = 0.159 ± 0.0501). **Figure S4** and **Data S2** further breaks down the model validation set performance during negative set selection and hyper parameter tuning across all negative sets for HgOnly and HgRmMm models to show that the final negative set maintains a balance of sensitivity and specificity genome-wide to predict open chromatin activity. **Data S3** reports the general test set performance metrics of the HgOnly and HgRmMm models using the nonCellEnhLargeGC negative set.

To assess cell type-specificity of our CTACIT models, we predict OCRs accessible in only the target cell type as open and OCRs only accessible in all other cell types as closed. Based on this definition of cell type exclusive positive OCRs and union of any other OCR negatives, we identified on average 26.5k cell type validation negative OCRs and 748 cell type validation positive OCRs. HgOnly models had cell type-specific true positive rates of 0.714 ± 0.116, true negative rates of 0.757 ± 0.119, and negative predictive value of 0.988 ± 0.0103. HgRmMm models had cell type-specific true positive rates of 0.731 ± 0.0860, true negative rates of 0.799 ± 0.0707, and negative predictive value of .988 ± 0.016. The gain in cell type-specific tnr and tpr suggests that HgRmMm models are able to be more specific and sensitive. Since we also had more negatives than positives for this evaluation, we evaluated the auPRC for this task and found that it was generally high (auPRC.adj_HgOnly_ = 0.186± 0.163, auPRC.adj_HgRmMm_ = 0.245 ± 0.197). **Figure S5** and **Data S2** further breaks down the cell type-specific validation set performance during negative set selection and hyper parameter tuning across all models to conclude that the final set of models do maintain cell type specificity in OCR predictions. **Data S3** reports the cell type-specific test set performance metrics of the HgOnly and HgRmMm models using the nonCellEnhLargeGC negative set.

To assess the ability of our CTACIT models to detect species-specific open chromatin changes, we predict the activity state in cell type OCRs where the ortholog in another species is accessible or where the ortholog is inaccessible. Based on this definition of OCR cross-species change, we identified on average for a cell type 3,990 species-specific validation OCRs where in one species had lost OCR activity and 3801 validation OCRs where there is sustained activity across species. The HgOnly models had an average true positive rate of 0.757 ± 0.152, true negative rate of 0.5559 ± 0.160, and negative predictive value of 0.698 ± 0.206, **Figure S6B**. The HgRmMm models had an average true positive rate of 0.735 ± 0.0185, true negative rate of 0.624 ± 0.149, and negative predictive value of 0.713 ± 0.204, **Figure S6A**. These results suggest that, with more species-specific examples, the models are better able to detect accurate loss in cell type OCR activity across species. Curiously, the HgRmMm cross species model was better at predicting true maintenance or loss of OCR activity in rhesus macaque and mouse (tnr_hg38_ = 0.526, tnr_rheMac10_ = 0.648, tnr_mm10_ = 0.697, tpr_hg38_ = 0.612, tpr_rheMac10_ = 0.849, tpr_mm10_ = 0.743). This may reflect the underlying data quality where the human snATAC-seq samples came from frozen postmortem human brains while the rhesus macaque snATAC-seq and cSNAIL mouse ATAC-seq came from freshly dissected tissue.

When we stratified species-specific prediction accuracy across cell types, the HgOnly often successfully predicted changes in chromatin status between species, and the HgRmMm models were even more specific (**Figure S6C**). D1 MSN, D2 MSN, and Pvalb interneuron models had high auROC and auPRC across all three species OCR sets. Species-specificity auROC and auPRC in human D1H MSNs and glia were lower compared to the rhesus macaque and mouse OCRs sets. The higher auROC and auPRC in rhesus macaque and mouse suggests the models may be able to predict cross-species changes well within the primate lineage in spite of the poor human performance, which could be a result of the data coming from frozen brain tissue. **Figures S7A** and **S7B** and **Data S2** further breaks down the species-specific validation set performance during negative set selection and hyper parameter tuning across all models to conclude that the final set of models do maintain cell type specificity in OCR predictions. **Data S3** reports the species-specific test set performance metrics of the HgOnly and HgRmMm models using the nonCellEnhLargeGC negative set.

Lastly, we assessed the phylogeny-matched correlations by determining the correlation between the model predictions and the years diverged from source OCRs. We do not expect all species to use the same set of OCRs across species [22]. Furthermore, we expect the OCRs measured in one species will be less active on a genome-wide level when mapped across species in more distant species due to evolutionary drift and usage of cell type-specific gene regulation [104]. We mapped human OCRs across species and observed the expected decline in the number of OCRs that can be mapped in increasingly distant species (**Figure 1B**, **Figure S7**). This mappability may also be lower in certain species due to lower quality genomes or representation within Zoonomia [29]. From these orthologous DNA regions, we found a negative correlation with the average CTACIT raw scores of each species and the number of years diverged from humans, **Figure S8A**. When we performed the calibration on the CTACIT scores, we found that the expected negative correlation improved and were comparable across cell type models (**Methods**, **Figure S10B**). This phylogeny-matched correlation analysis suggests our models were able to predict the regulatory decay of cell type OCRs across species.

### Pairwise cross-species LD score regression

We applied genome-wide SNP-heritability enrichment analyses using stratified LD score regression (LDSC) on human neurological and psychiatric traits comparing the changes in pairwise OCR overlap in three species to human OCRs to complement enrichment analyses made in OCRs that to be active across hundreds of mammalian genomes using CTACIT. We considered annotations using human cell type OCRs, mappability of cell type OCRs to distant mammalian genomes, and conservation of cell type OCRs in model species (Methods). Neurological and psychiatric traits with a wide range of SNP heritability and power to detect genome-wide significant loci were selected across GWAS (**Figure S9A**), and their genetic architectures were shared between traits (**Figure S9B**). We estimated contributions to GWAS traits with stratified LD score regression (LDSC) using the per-SNP independent effect of having a risk SNP in cell type OCRs, τ*, conditioned on the baseline annotations of the genome (Methods). We compared the changes to τ* from partitioning GWAS traits by cell type OCRs from humans, mappable to or from rhesus macaque or mouse, and cell type OCR orthologs shared between humans and the model organism (numerical results in **Data S5**). First, we found that neuronal OCRs tend to be significantly enriched across the panel of largely brain-related traits (**Figure S10A**) compared to glial OCRs (**Figure S10B**). Neuronal cell type OCRs measured in macaque and mappable to human or measured in humans and mappable to macaque had larger effect sizes than the larger set of human OCRs (**Figure S10A**, columns 1-2, **Data S5**, linear regression slope= 0.36-0.89, FDR= 0.0044 - 4.68 x 10^-20^). OCR mappability to mouse had even larger effect sizes than human cell type OCRs (**Figure S10A**, columns 4-5, **Data S5**, linear regression slope= 0.517-1.76, FDR = 0.004-6.77 x 10^-63^). Neuronal OCRs shared between species had larger effect sizes on top of being able to map to those species alone (**Figure S10A**, columns 3,6, **Data S5**, linear regression slope= 0.65-1.24, FDR= 0.0249-2.34 x 10^-10^). These results suggest that complex brain traits are highly enriched in human neuronal OCRs that are conserved in both the DNA sequence and regulatory genomic function within orthologous cell types in rhesus macaque and mouse.

### Polyfun Fine-mapping

Based on the genome-wide enrichment for trait heritability using cell type OCR profiles from humans and model species and work on mammalian and primate genome constraint scores [95] above many other functional annotations, we expect that these additional conservation annotations will improve base-resolution fine-mapping of genetic variants associated with common traits. We applied functionally-informed fine-mapping (polyfun) with these annotations and the polyfun baseline_LF annotations for 43 brain-related traits from GWAS with significant overlap with the UKBB or European ancestry populations for which there is pre-computed LD matrices across >20 million genetic variants (MAF > 0.005) [94]. Across these traits, 41 GWAS returned 7,423 SNPs with non-marginal posterior inclusion probability (PIP) of being the candidate causal SNPs (PIP > 0.1, **Data S9**), with 748 SNPs highly confidently fine-mapped as candidate causal (PIP > 0.95).

We annotated 1,233 SNPs to regions constrained in mammals across the whole genome (phyloP ≥ 2.270, Posterior inclusion probability of being the causal SNP, PIP > 0.1) and 693 of these are constrained in mammals at distal, non-exonic regions of the human genome. In contrast, more fine-mapped SNPs, 946 unique SNPs, were distal, non-exonic and overlap a caudate cell type OCR with non-zero CTACIT Age (**Data S10**, PIP > 0.1). Both of these measures of mammalian conservation correlate with higher fine-mapping PIP scores. Although CTACIT Age and phyloP are correlated with each other, both are still correlated with higher PIP scores when adjusting for each other (p-value_phyloPcons_ = 9.99 x 10^-6^, p-value_CTACIT-Age_ = 0.0166, nested mixed effect linear regression ANOVA).

The number of non-marginal fine-mapped traits (PIP > 0.10) significantly correlates with the power of the GWAS, estimated by the z-score from h^2^ estimation, (linear regression P_h2_zscore_ = 0.00240) but not the total heritability of the traits (P_h2g_ = 0.0645). This suggests that larger GWAS in the less well-powered traits may identify more fine-mapped SNPs. Most fine-mapped SNPs were from well-powered neurological traits that tend to be easier to phenotype across large populations (Neuro_avg_ = 547 SNPs per trait, BMI=2,823 SNPs, PIP > 0.10). Psychiatric traits that require more expertise to diagnose and report had mere fractions in number of fine-mapped SNPs as neurological traits (Psych_avg_ = 168 SNPs per trait, 586 Schizophrenia SNPs, PIP > 0.10, **Figure S27**). Substance abuse and substance abuse disorder traits had the fewest fine-mapped SNPs per trait (SU_avg_ = 40.3 SNPs per trait, SUD_avg_ = 5.6 SNPs per trait). Many of these GWAS have yet to reach the point of detecting all trait-associated loci [105], so functionally informed fine-mapping and interpretation of fine-mapped SNPs will need to be performed in the future when more of these GWAS become sufficiently powered. We provide the polyfun annotations from this study online (**Data Availability**) to facilitate re-fine-mapping of subsequent GWAS.

### Additional fine-mapped loci

We used the CTACIT Age to improve fine-mapping of SNPs from several well-powered GWAS. In the neuroticism locus around the *GRM3* gene (**Figure S32**), many SNPs were fine-mapped to be candidate causal SNPs with varying degrees of confidence. The top two SNPs, rs274622 and rs802425, had nearly equal PIP scores, yet only the latter SNP overlapped a D2 MSN OCR with high CTACIT D2 Age (chr7:86,665,609, D2 Age=38). The first SNP may still be a candidate causal variant, but it may be involved through a different cell type or perhaps different mechanism. Both SNPs were found by GTEx to be eQTLs for *GRM3* within the cerebrum GTEx tissue and *GRM3* expression was enriched in D2 MSNs of the macaque [9,90]. We pose that both variants may be functional.

We used CTACIT to refine the fine-mapped loci from moderately powered GWAS. We previously reported that smoking-related traits had genome-wide enrichment for D1 and D2 mouse neuron subtypes from the striatum [16] (**Figure S33**). We improved our tissue and cell type predictions with CTACIT MSN_D1 models and predicted that the SNP rs148428140, from the SmokingCessation and CigarettesPerDay GWAS, had the highest CTACIT D1 Age within D1 OCRs in the region (chr9:133,609,189, D1 Age=42) (**Figure 3A**). This D1 OCR had high predicted activity in almost every species compared to other OCRs in the locus.

We investigated a locus associated with Alzheimer’s Disease (AD) in AD and AD-by-proxy GWAS, which had strong, conserved neuroimmune signatures [106] (**Figure S34**). We found two SNPs, rs2632516 and rs2526377, within microglia OCRs with high CTACIT Microglia Ages, (chr17:58,331,292, Microglia Age: 44; chr17:58,332,310, Microglia Age: 15, respectively). These SNPs and other fine-mapped SNPs in the locus fell in regions constrained across mammals (phyloP ≥ 2.270) and overlapped with human microglia OCRs. High CTACIT Microglia Age points to these SNPs as likely candidate causal variants contributing to AD. Both SNPs were eQTLs for the nearby gene *TSPOAP1* and a gene outside the plotted region, *TRIM3*, in blood and skin GTEx tissues [90]; rs2632516 is an eQTL in peripheral blood cells of systemic lupus erythematosus patients for the microRNA miR-142 [107], which regulates other AD-associated gene regulation when manipulated in human induced pluripotent neural progenitor cells and mouse hippocampus [108]. These examples and many other loci exist where CTACIT cross-species predictions and CTACIT Age could suggest which SNPs in cell type OCR may be active to prioritize downstream testing (**Data S6**). Regulatory conservation inferences may guide how human cell type-specific genetic variant could be studied in orthologous OCRs of model and non-model species (e.g. rodents, non-human primates, prairie voles)

These examples and many other loci exist where CTACIT cross-species predictions and CTACIT Age could suggest which SNPs in cell type OCR may be active to prioritize downstream testing (**Data S10**). Regulatory conservation inferences may guide how human cell type-specific genetic variants could be studied in orthologous OCRs of model and non-model species (e.g. rodents, non-human primates, prairie voles).

### Limitations

CTACIT, which was introduced by [24], is the first method to associate differences in regulatory activity between species in specific cell types. CTACIT and the experimental validations we demonstrate in this paper relate the regulatory conservation of cell types across placental mammals to the heritability of human traits in ways that are useful but have limitations.

A major limitation of CTACIT is that species, sample quality, sample preparation, research facilities, and data annotation are often confounded with each other. To the best of our abilities, we replaced or augmented the data to represent the highest-quality cell type OCR profiles from human, rhesus macaque, and mouse. The lower performance in our cross-species analyses of glia and D1H MSN cell types in humans but not rhesus macaque suggests that our model training approach may need to be further optimized, that the regulatory code in some of the cell types for which we have models is only partially conserved between euarchonta and glires, or that there are confounding factors due to differences between the labs in which the data were generated that prevented some of our models from making consistently accurate clade-specific open and closed chromatin predictions. These sub-optimally accurate clade-specific open and closed chromatin predictions decrease the reliability of results. Due to the challenges in obtaining open chromatin from most cell types and inherent differences in species’ lifestyles, standardizing conditions across species for data generation is often impossible.

Our expansion of CTACIT has additional limitations beyond sub-optimal performance of some models for some evaluation metrics. We made predictions across the placental mammalian phylogeny, but our models were trained using only species from primates and mouse, so the regulatory code that they learned may not be conserved in all of the species where we made predictions. Determining if the regulatory code in specific cell types is conserved beyond euarchontoglires requires regulatory genomics data from specific cell types in at least one mammal outside of euarchontoglires, which may be infeasible to collect due to the difficulty of obtaining tissue samples from most non-euarchontoglires mammals. In addition, we trained our models using only sequences at least 20,000 bp from the nearest TSS so that they would learn the enhancer and not promoter regulatory code, but, for some of our analyses, we made predictions at OCRs at least 5,000 bp from the nearest TSS to increase the number of OCRs we considered; some of these predictions may be inaccurate because some of these OCRs may be promoters, and further research is needed to improve the ability of convolutional neural network models for regulatory DNA sequence prediction to distinguish between promoters and enhancers. Furthermore, many neurological conditions may be caused by regulatory elements that are accessible only when a neuron is activated, so such regulatory elements may not be present in our data [109].

Our models make binary predictions, but open chromatin in a cell type representing multiple cells is a continuous signal; an exciting extension to this work would be to predict open chromatin signal across species [80] and compute an improved CTACIT Age metric that incorporates the level of predicted open chromatin. We anticipate that, as experimental technology for obtaining cell type-specific [110] and single-cell [111] regulatory genomics data as well as our ability to distinguish between promoters and enhancers and compare open chromatin signals between species improves, CTACIT’s reliability and applicability will increase.

We presented *in vivo* cell type-specific reporter assays that demonstrate the strength of CTACIT predictions, yet these experiments are limited by the low throughput and method to ascertain cell type-specificity. We took a targeted approach to test the conserved regulatory activity by testing two candidate sequences using both human and mouse orthologs. The experimental burden to synthesize, clone, package into high titer AAV virus that crosses the blood brain barrier, and transduce entire brain regions precludes testing more human loci or open chromatin regions or more orthologs from distantly related mammalian species. We were size-limited by DNA synthesis technologies and only tested the central 227bp of the candidate enhancers rather than the full 501bp sequence. Our computational models predicted the shorter 227bp to be weaker and less cell type-specific than the full 501bp sequences (**Figure 4C**, mouse and human squares vs. **Figure 4D**), a result that experimentally aligns with previous reports on how the length of synthesized candidate enhancer affects the measured strength and variation of reporter gene transcription in reporter assays [112]. The peripheral DNA sequences–in combination with the central region–may contain cell type-specific regulatory motifs or stabilize enhancers during transcription. Our reporter assay experiments leverages AAV technology to efficiently transduce high copy-numbers of our candidate enhancers into the mouse brain, but this measures only the candidate enhancer in an episomal context rather than in the native genome. Orthogonal approaches using CRISPR-knockout or CRISPR-inactivation combined with cell type targeting can be useful to confirm our results [113,114].

Within the field of rodent neurobiology that studies the caudoputamen, transgenic reporter strains are the cornerstone to selectively measure or manipulate D1 and D2 MSNs. D1 and D2 receptors have strong homologies with other G-protein coupled receptors, including D3, D4, and D5 receptors, so few antibodies have been successfully validated to specifically label D1 or D2 MSNs [115]. The membrane-trafficking property of dopamine receptors proves an additional challenge to using D1 or D2 antibodies as cell type-specific counterstains in our reporter assays. Since we predicted our candidate enhancer sequences to be equally active in both D1 and D2 MSNs but differentially active in neurons vs. glia, we circumvented these problems by using antibodies against the nuclear NeuN stain to counterstain caudoputamen neurons, which are overwhelmingly D1 or D2 MSNs [19]. Ascertaining finer cell type-specific impacts of other human candidate enhancers would have to rely on single- or double-labeling D1 or D2 reporter rodent strains [116,117], which excludes testing candidate enhancers in other species, or using single-cell transcriptomics methods, which add different sets of challenges [118,119].

Our *in vivo* experiments tested the gene regulation of mouse brains, irrespective of behavioral stratification, environmental exposures, or experimental conditions. Future work could expand these *in vivo* experiments to identify interactions between human risk variants, gene regulatory mechanisms, and the specific neural circuits underlying behavior in rodents, primates, or non-traditional model organisms of traits and behaviors. Some experiments can be paired with established paradigms such as in intravenous drug self-administration for substance use and withdrawal behaviors or multi-arm bandit gambling tasks for reward and risk-taking behaviors. The translational models on the behavior side complement the regulatory predictions provided by CTACIT to dissect the gene-to-behavior relationships in specific neural circuits of the brain that ultimately are key to understanding the human genetics of complex traits and behavior.

Several human traits that we analyze–especially schizophrenia–do not have known related behavioral paradigms or phenotypes measurable in other species. While we find increased heritability enrichment in measured or predicted active regulatory elements mappable from distant species, we do not presume that those species have these traits. Rather, our findings suggest that, across mammalian evolution, the conserved regulatory elements contribute to increased fitness, so genetic variation at these elements in the human genome may predispose humans to diseases or disorders. Experimental manipulation of these conserved regulatory elements in behaving model organisms may be able to reveal the biological mechanisms and pathophysiology of human genetic variants, as animal knockout models have done for rare mono-genetic human disorders [120]. For example, investigation of the locus from **Figure 3D,** and **Figure 4A**, a region containing eQTLs of DRD2 and accessible in human and mouse D2 neurons, may identify pathological regulation of the D2 receptors, altering critical basal ganglia dopamine signaling.

Supplemental Figures

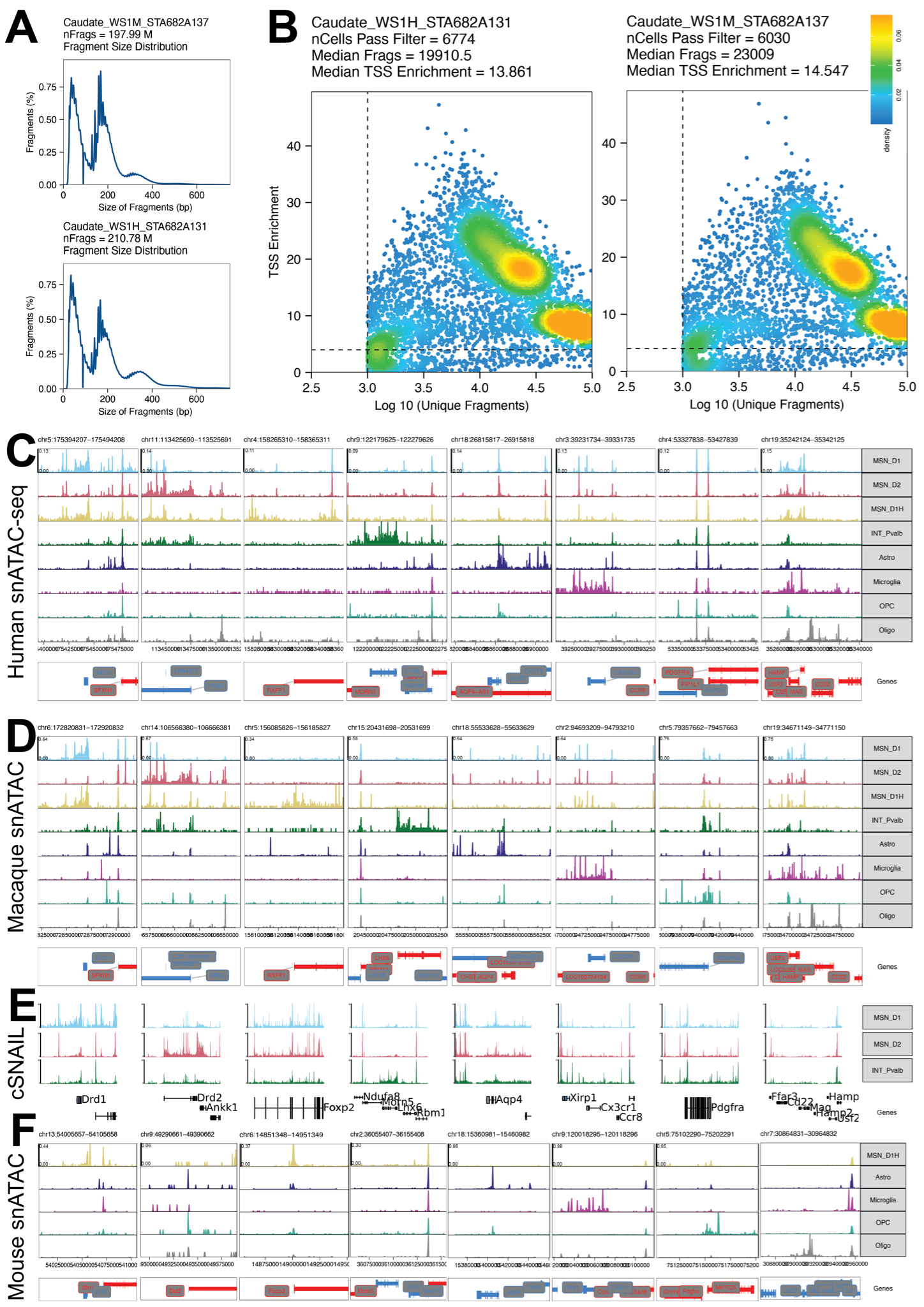

#### Figure S1. Identification of caudate cell types in human, macaque, and mouse snATAC-seq

(A) Periodicity plots of 2 technical replicates of rhesus macaque caudate snATAC-seq libraries show high quality fragment distribution matching expedited pitch and periodicity of intact chromatin.
(B) Quality control plots of per cell number of fragments and transcription start site enrichment score.
(C) Pseudo-bulk ATAC-seq signal plots generated with ArchR around marker gene loci in 8 caudate cell types from human snATAC-seq support cell type-specific of identified clusters from Figure 1A [7,20]. Canonical marker genes for each track are in order *DRD1*, *DRD2*, *RXFP1*, *LHX6*, *AQP4*, *PDGFRA*, and *MAG* to collectively discriminate the 8 caudate cell types, as previously reported [9].
(D) Pseudo-bulk ATAC-seq signal plots around marker gene loci in rhesus macaque caudate snATAC-seq. Canonical marker genes for each track are the same as (C).
(E) ATAC-seq signal plots around marker gene loci mouse caudoputamen using cSNAIL in D1-, D2-, and Pvalb-cre mouse strains to label MSN_D1, MSN_D2, and INT_Pvalb neuronal nuclei [16]. Canonical marker genes for each track are in order *DRD1*, *DRD2*, *FOXP2*, *LHX6*, *AQP4*, *PDGFRA*, and *MAG* to collectively discriminate the 8 caudate cell types.
(F) Pseudo-bulk ATAC-seq signal plots around marker gene loci mouse caudoputamen snATAC-seq for MSN_SN, and glial cell types. Marker gene tracks are the same as (E). MSN_D1: D1 medium spiny neurons, MSN_D2: D2 medium spiny neurons, *PVALB*/*Pvalb*+ interneurons, Astro: astrocytes, Oligo: oligodendrocytes, OPC: oligodendrocyte precursors.

| **celltype** | **Shared motifs identified across species** |
| --- | --- |
| MSN_D1 | AR, CENPB, GATA3, GFI1, HLF, HOXB1, HSF2, HSF4, LBX2, MEF2B, MGA, MYBL2, NFATC1, NKX2-6, NPAS4, PAX6, RHOXF1, SIM1, SIX5, SOX2, STAT1, STAT3, STAT4, STAT6, TBX1, TBX10, TBX15, UBP1, XBP1, ZNF274, ZNF282, ZNF75A, ZNF784 |
| MSN_D2 | AR, CEBPA, CEBPB, CEBPD, CENPB, GATA3, GFI1, HLF, HOXB1, HSF2, HSF4, IRX2, IRX5, MGA, MYBL2, NFATC1, NKX2-6, NPAS4, NR1I3, OVOL1, PAX6, SIM1, SIX5, STAT1, STAT3, STAT4, STAT6, T, TBX1, TBX10, TBX15, UBP1, XBP1, ZNF274, ZNF75A, ZNF784 |
| MSN_SN | ALX3, CENPB, DLX3, DLX5, EMX1, EMX2, EVX1, GBX2, GSX1, GSX2, HESX1, HMX1, HOXD1, HOXD3, LBX1, LBX2, MEOX2, MGA, NOTO, OTP, PHOX2B, POU2F2, POU3F4, POU6F2, RAX, RHOXF1, TBX10, TFEB, VAX1, VAX2, VSX1, VSX2, ZNF784 |
| INT_Pvalb | BHLHE22, BHLHE23, HOXB1, NEUROG1, NEUROG2, NEUROG3, POU3F3, TFEB, UBP1, XBP1 |
| Astro | EN1, HNF4A, HSF1, NFKB1, NR1I3, NR2C1, NR5A2, PGR, PRDM4, RARA, RARG, RELA, RXRA, THRA, ZKSCAN3, ZNF75A |
| Microglia | ELF5, ETV2, HSF1, IRF1, IRF3, IRF6, NR1I3, NR2C1, NR2F6, PRDM1, RARA, RELA, SPIB, SPIC, STAT2, ZNF282 |
| OPC | RXRA, SOX10, ZKSCAN3 |
| Oligo | HNF4A, NEUROG1, NEUROG2, NEUROG3, NR2C1, NR5A2, PGR, RARA, RUNX3, RXRA, SOX10, SOX11, SOX12, SOX13, SOX14, SOX15, SOX17, SOX18, SOX21, SOX3, SOX5, SOX6, SOX7, SOX8, SOX9, SRY, TCF7, THRA, ZKSCAN3 |

#### Figure S2. Shared regulatory code in cross-species motif analyses of distal open chromatin regions

MEME-chip significantly enriched transcription factor binding sites identified to be shared in each cell type across human, rhesus macaque, and mouse caudate cell type.

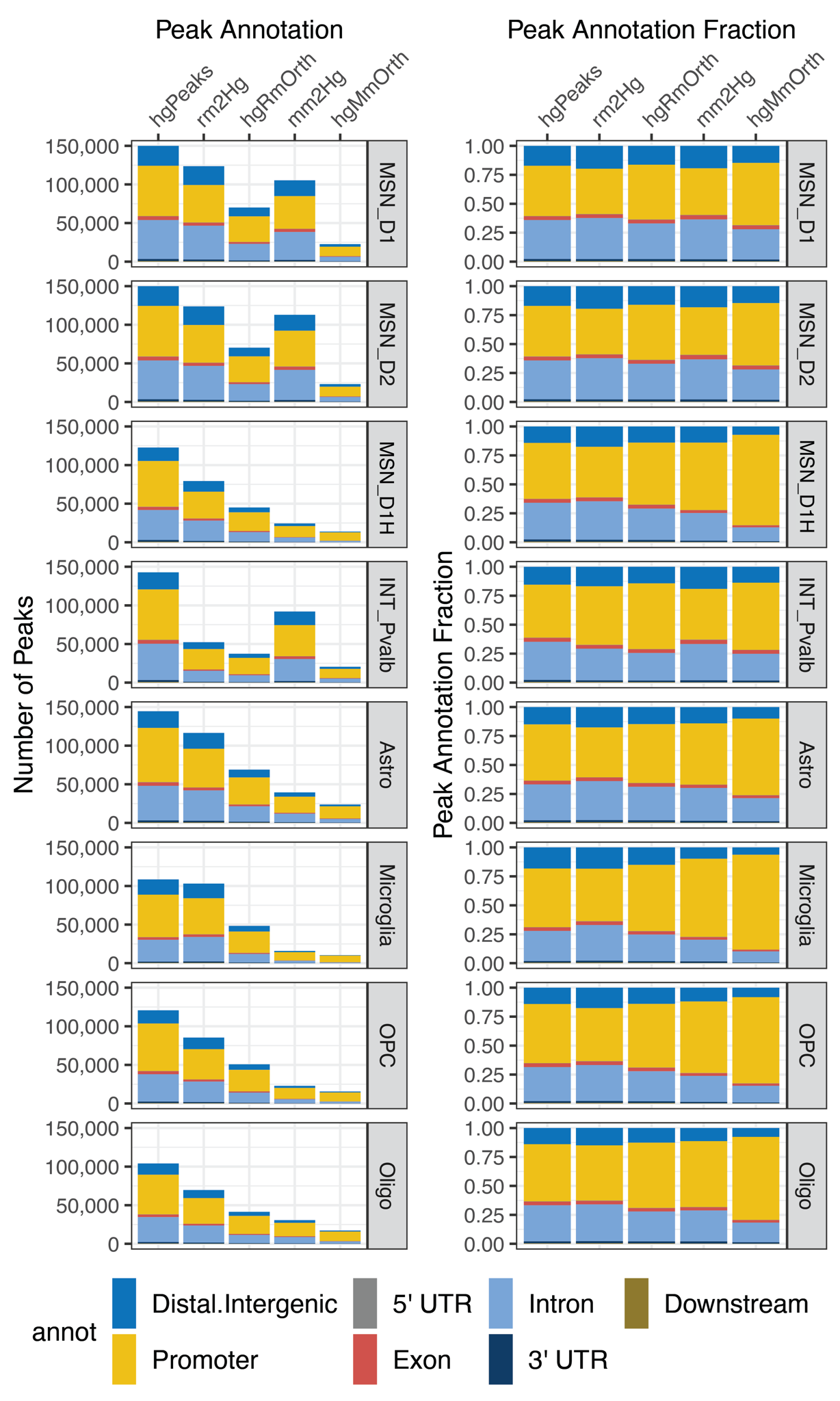

#### Figure S3. Human cell type OCRs are less mappable across more distantly diverged species

(Left) The number of human cell type OCRs that can be mapped to another species. Each point is a species. The x-axis represents the number of million years (MY) that species diverged from humans. The point color is the membership of the species with a mammalian taxonomic order. (Right) the proportion of all human cell type OCRs that are mappable to distantly related species.

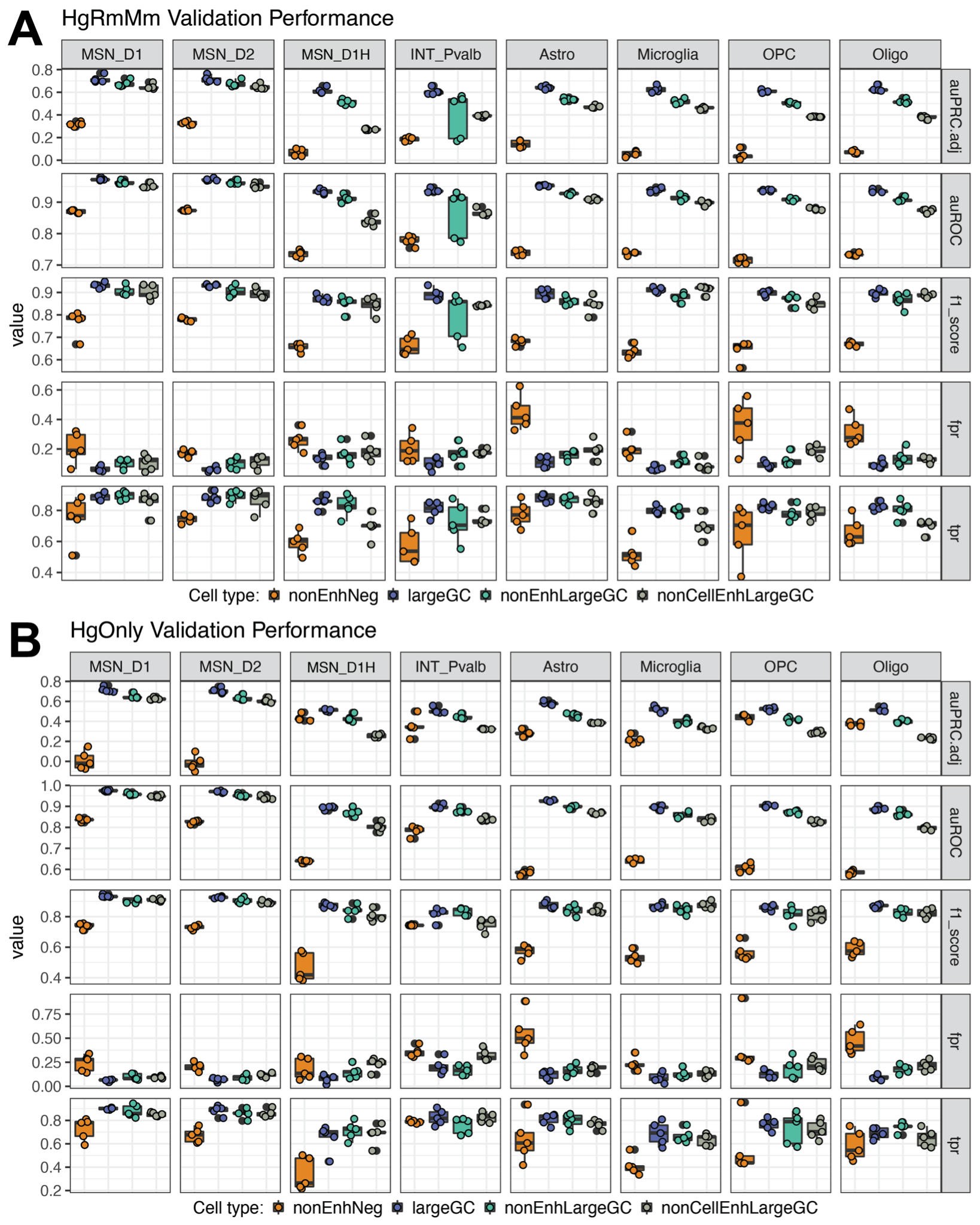

#### Figure S4. CTACIT caudate cell type models cross-validation performance with single- and multi-species data across various negative sets.

Negative sets evaluation of orthologs of cell type OCRs in other species that are not accessible (nonEnhNeg), large set of GC-matched genomic regions (largeGC), union of the nonEnhNeg and largeGC (nonEnhLargeGC), union of nonEnhLargeGC, and accessible regions in other cell types (nonCellEnhLargeGC). (A) Overall cross-validation performance metrics of CTACIT models trained with OCRs detected in 3 species across a number of negative sets (HgRmMm). (B) Overall cross-validation performance of models trained with only human sequences (HgOnly). Each point is a different cell type, cross-validation fold, negative set combination. Metrics (details in Methods): area under the Precision-Recall curve adjusted for the proportion of the minority class (auPRC.adj); area under the receiver-operator curve (auROC); false positive rate (fpr); true positive rate (tpr).

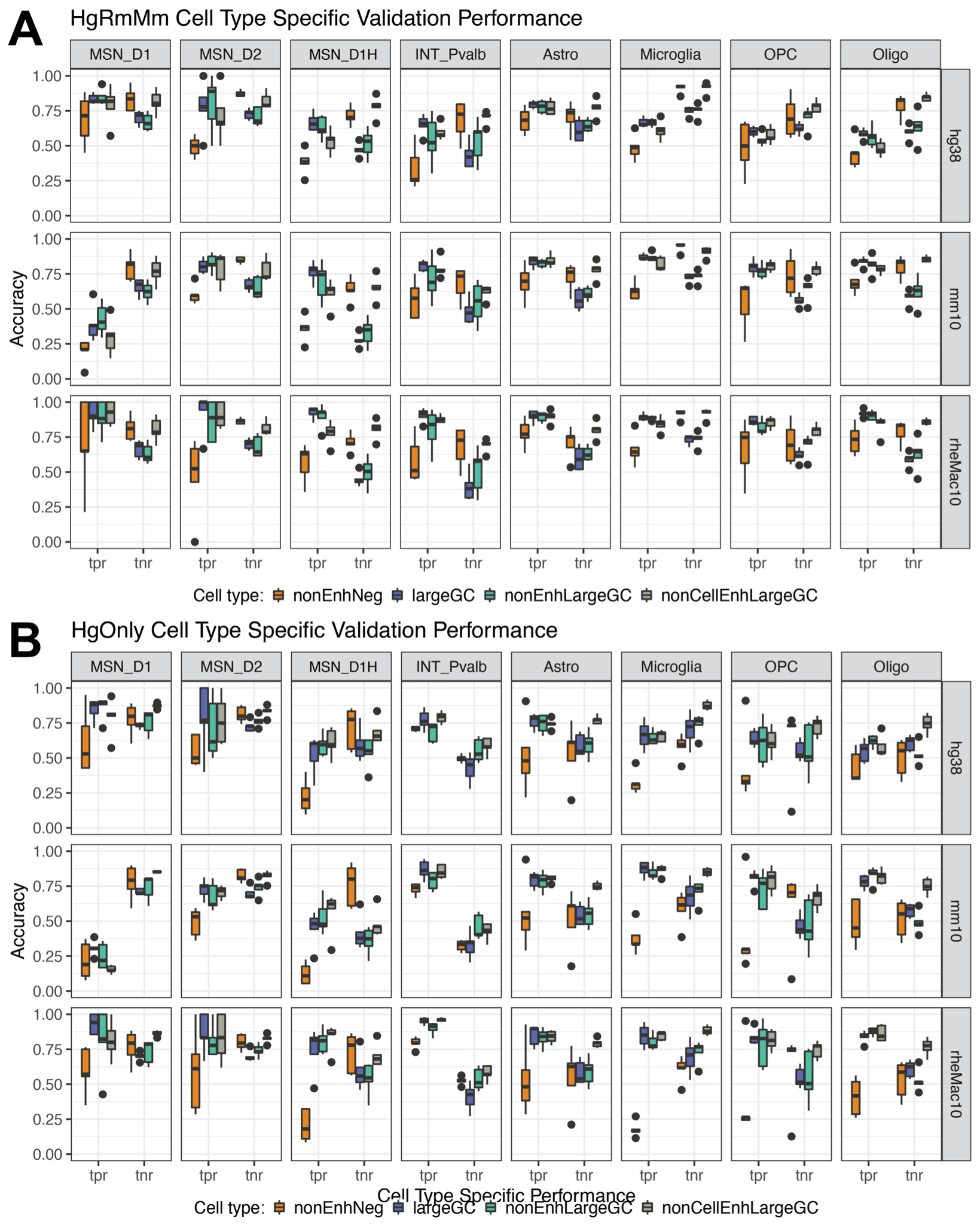

#### Figure S5. Negative set comparisons across subsets of data designed to evaluate models’ ability to predict cell type-specific open chromatin.

Negative sets evaluation of cell type-specific positive and negative cross-validation for CTACIT models trained using negative sets from orthologs of cell type OCRs in other species that are not accessible (nonEnhNeg), large set of GC-matched genomic regions (largeGC), union of the nonEnhNeg and largeGC (nonEnhLargeGC), union of nonEnhLargeGC and accessible regions in other cell types (nonCellEnhLargeGC).

**(A)** Cell type-specific positive and negative cross-validation performance metrics of CTACIT models trained with OCRs detected in 3 species across a number of negative sets (HgRmMm).

**(B)** Cell type-specific positive and negative cross-validation performance metrics of CTACIT models trained with OCRs detected in models trained with only human sequences (HgOnly).

Each point is a different cell type, cross-validation fold, negative set combination.

Metrics (details in Methods): true positive rate (tpr); true negative rate (tnr).

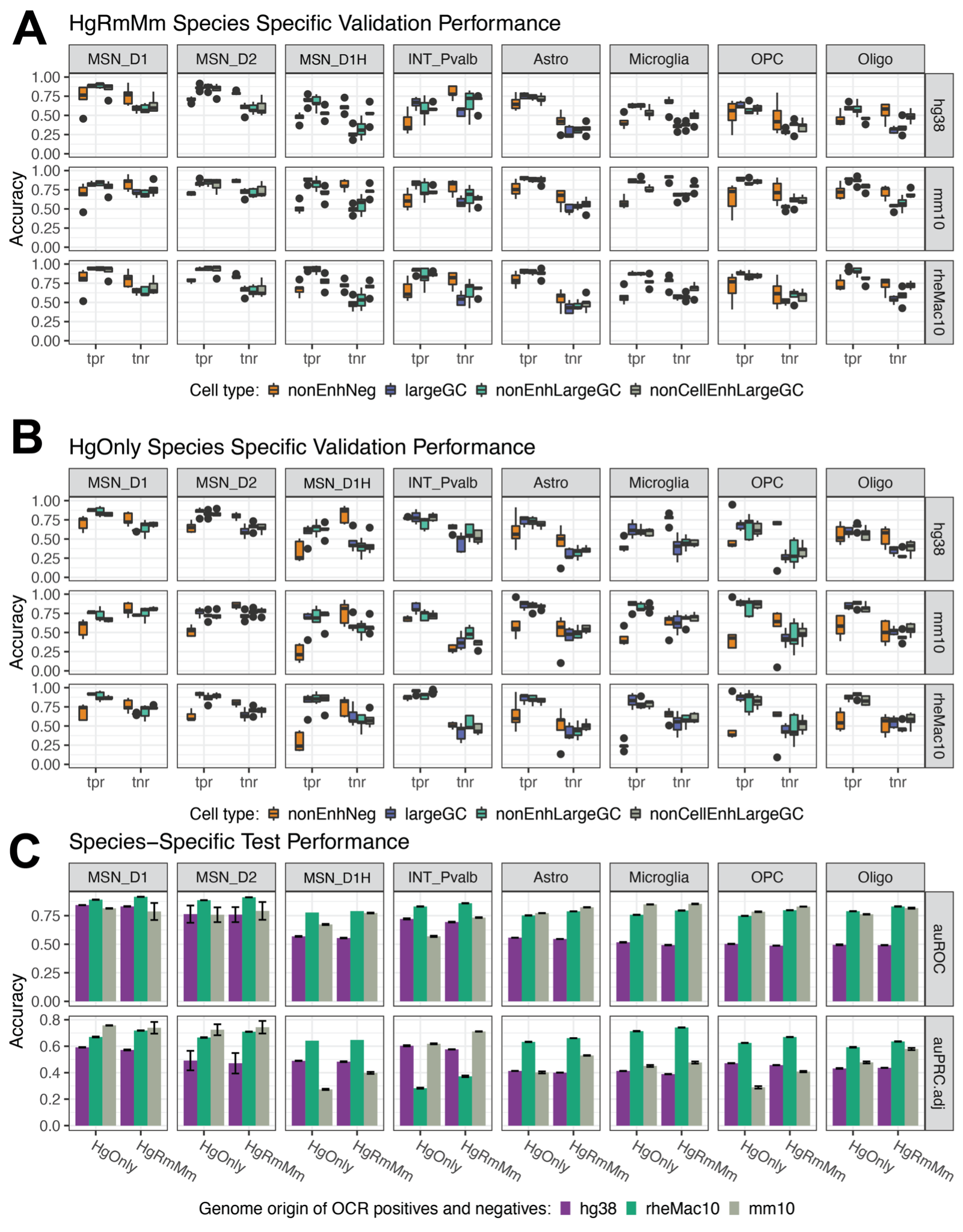

#### Figure S6. Negative set comparisons across subsets of data designed to evaluate models’ ability to predict species-specific open chromatin changes.

**(A)** Species-specific positive and negative cross-validation performance metrics of CTACIT models trained with OCRs detected in 3 species across a number of negative sets (HgRmMm). **(B)** Cell type-specific positive and negative validation set performance metrics of CTACIT models trained with OCRs detected in models trained with only human sequences (HgOnly). **(C)** Test set performance using the species-specificity positive and negative sets using auPRC.adj and auROC across both HgRmMm and HgOnly models show both models have similar species-specific prediction accuracies. In parts **(A)** and **(B)**, each point is a different cell type, cross-validation fold, negative set combination. Positives come from OCRs in one species whose ortholog is still accessible in another species. Negatives are inaccessible regions whose orthologs are accessible in other species. Metrics (details in Methods): true positive rate (tpr); true negative rate (tnr); area under the precision-recall curve adjusted for the proportion of the minority class (auPRC.adj); area under the receiver-operator curve (auROC).

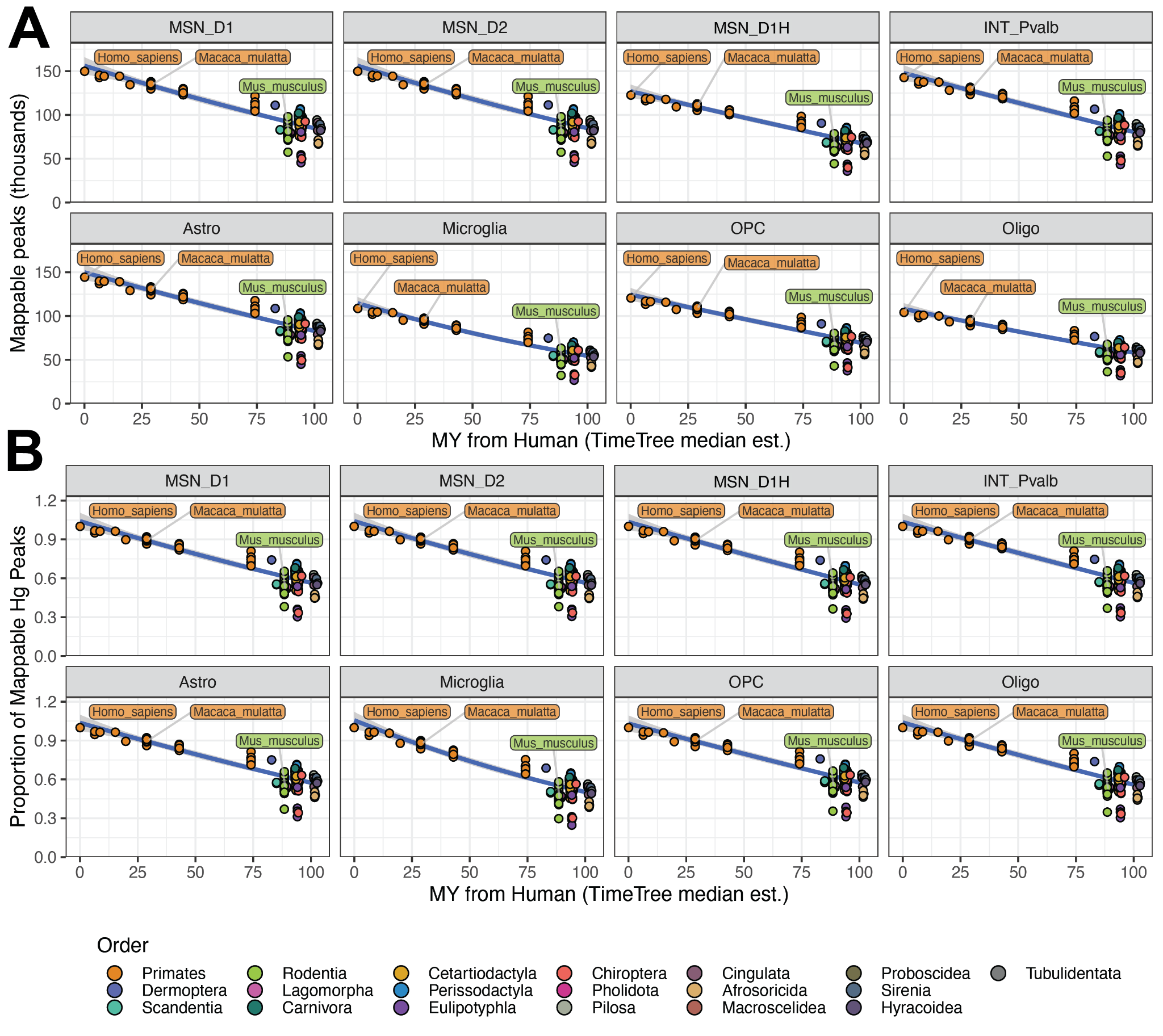

#### Figure S7. Mappability of macaque and mouse cell type OCRs to the human genome and overlap with human cell type OCRs.

Cell type orthologs (OCRs in a cell type shared across humans and another species) are a subset of peaks mappable from the other species. The relative proportions of annotations of cell type OCRs are generally similar across species and cell types. hgPeaks: human cell type OCRs; rm2hg: macaque peaks mappable to the human genome; hgRmOrth: cell type OCRs accessible in both human and macaque; mm2hg: mouse peaks mappable to the human genome; hgMmOrth: cell type OCRs accessible in both human and mouse. UTR: untranslated region, annot: annotation.

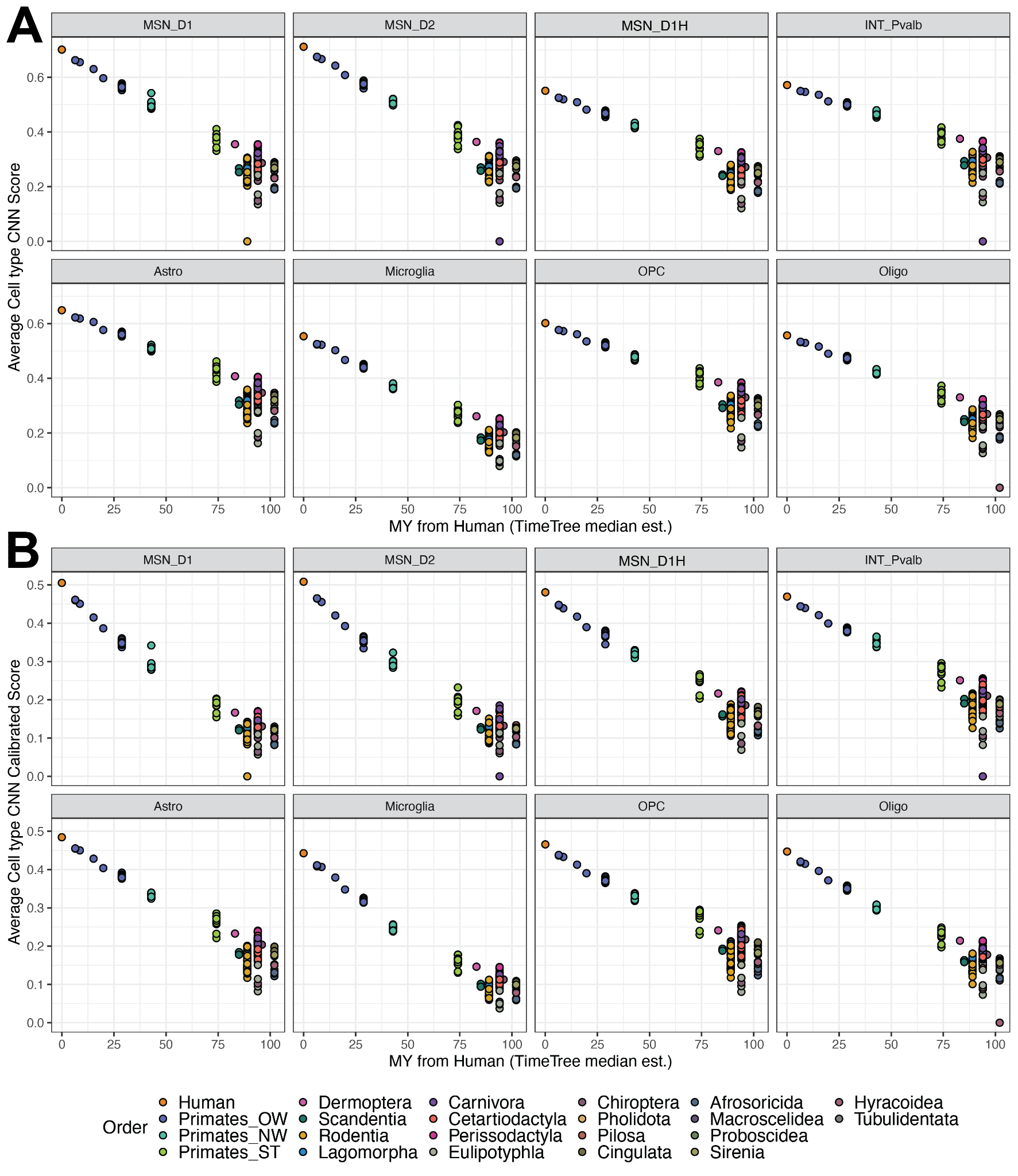

#### Figure S8. CTACIT predictions of human cell type OCR orthologs in distant species reflect sequence-level drift and loss of regulatory function with species divergence.

**(A)** The average CTACIT scores of cell type orthologs of human OCRs in each species versus the million years (MY) that species diverged from humans. Each point is a species and the point color is the membership of the species with a mammalian taxonomic order. **(B)** the CTACIT scores that are calibrated to the percentage of the model validation positives with a lower score. More negative correlations are seen with these calibrated scores and the MY diverged from humans.

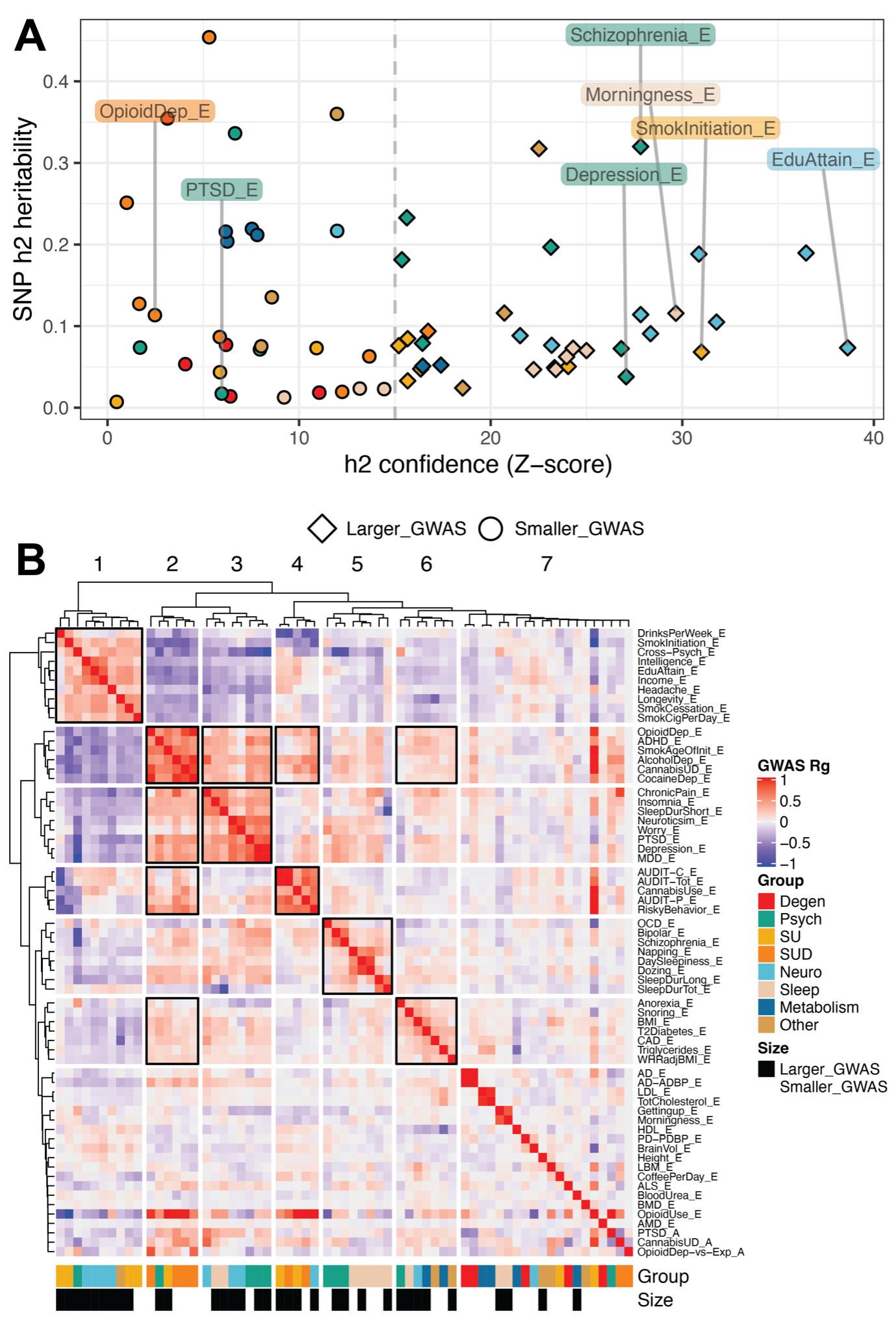

#### Figure S9. Shared genetic architecture of neurological and psychiatric traits from GWAS of varying heritability and power.

(A) Scatter plot of LD score regression-estimated SNP heritability (h_2g_) by the confidence of estimation (h_2g_ z-score) across 64 GWAS. Each study is a point colored by the trait group (legend in panel B). Round points with h_2g_ z-score < 15 indicate smaller, less well-powered GWAS. Diamond points with h_2g_ z-score > 15 indicate larger, more well-powered GWAS.
(B) Genetic correlation heat map clustered with hierarchical clustering into 7 trait groups with shared genetic architectures. GWAS are labeled by the trait and the source population (_E for European ancestry subjects and _A for African ancestry subjects). Black-outlined blocks indicate blocks of correlated traits for which more than 70% of the trait-trait correlations have a positive genetic correlation (r_g_) greater than 0.10. Traits are also annotated with group traits and whether the trait had a larger or smaller GWAS from (A).

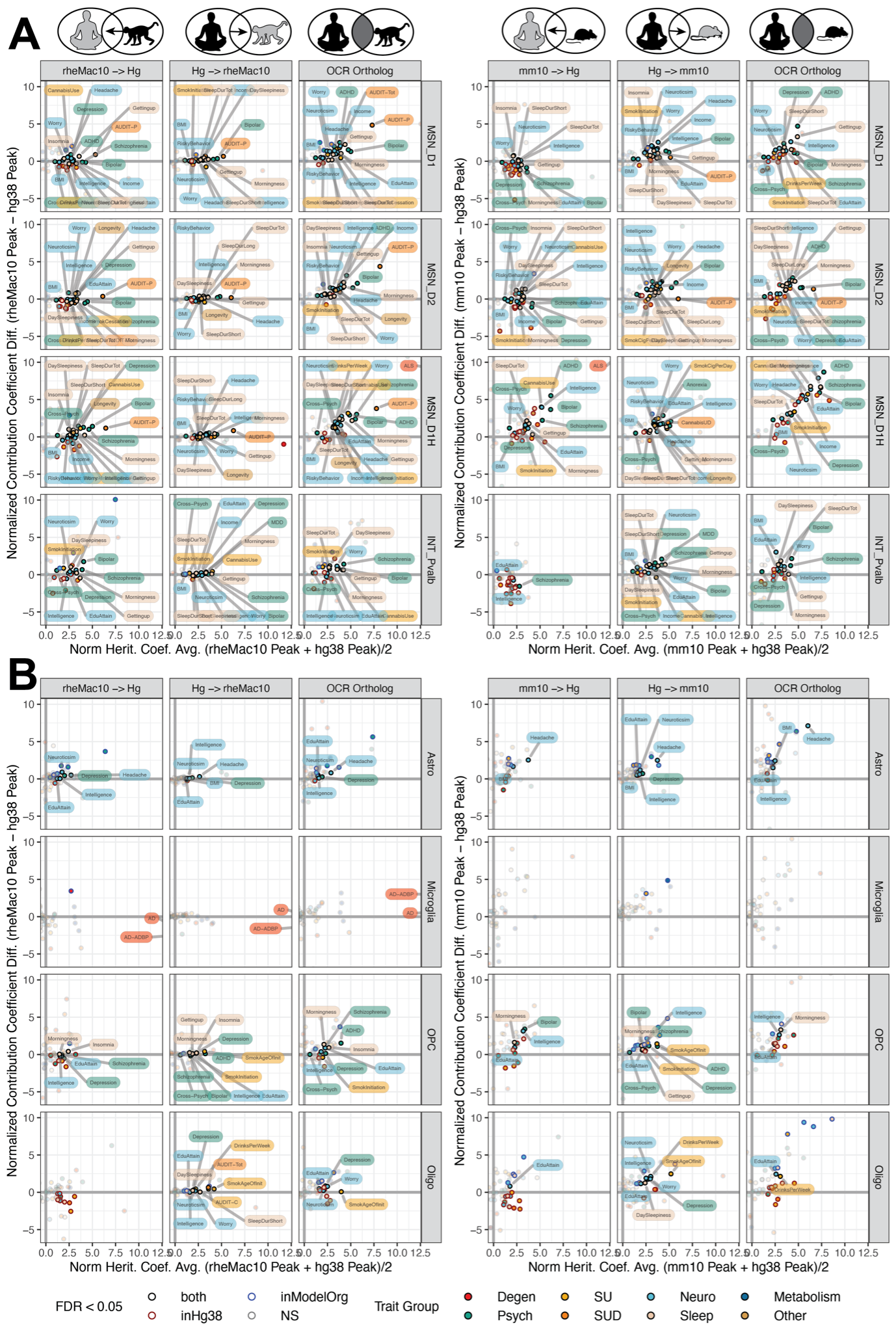

#### Figure S10. Pairwise human-model species caudate cell type OCR comparisons demonstrate complex traits enriched within conserved open chromatin in the human genome.

Conditional independence (*τ**, normalized heritability coefficient) LD score regression (LDSC) analyses of caudate cell type open chromatin regions (OCRs) using human, rhesus macaque, and mouse open chromatin. Each point is a GWAS colored by the trait group (Data S1). The *τ** was estimated for each GWAS in 7 OCR sets for each cell type: 0) human OCRs; 1) rhesus macaque OCRs mapped to human genome, rheMac10 -> Hg; 2) human OCRs that can be mapped to rhesus macaque, Hg -> rheMac10; 3) overlap of human and rhesus OCRs and OCR orthologs 4) mouse OCRs that can be mapped to human, mm10 -> Hg; 5) human OCRs that can be mapped to mouse, Hg -> mm10; and 6) overlap of human and mouse OCRs and OCR ortholog. The *τ** for OCR sets mappable or open chromatin conserved in other species (1-6) are compared to human OCRs (0) using a mean-difference plot where the average *τ** are plotted on the x-axis (normalized heritability coefficient average) and the change in *τ** with mappability or orthology in model species are plotted on the y-axis (normalized heritability coefficient difference). Points above the y=0 line demonstrate higher *τ** with genome or open chromatin conservation information above human cell type OCRs alone. Significant *τ** LDSC estimates are outlined with red for significance in human OCRs (inHg38), significant in Model Species (inModelOrg, 1-6), or both (FDR < 0.05, computed over all 8 cell types, 64 traits, 7 OCR set comparisons). Traits that does not have significant *τ** estimates are plotted with transparency. **(A)** Neuronal cell type comparisons: MSN_D1, MSN_D2, MSN_SN, and INT_Pvalb. **(B)** Glial cell type comparisons: Astro, Microglia, OPC, and Oligo.

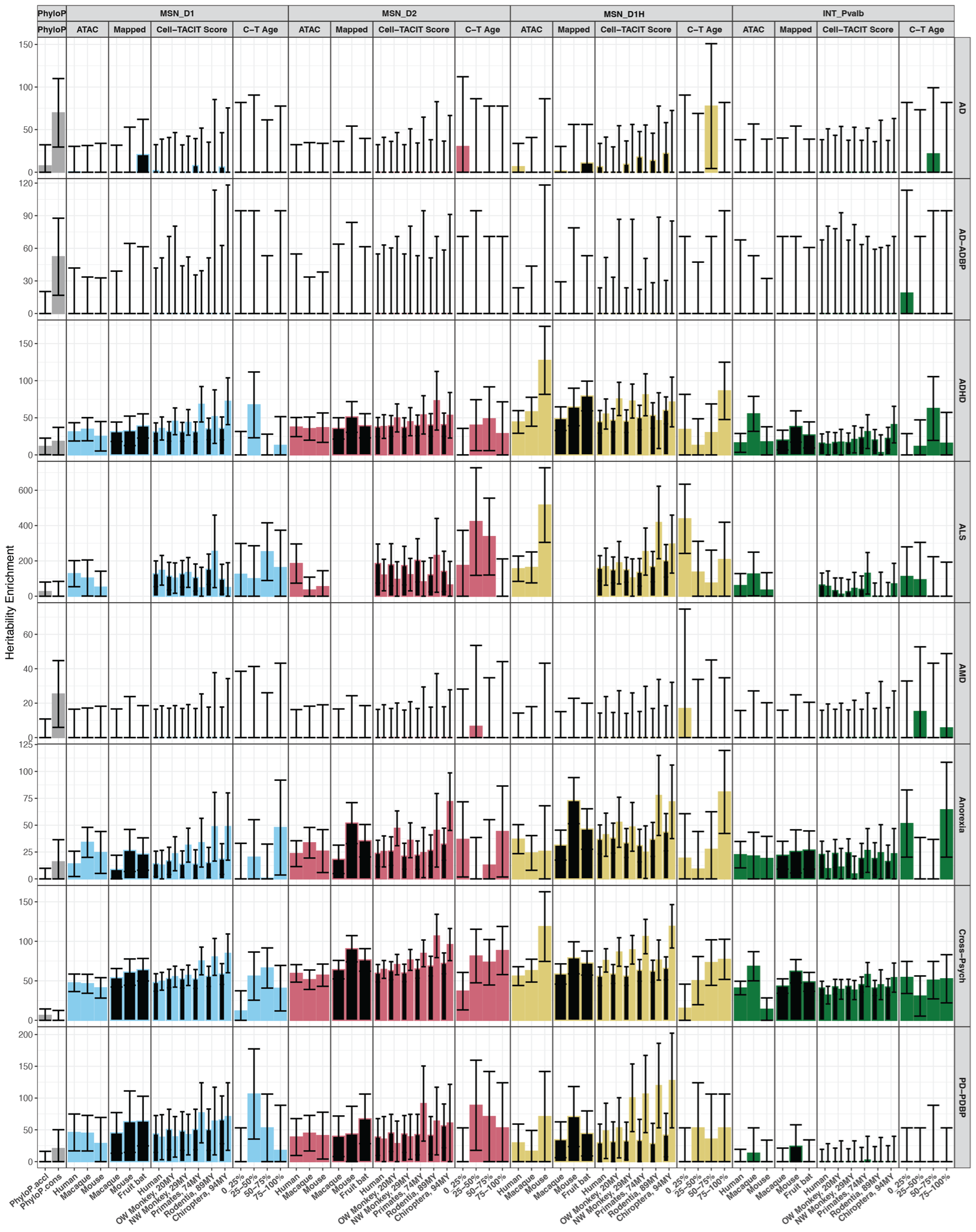

#### Figure S11. Heritability enrichment of neurodegenerative and psychiatric traits and the conserved neuronal epigenome

Heritability enrichments in D1, D2, and D1H MSNs, Pvalb interneurons cell type OCRs from a number of sources showed cell type OCRs measured or predicted activity in other species relevant to human complex traits. Barplots are the heritiability enrichment from LD score regression analyses with error bars plotting standard error of the enrichment.

phyloP: mammalian phyloP constrained (phyloP.cons) or accelerated regions (phyloP.accel), gray bars.

ATAC-seq: cell type OCRs measured in human, human and macaque, or human and mouse.

CTACIT Score: human cell type OCRs mapabble to distant species (black bars) or mappable and predicted active by CTACIT machine learning models (colored bars). C-T Age: CTACIT Age estimating whether human cell type open chromatin is predicted active in closely or across distantly diverged placental mammals. The top quartile corresponds to open chromatin regions that are “older” in placental mammalian genomes.

CTACIT Score: human cell type OCRs mapabble to distant species (black bars) or mappable and predicted active by CTACIT machine learning models (colored bars).

C-T Age: CTACIT Age estimating whether human cell type open chromatin is predicted active in closely or across distantly diverged placental mammals. The top quartile corresponds to open chromatin regions that are “older” in placental mammalian genomes.

Trait: Alzheimer’s Disease, AD; Alzheimer’s disease or AD-by-proxy (AD-ADBP); ADHD Amyotrophic Lateral Sclerosis (ALS); Age-related macular degeneration (AMD); Parkinson’s Disease or Parkinson’s Disease-by-proxy (PD-PDBP).

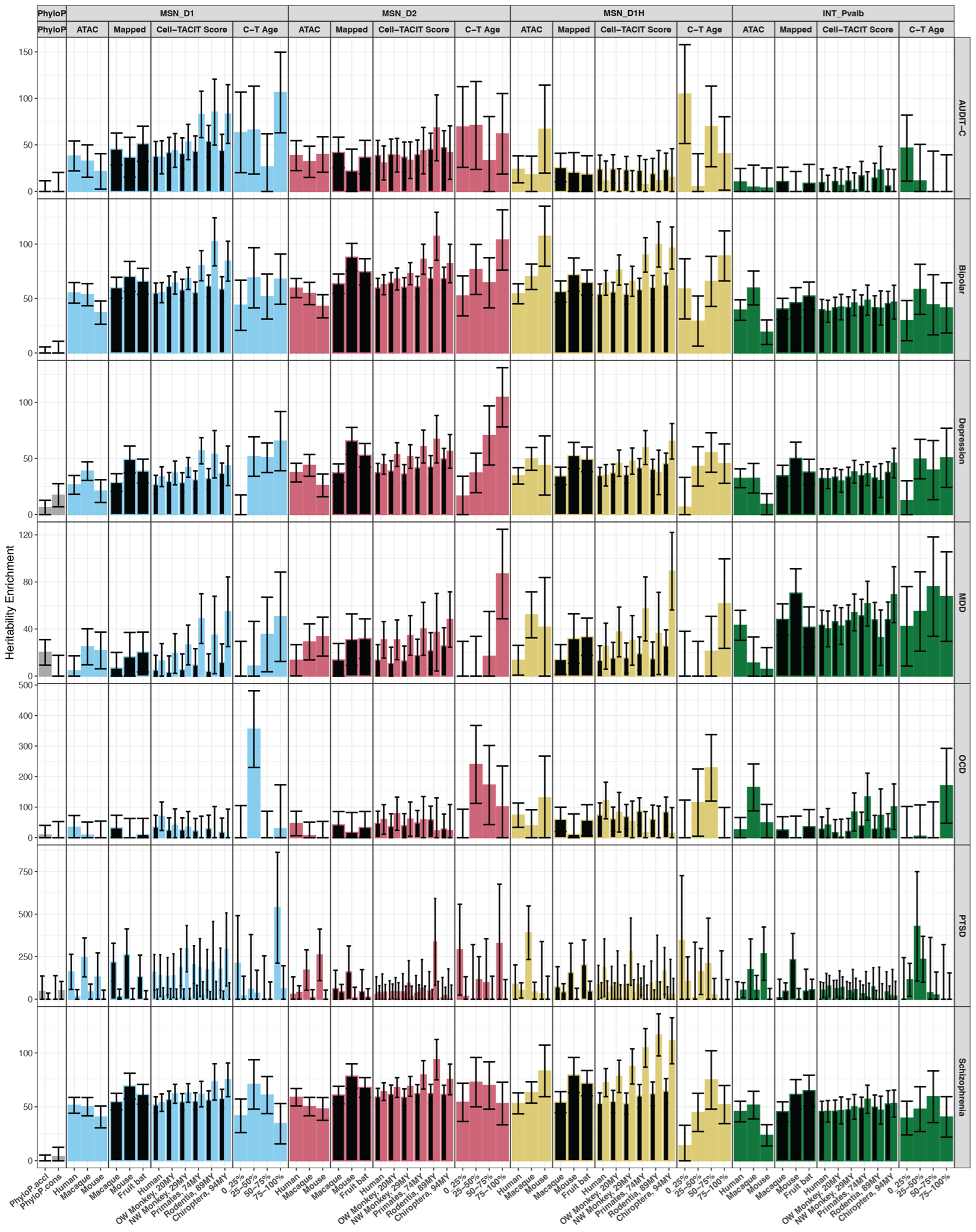

#### Figure S12. Heritability enrichment of substance abuse and psychiatric traits and the conserved neuronal epigenome

Heritability enrichments in D1, D2, and D1H MSNs, Pvalb interneurons cell type OCRs from a number of sources showed cell type OCRs measured or predicted activity in other species relevant to human complex traits. Barplots are the heritiability enrichment from LD score regression analyses with error bars plotting standard error of the enrichment.

phyloP: mammalian phyloP constrained (phyloP.cons) or accelerated regions (phyloP.accel), gray bars.

ATAC-seq: cell type OCRs measured in human, human and macaque, or human and mouse

CTACIT Score: human cell type OCRs mapabble to distant species (black bars) or mappable and predicted active by CTACIT machine learning models (colored bars). C-T Age: CTACIT Age estimating whether human cell type open chromatin is predicted active in closely or across distantly diverged placental mammals. The top quartile corresponds to open chromatin regions that are “older” in placental mammalian genomes.

Trait: Alcohol use-disorder identification test, consumption domain (AUDIT-C); MDD, major depressive disorder; OCD, obsessive-compulsive disorder; PTSD, post-traumatic disorder.

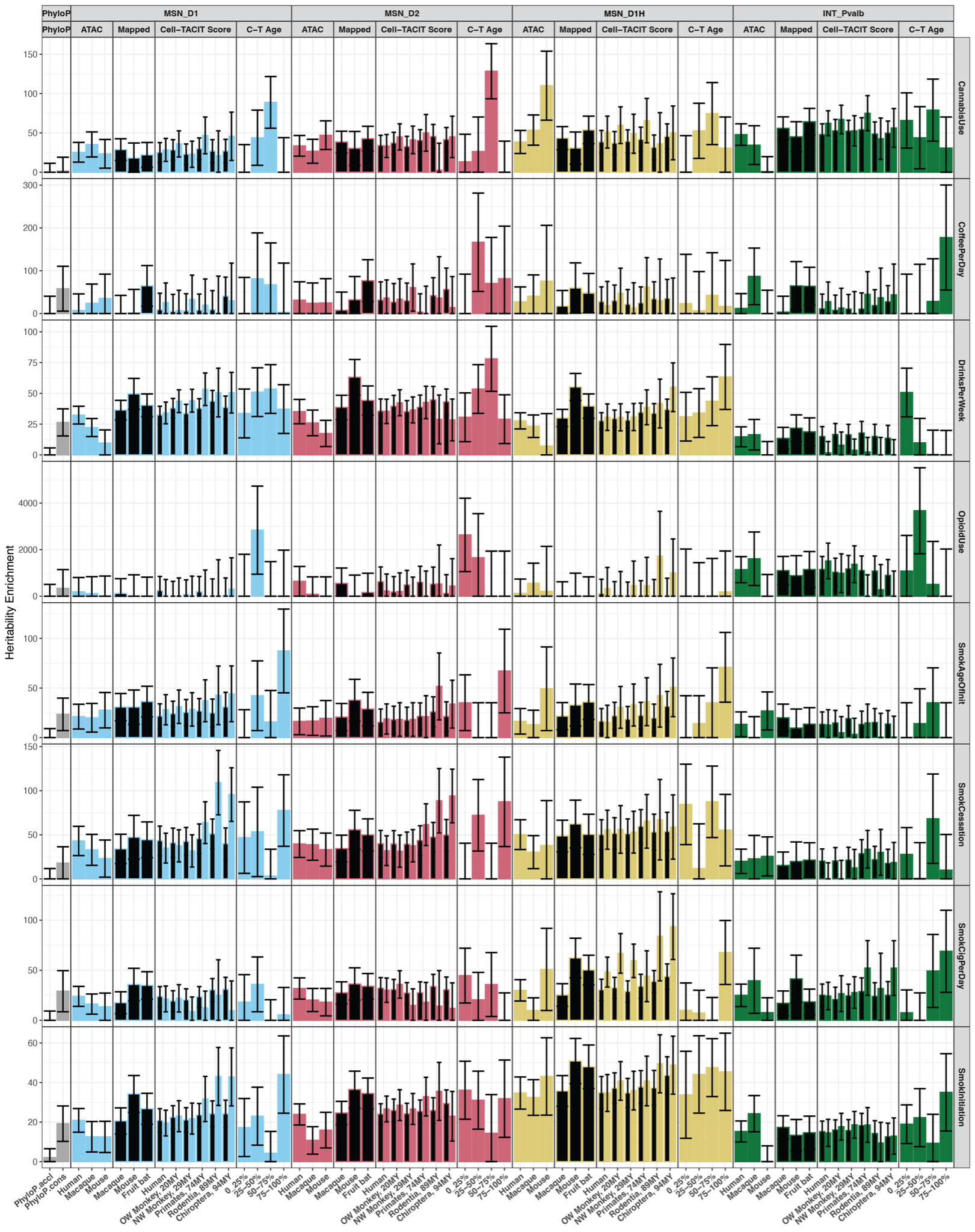

#### Figure S13. Heritability enrichment of substance abuse traits and the conserved neuronal epigenome

Heritability enrichments in D1, D2, and D1H MSNs, Pvalb interneurons cell type OCRs from a number of sources showed cell type OCRs measured or predicted activity in other species relevant to human complex traits. Barplots are the heritiability enrichment from LD score regression analyses with error bars plotting standard error of the enrichment.

phyloP: mammalian phyloP constrained (phyloP.cons) or accelerated regions (phyloP.accel), gray bars.

ATAC-seq: cell type OCRs measured in human, human and macaque, or human and mouse.

CTACIT Score: human cell type OCRs mapabble to distant species (black bars) or mappable and predicted active by CTACIT machine learning models (colored bars). C-T Age: CTACIT Age estimating whether human cell type open chromatin is predicted active in closely or across distantly diverged placental mammals. The top quartile corresponds to open chromatin regions that are “older” in placental mammalian genomes.

Trait: age of smoking initiation (SmokAgeInit); previous smoker vs. current smoker (SmokCessation); current smoker vs. non-smoker (SmokInitiation).

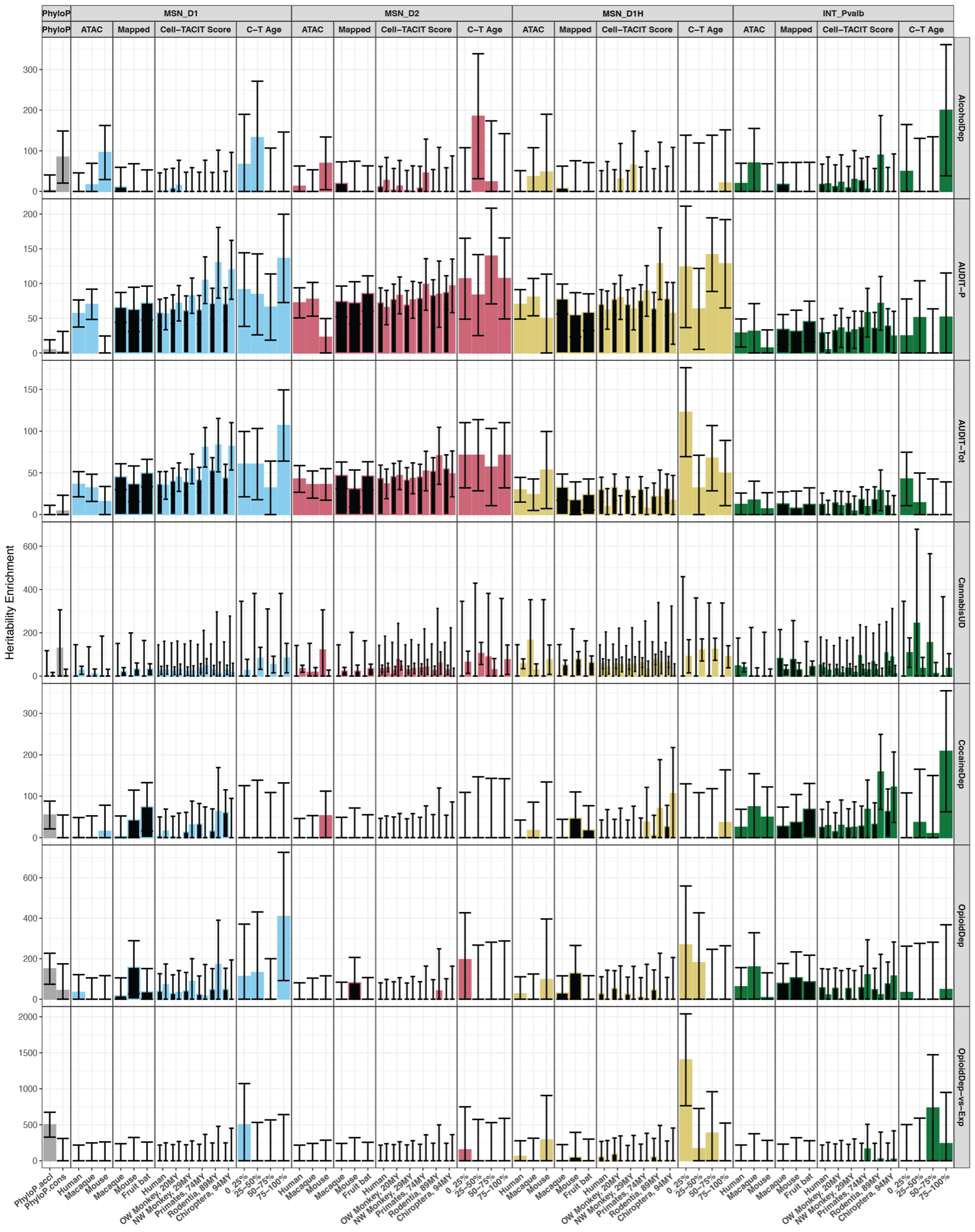

#### Figure S14. Heritability enrichment of substance use disorder traits and the conserved neuronal epigenome

Heritability enrichments in D1, D2, and D1H MSNs, Pvalb interneurons cell type OCRs from a number of sources showed cell type OCRs measured or predicted activity in other species relevant to human complex traits. Barplots are the heritiability enrichment from LD score regression analyses with error bars plotting standard error of the enrichment.

phyloP: mammalian phyloP constrained (phyloP.cons) or accelerated regions (phyloP.accel), gray bars.

ATAC-seq: cell type OCRs measured in human, human and macaque, or human and mouse.

CTACIT Score: human cell type OCRs mapabble to distant species (black bars) or mappable and predicted active by CTACIT machine learning models (colored bars). C-T Age: CTACIT Age estimating whether human cell type open chromatin is predicted active in closely or across distantly diverged placental mammals. The top quartile corresponds to open chromatin regions that are “older” in placental mammalian genomes.

Trait: Alcohol dependence (AlcDependence); alcohol use disorder identification test, problematic use domain (AUDIT-P); alcohol use disorder identification test, total score (AUDIT-Tot); cannabis use disorder (CannabisUD); Cocaine dependence (CocaineDep); Opioid dependence (OpioidDep); Opioid dependence vs. opioid exposure (OpioidDep-vs-Exp).

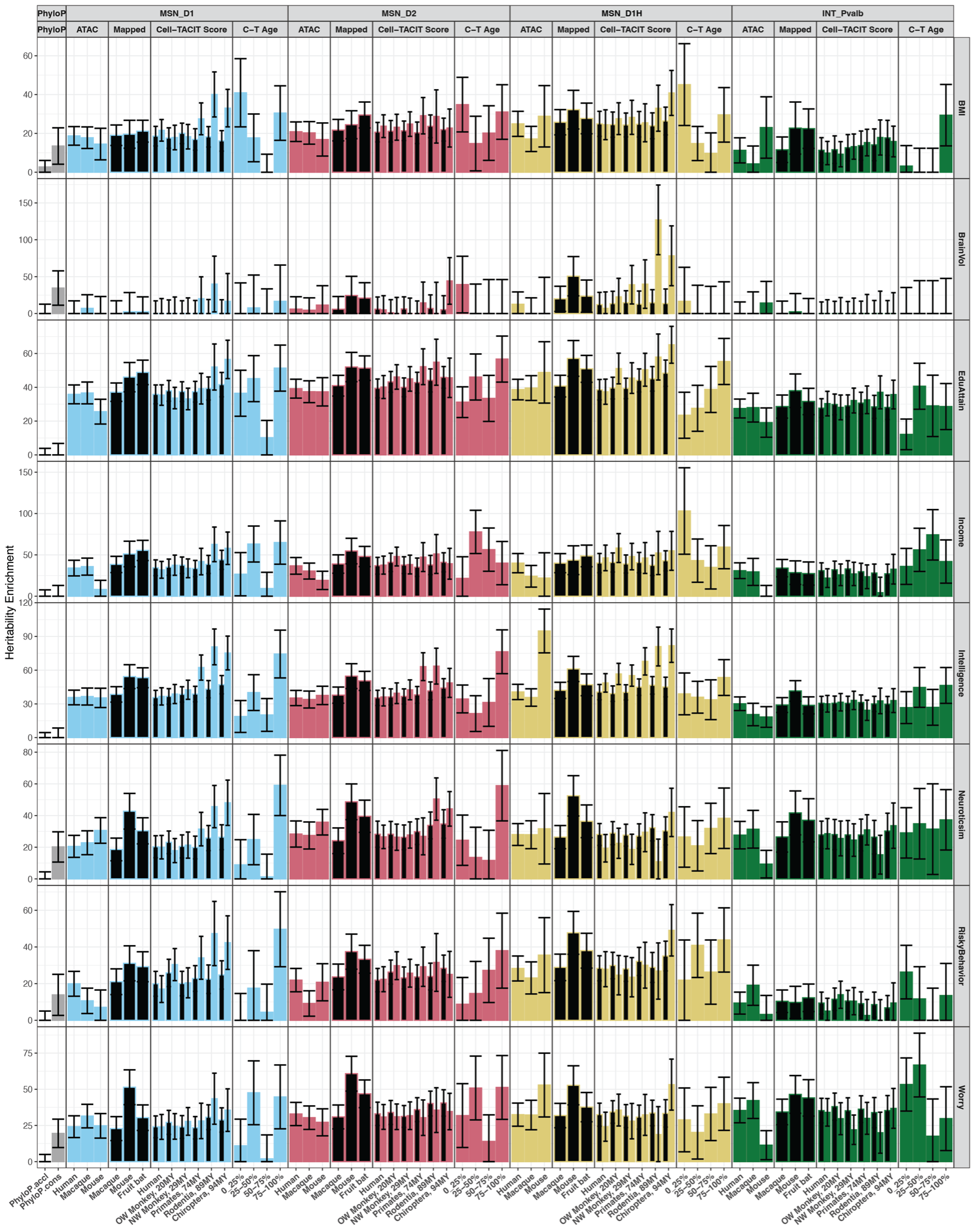

#### Figure S15. Heritability enrichment of neurological traits and the conserved neuronal epigenome

Heritability enrichments in D1, D2, and D1H MSNs, Pvalb interneurons cell type OCRs from a number of sources showed cell type OCRs measured or predicted activity in other species relevant to human complex traits. Barplots are the heritiability enrichment from LD score regression analyses with error bars plotting standard error of the enrichment.

phyloP: mammalian phyloP constrained (phyloP.cons) or accelerated regions (phyloP.accel), gray bars.

ATAC-seq: cell type OCRs measured in human, human and macaque, or human and mouse.

CTACIT Score: human cell type OCRs mapabble to distant species (black bars) or mappable and predicted active by CTACIT machine learning models (colored bars). C-T Age: CTACIT Age estimating whether human cell type open chromatin is predicted active in closely or across distantly diverged placental mammals. The top quartile corresponds to open chromatin regions that are “older” in placental mammalian genomes.

Trait: Body-mass index (BMI), brain volume (BrainVol), educational attainment (EduAttain).

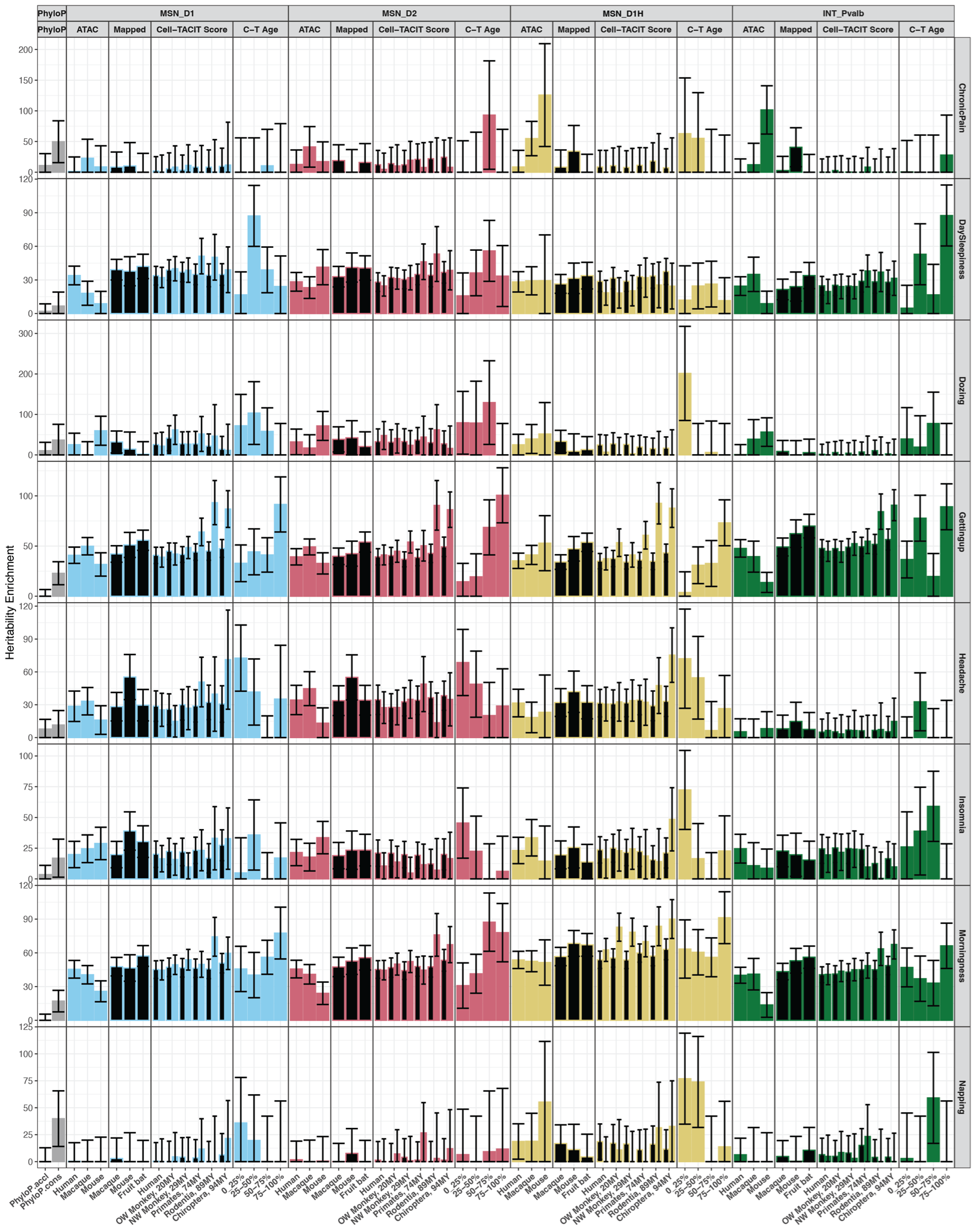

#### Figure S16. Heritability enrichment of chronic pain and sleep-related traits and the conserved neuronal epigenome

Heritability enrichments in D1, D2, and D1H MSNs, Pvalb interneurons cell type OCRs from a number of sources showed cell type OCRs measured or predicted activity in other species relevant to human complex traits. Barplots are the heritiability enrichment from LD score regression analyses with error bars plotting standard error of the enrichment.

phyloP: mammalian phyloP constrained (phyloP.cons) or accelerated regions (phyloP.accel), gray bars.

ATAC-seq: cell type OCRs measured in human, human and macaque, or human and mouse.

CTACIT Score: human cell type OCRs mapabble to distant species (black bars) or mappable and predicted active by CTACIT machine learning models (colored bars). C-T Age: CTACIT Age estimating whether human cell type open chromatin is predicted active in closely or across distantly diverged placental mammals. The top quartile corresponds to open chromatin regions that are “older” in placental mammalian genomes.

Trait: multi-site chronic pain (ChronicPain), daytime sleepiness (DaySleepiness), ease of getting up in the morning (GettingUp), preference for morning vs. evening (morningness).

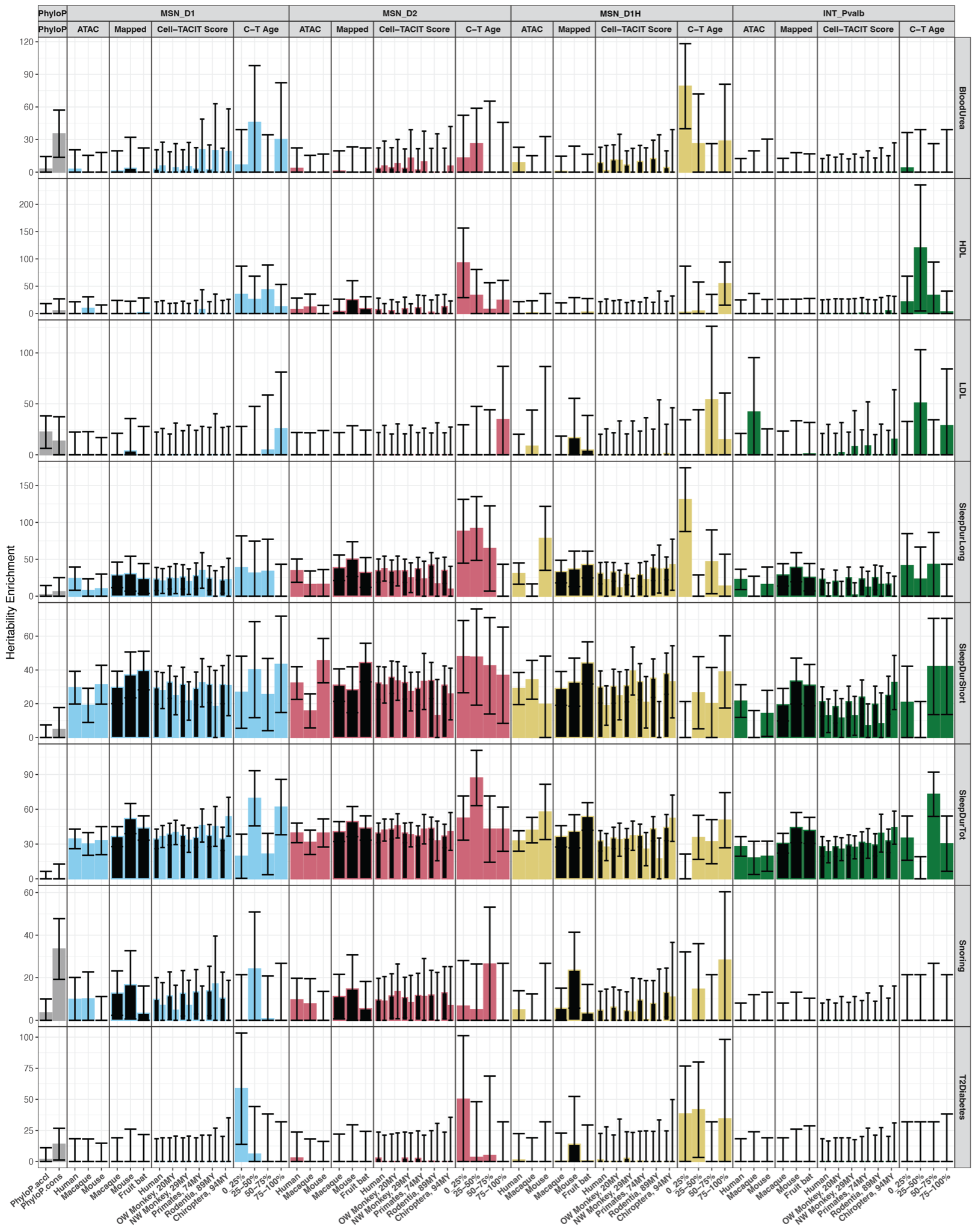

#### Figure S17. Heritability enrichment of metabolic and sleep-related traits and the conserved neuronal epigenome

Heritability enrichments in D1, D2, and D1H MSNs, Pvalb interneurons cell type OCRs from a number of sources showed cell type OCRs measured or predicted activity in other species relevant to human complex traits. Barplots are the heritiability enrichment from LD score regression analyses with error bars plotting standard error of the enrichment.

phyloP: mammalian phyloP constrained (phyloP.cons) or accelerated regions (phyloP.accel), gray bars.

ATAC-seq: cell type OCRs measured in human, human and macaque, or human and mouse.

CTACIT Score: human cell type OCRs mapabble to distant species (black bars) or mappable and predicted active by CTACIT machine learning models (colored bars). C-T Age: CTACIT Age estimating whether human cell type open chromatin is predicted active in closely or across distantly diverged placental mammals. The top quartile corresponds to open chromatin regions that are “older” in placental mammalian genomes.

Trait: high-density lipoprotein (HDL), low-density lipoprotein (LDL), long sleep duration (SleepDurLong), short sleep duration (SleepDurShort), total hours of sleep (SleepDurTot), type 2 diabetes mellitus (T2Diabetes).

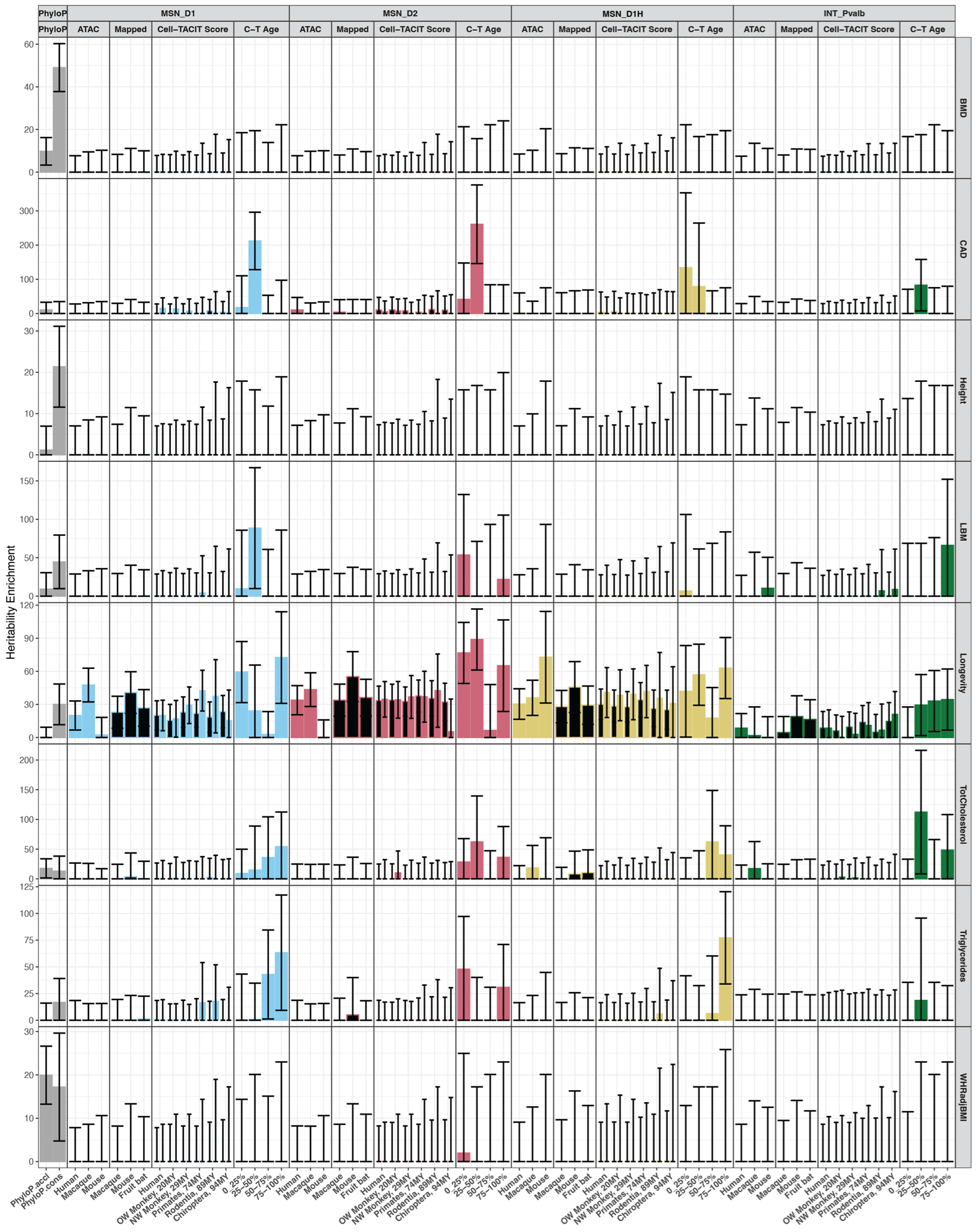

#### Figure S18. Heritability enrichment of anthropomorphic and sleep-related traits and the conserved neuronal epigenome

Heritability enrichments in D1, D2, and D1H MSNs, Pvalb interneurons cell type OCRs from a number of sources showed cell type OCRs measured or predicted activity in other species relevant to human complex traits. Barplots are the heritiability enrichment from LD score regression analyses with error bars plotting standard error of the enrichment.

phyloP: mammalian phyloP constrained (phyloP.cons) or accelerated regions (phyloP.accel), gray bars.

ATAC-seq: cell type OCRs measured in human, human and macaque, or human and mouse.

CTACIT Score: human cell type OCRs mapabble to distant species (black bars) or mappable and predicted active by CTACIT machine learning models (colored bars). C-T Age: CTACIT Age estimating whether human cell type open chromatin is predicted active in closely or across distantly diverged placental mammals. The top quartile corresponds to open chromatin regions that are “older” in placental mammalian genomes.

Trait: heel T-score bone mineral density (BMD), coronary artery disease (CAD), lean body mass (LBM), parental longeivity (Longevity), total cholesterol (TotCholesterol), waist-hip ratio adjusted for body mass index (WHRadjBMI).

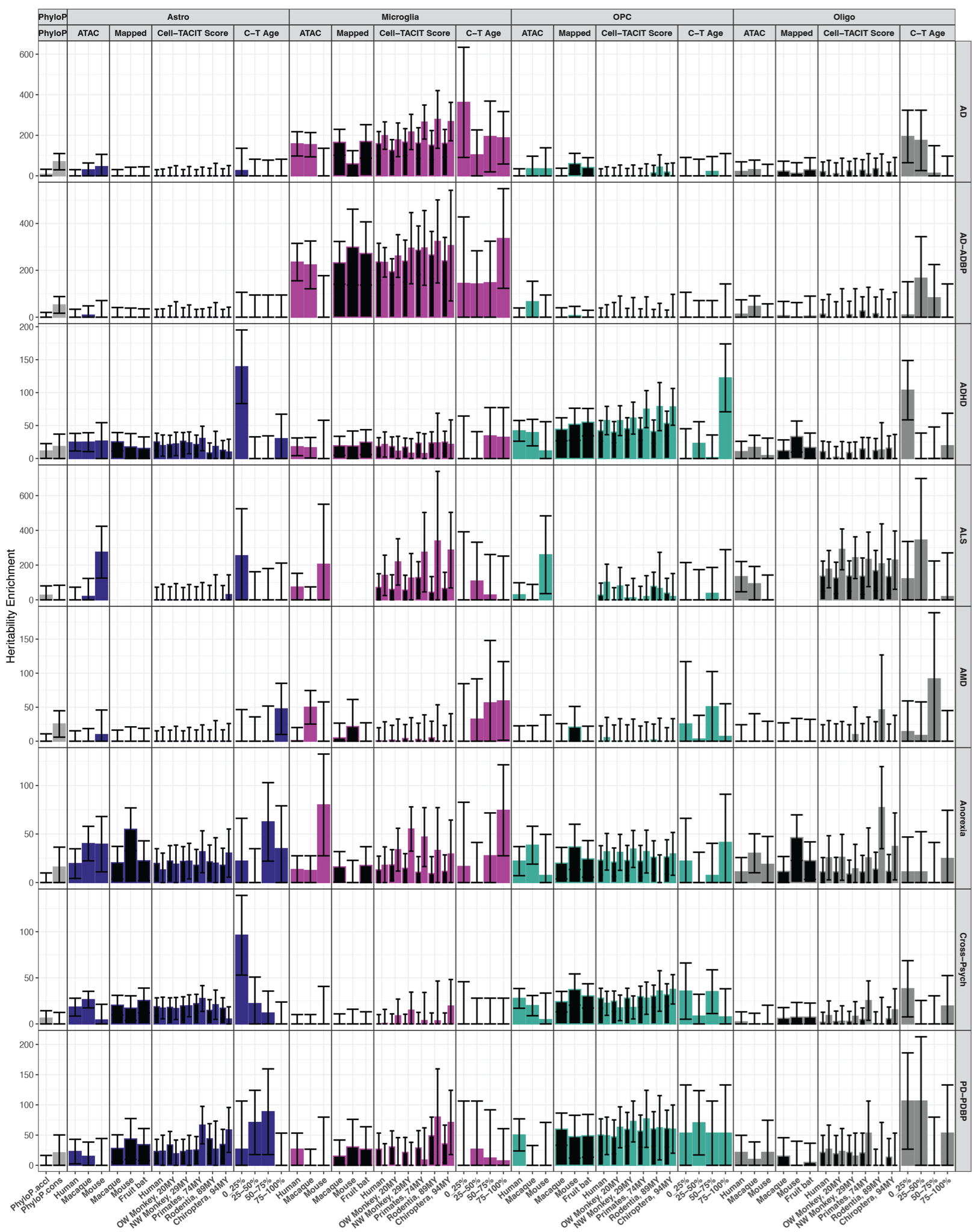

#### Figure S19. Heritability enrichment of neurodegenerative and psychiatric traits and the conserved glia and liver epigenome

Heritability enrichments in astrocyte, microglia, oligodendrocyte precursor cells, oligodendrocyte, and liver cell type OCRs from a number of sources showed cell type OCRs measured or predicted activity in other species relevant to human complex traits. Barplots are the heritiability enrichment from LD score regression analyses with error bars plotting standard error of the enrichment.

phyloP: mammalian phyloP constrained (phyloP.cons) or accelerated regions (phyloP.accel), gray bars.

ATAC-seq: cell type OCRs measured in human, human and macaque, or human and mouse.

CTACIT Score: human cell type OCRs mapabble to distant species (black bars) or mappable and predicted active by CTACIT machine learning models (colored bars). C-T Age: CTACIT Age estimating whether human cell type open chromatin is predicted active in closely or across distantly diverged placental mammals. The top quartile corresponds to open chromatin regions that are “older” in placental mammalian genomes.

Trait: Alzheimer’s Disease, AD; Alzheimer’s disease or AD-by-proxy (AD-ADBP); ADHD Amyotrophic Lateral Sclerosis (ALS); Age-related macular degeneration (AMD); Parkinson’s Disease or Parkinson’s Disease-by-proxy (PD-PDBP).

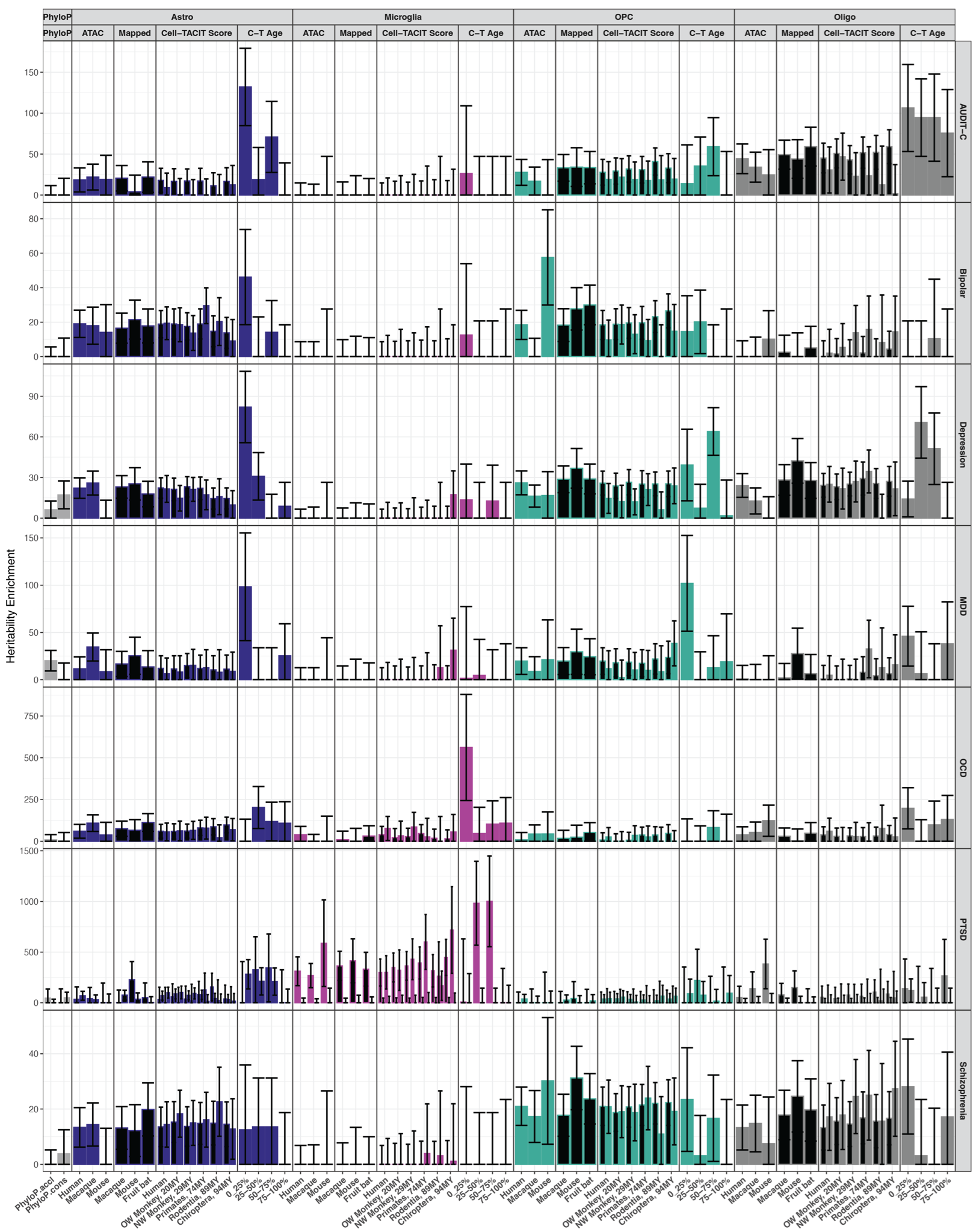

#### Figure S20. Heritability enrichment of substance use and psychiatric traits and the conserved glia and liver epigenome

Heritability enrichments in astrocyte, microglia, oligodendrocyte precursor cells, oligodendrocyte, and liver cell type OCRs from a number of sources showed cell type OCRs measured or predicted activity in other species relevant to human complex traits. Barplots are the heritiability enrichment from LD score regression analyses with error bars plotting standard error of the enrichment.

phyloP: mammalian phyloP constrained (phyloP.cons) or accelerated regions (phyloP.accel), gray bars.

ATAC-seq: cell type OCRs measured in human, human and macaque, or human and mouse.

CTACIT Score: human cell type OCRs mapabble to distant species (black bars) or mappable and predicted active by CTACIT machine learning models (colored bars). C-T Age: CTACIT Age estimating whether human cell type open chromatin is predicted active in closely or across distantly diverged placental mammals. The top quartile corresponds to open chromatin regions that are “older” in placental mammalian genomes.

Trait: Alcohol use-disorder identification test, consumption domain (AUDIT-C); major depressive disorder (MDD); obsessive-compulsive disorder (OCD); post-traumatic disorder (PTSD).

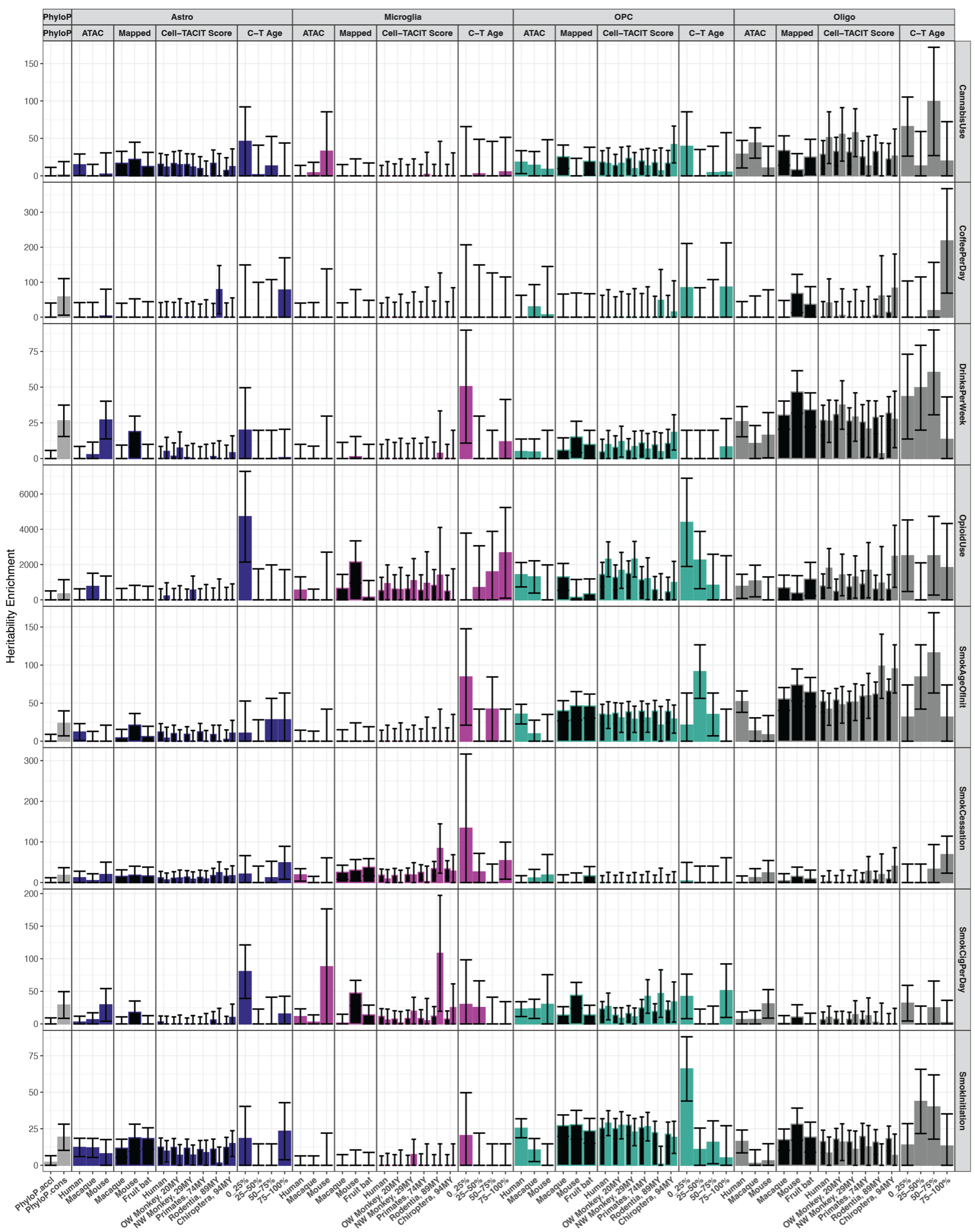

#### Figure S21. Heritability enrichment of substance use traits and the conserved glia and liver epigenome

Heritability enrichments in astrocyte, microglia, oligodendrocyte precursor cells, oligodendrocyte, and liver cell type OCRs from a number of sources showed cell type OCRs measured or predicted activity in other species relevant to human complex traits. Barplots are the heritiability enrichment from LD score regression analyses with error bars plotting standard error of the enrichment.

phyloP: mammalian phyloP constrained (phyloP.cons) or accelerated regions (phyloP.accel), gray bars.

ATAC-seq: cell type OCRs measured in human, human and macaque, or human and mouse

CTACIT Score: human cell type OCRs mapabble to distant species (black bars) or mappable and predicted active by CTACIT machine learning models (colored bars). C-T Age: CTACIT Age estimating whether human cell type open chromatin is predicted active in closely or across distantly diverged placental mammals. The top quartile corresponds to open chromatin regions that are “older” in placental mammalian genomes.

Trait: age of smoking initiation (SmokAgeInit); previous smoker vs. current smoker (SmokCessation); current smoker vs. non-smoker (SmokInitiation).

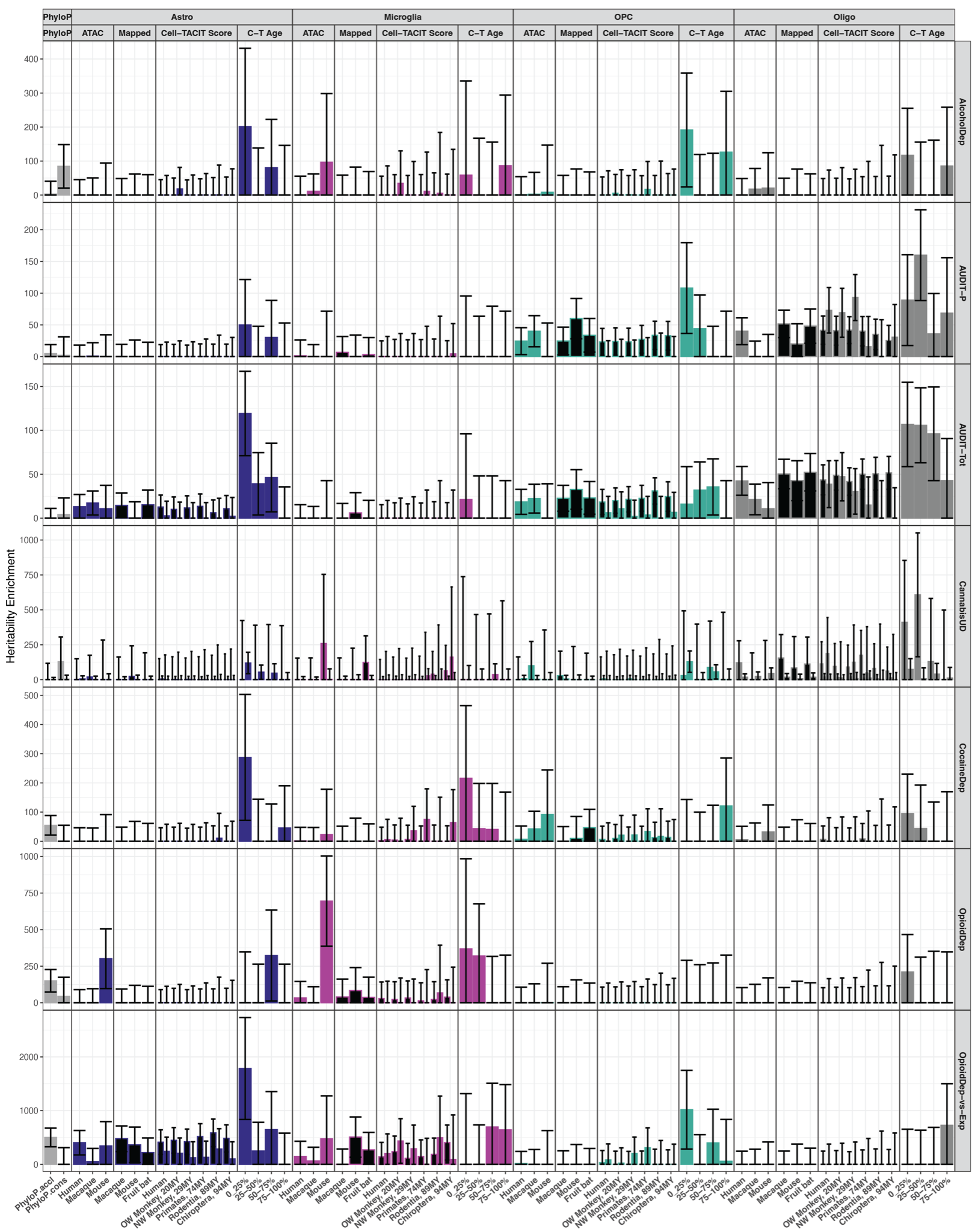

#### Figure S22. Heritability enrichment of substance use disorder traits and the conserved glia and liver epigenome

Heritability enrichments in astrocyte, microglia, oligodendrocyte precursor cells, oligodendrocyte, and liver cell type OCRs from a number of sources showed cell type OCRs measured or predicted activity in other species relevant to human complex traits. Barplots are the heritiability enrichment from LD score regression analyses with error bars plotting standard error of the enrichment.

phyloP: mammalian phyloP constrained (phyloP.cons) or accelerated regions (phyloP.accel), gray bars.

ATAC-seq: cell type OCRs measured in human, human and macaque, or human and mouse.

CTACIT Score: human cell type OCRs mapabble to distant species (black bars) or mappable and predicted active by CTACIT machine learning models (colored bars). C-T Age: CTACIT Age estimating whether human cell type open chromatin is predicted active in closely or across distantly diverged placental mammals. The top quartile corresponds to open chromatin regions that are “older” in placental mammalian genomes.

Trait: Alcohol dependence (AlcDependence); alcohol use disorder identification test, problematic use domain (AUDIT-P); alcohol use disorder identification test, total score (AUDIT-Tot); cannabis use disorder (CannabisUD); Cocaine dependence (CocaineDep); Opioid dependence (OpioidDep); Opioid dependence vs. opioid exposure (OpioidDep-vs-Exp).

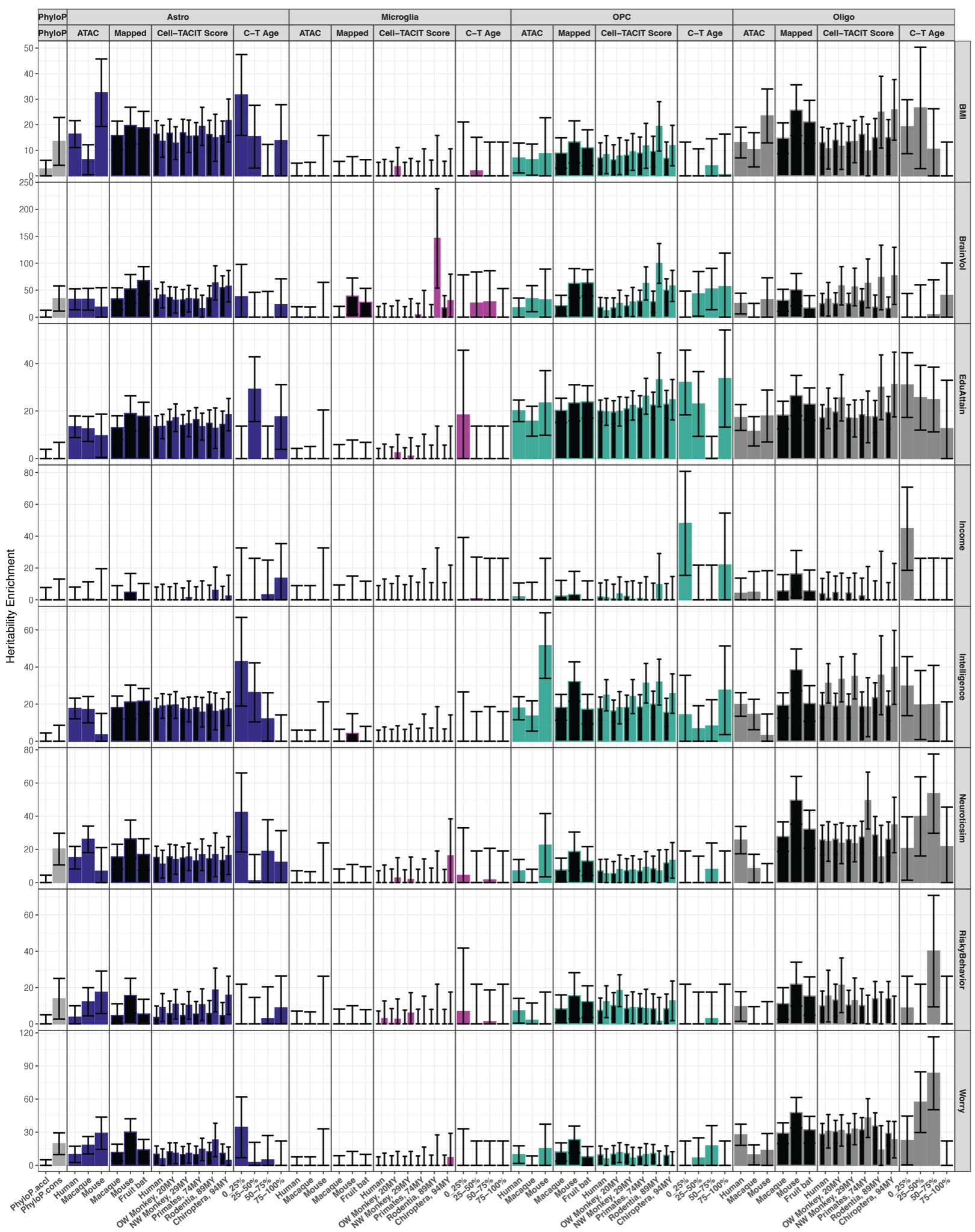

#### Figure S23. Heritability enrichment of neurological traits and the conserved glia and liver epigenome

Heritability enrichments in astrocyte, microglia, oligodendrocyte precursor cells, oligodendrocyte, and liver cell type OCRs from a number of sources showed cell type OCRs measured or predicted activity in other species relevant to human complex traits. Barplots are the heritiability enrichment from LD score regression analyses with error bars plotting standard error of the enrichment.

phyloP: mammalian phyloP constrained (phyloP.cons) or accelerated regions (phyloP.accel), gray bars.

ATAC-seq: cell type OCRs measured in human, human and macaque, or human and mouse.

CTACIT Score: human cell type OCRs mapabble to distant species (black bars) or mappable and predicted active by CTACIT machine learning models (colored bars). C-T Age: CTACIT Age estimating whether human cell type open chromatin is predicted active in closely or across distantly diverged placental mammals. The top quartile corresponds to open chromatin regions that are “older” in placental mammalian genomes.

Trait: Body-mass index (BMI), brain volume (BrainVol), educational attainment (EduAttain).

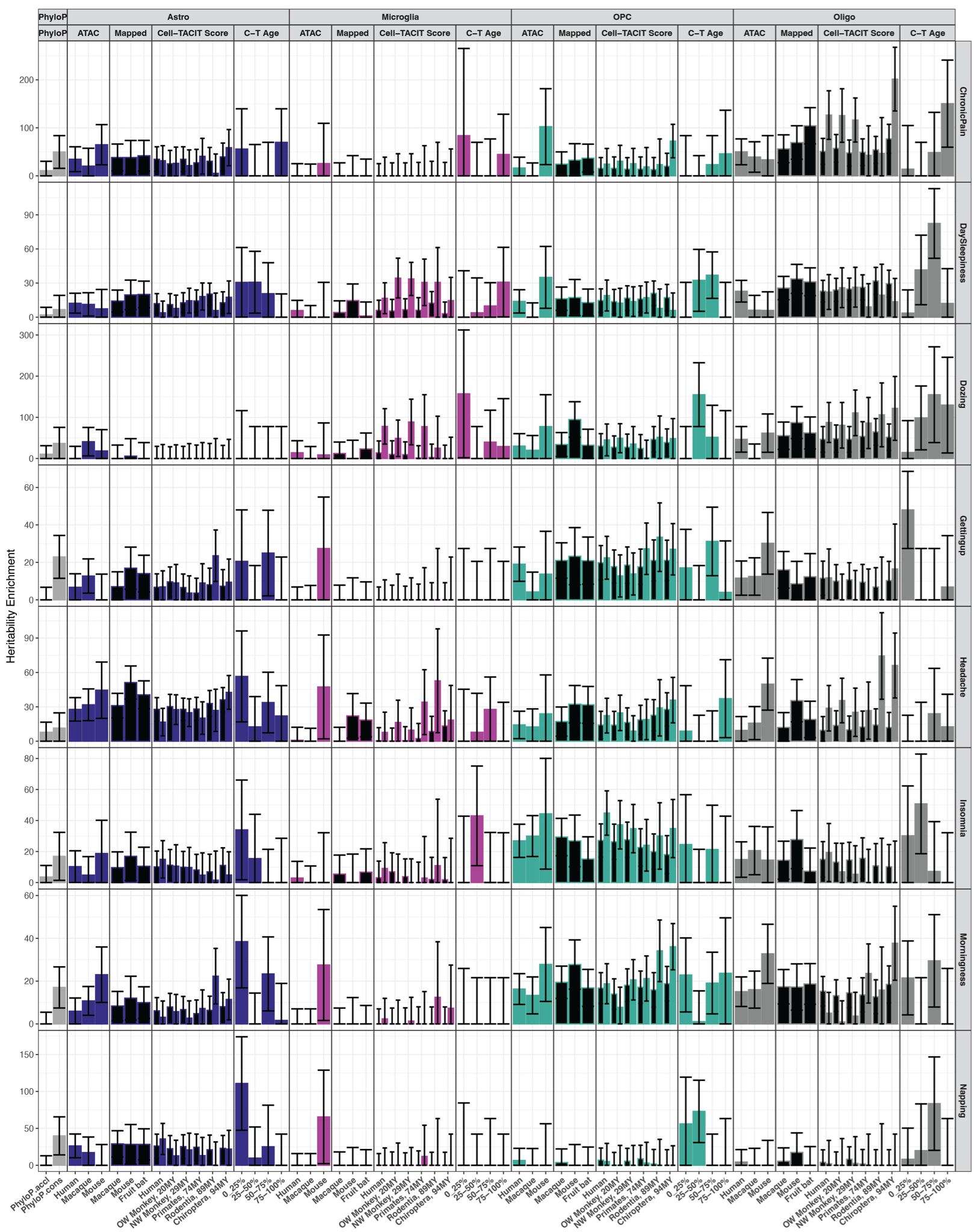

#### Figure S24. Heritability enrichment of chronic pain and sleep-related traits and the conserved glia and liver epigenome

Heritability enrichments in astrocyte, microglia, oligodendrocyte precursor cells, oligodendrocyte, and liver cell type OCRs from a number of sources showed cell type OCRs measured or predicted activity in other species relevant to human complex traits. Barplots are the heritiability enrichment from LD score regression analyses with error bars plotting standard error of the enrichment.

phyloP: mammalian phyloP constrained (phyloP.cons) or accelerated regions (phyloP.accel), gray bars.

ATAC-seq: cell type OCRs measured in human, human and macaque, or human and mouse.

CTACIT Score: human cell type OCRs mapabble to distant species (black bars) or mappable and predicted active by CTACIT machine learning models (colored bars). C-T Age: CTACIT Age estimating whether human cell type open chromatin is predicted active in closely or across distantly diverged placental mammals. The top quartile corresponds to open chromatin regions that are “older” in placental mammalian genomes.

Trait: multi-site chronic pain (ChronicPain), daytime sleepiness (DaySleepiness), ease of getting up in the morning (GettingUp), preference for morning vs. evening (morningness).

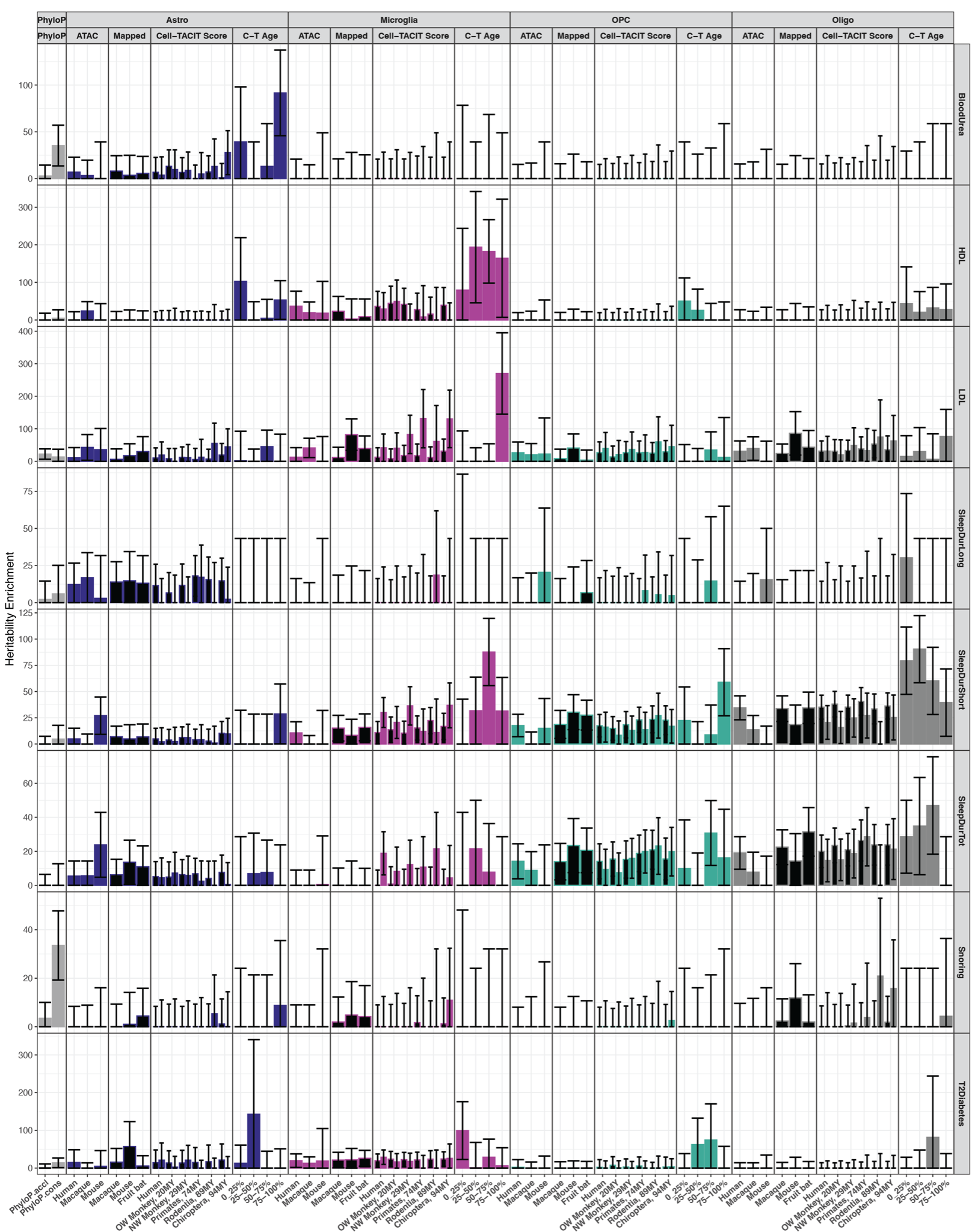

#### Figure S25. Heritability enrichment of metabolic and sleep-related traits and the conserved glia and liver epigenome

Heritability enrichments in astrocyte, microglia, oligodendrocyte precursor cells, oligodendrocyte, and liver cell type OCRs from a number of sources showed cell type OCRs measured or predicted activity in other species relevant to human complex traits. Barplots are the heritiability enrichment from LD score regression analyses with error bars plotting standard error of the enrichment.

phyloP: mammalian phyloP constrained (phyloP.cons) or accelerated regions (phyloP.accel), gray bars.

ATAC-seq: cell type OCRs measured in human, human and macaque, or human and mouse.

CTACIT Score: human cell type OCRs mapabble to distant species (black bars) or mappable and predicted active by CTACIT machine learning models (colored bars). C-T Age: CTACIT Age estimating whether human cell type open chromatin is predicted active in closely or across distantly diverged placental mammals. The top quartile corresponds to open chromatin regions that are “older” in placental mammalian genomes.

Trait: high-density lipoprotein (HDL), low-density lipoprotein (LDL), long sleep duration (SleepDurLong), short sleep duration (SleepDurShort), total hours of sleep (SleepDurTot), type 2 diabetes mellitus (T2Diabetes).

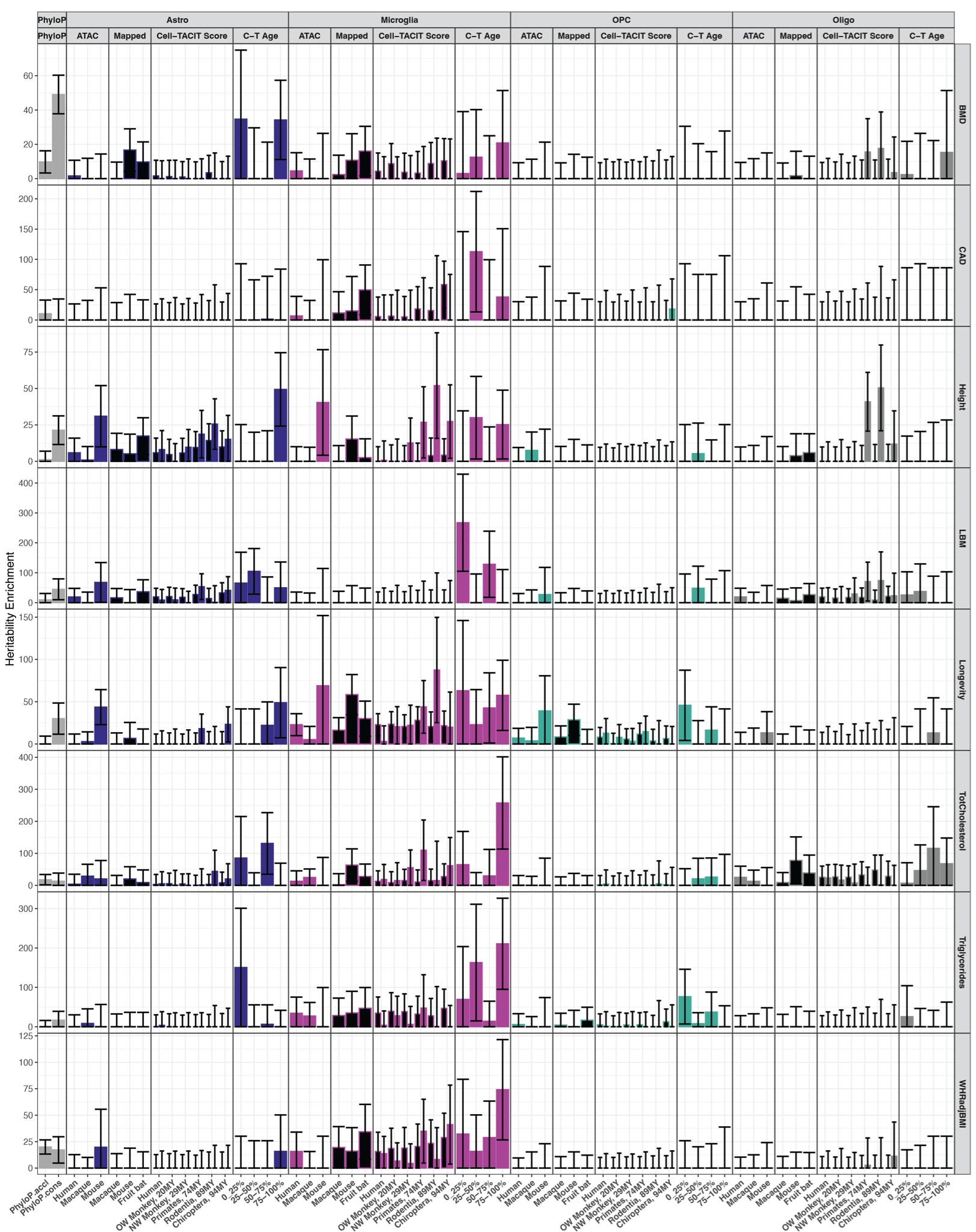

#### Figure S26. Heritability enrichment of anthropomorphic and sleep-related traits and the conserved glia and liver epigenome

Heritability enrichments in astrocyte, microglia, oligodendrocyte precursor cells, oligodendrocyte, and liver cell type OCRs from a number of sources showed cell type OCRs measured or predicted activity in other species relevant to human complex traits. Barplots are the heritiability enrichment from LD score regression analyses with error bars plotting standard error of the enrichment.

phyloP: mammalian phyloP constrained (phyloP.cons) or accelerated regions (phyloP.accel), gray bars.

ATAC-seq: cell type OCRs measured in human, human and macaque, or human and mouse.

CTACIT Score: human cell type OCRs mapabble to distant species (black bars) or mappable and predicted active by CTACIT machine learning models (colored bars). C-T Age: CTACIT Age estimating whether human cell type open chromatin is predicted active in closely or across distantly diverged placental mammals. The top quartile corresponds to open chromatin regions that are “older” in placental mammalian genomes.

Trait: heel T-score bone mineral density (BMD), coronary artery disease (CAD), lean body mass (LBM), parental longevity (Longevity), total cholesterol (TotCholesterol), waist-hip ratio adjusted for body mass index (WHRadjBMI).

From top to bottom, the tracks show human genes, genome recombination rate genetic maps in centiMorgan per million bases (cM/Mb) from the DECODE project [121], Zoonomia primate PhastCons scores, Zoonomia mammals positive phyloP scores [122], fine-mapped SNP lollipop plots, cell type OCR tracks, and corresponding CTACIT cell type OCR heatmaps.

Lollipop tracks of fine-mapped SNPs created with TrackViewer from polyfun fine-mapped SNPs (Methods) [123]. The lollipop feature's positive height reflects the max PIP of the fine-mapped SNPs. Every circle or square is 0.1 units of posterior inclusion probability, PIP, SNPs with less than 0.1 PIP are not plot), and features are jittered to better visualize clusters of fine-mapped SNPs across a large genomic region. The lollipop feature’s negative direction measures how many traits each SNP was mapped with each circle or square is one trait. Lollipop feature outlines are colored by the trait groups the fine-mapped SNPs came from (Methods), light blue: neurological trait, yellow: substance use trait, red: neurodegenerative trait, teal: psychiatric trait.

Cell type OCR tracks: human caudate cell types OCRs with heights reflecting the CTACIT Age of each OCR. MYA: million years ago diverged from humans.

CTACIT Models from MSN_D1: D1 medium spiny neurons, MSN_D2: D2 medium spiny neurons, Microglia. Arrows align cell type OCR track plots to corresponding columns in the heatmaps. Heatmaps display the average CTACIT calibrated scores of species within the same clade/divergence group from humans. OCRs not mappable to a clade are plotted in gray. OCRs labels along the abscissa are truncated to show only the chromosome and start of each 501bp OCR.

(A) Locus around the GRM3 gene with SNPs from the neuroticism GWAS [63,123].

(B) Locus 100kb downstream from the DRD2 gene with SNPs from education attainment, schizophrenia, cigarettes per day, and neuroticism GWAS [52,59,63,124].

(C) Locus around the FAM163B gene locus with SNPs from the SmokingCessation and CigarrettesPerDay GWAS [52].

(D) Locus around the TSPOAP1 gene locus with SNPs from the AlzheimerDisease GWAS and AD & AD-by-Proxy GWAS [37,41].

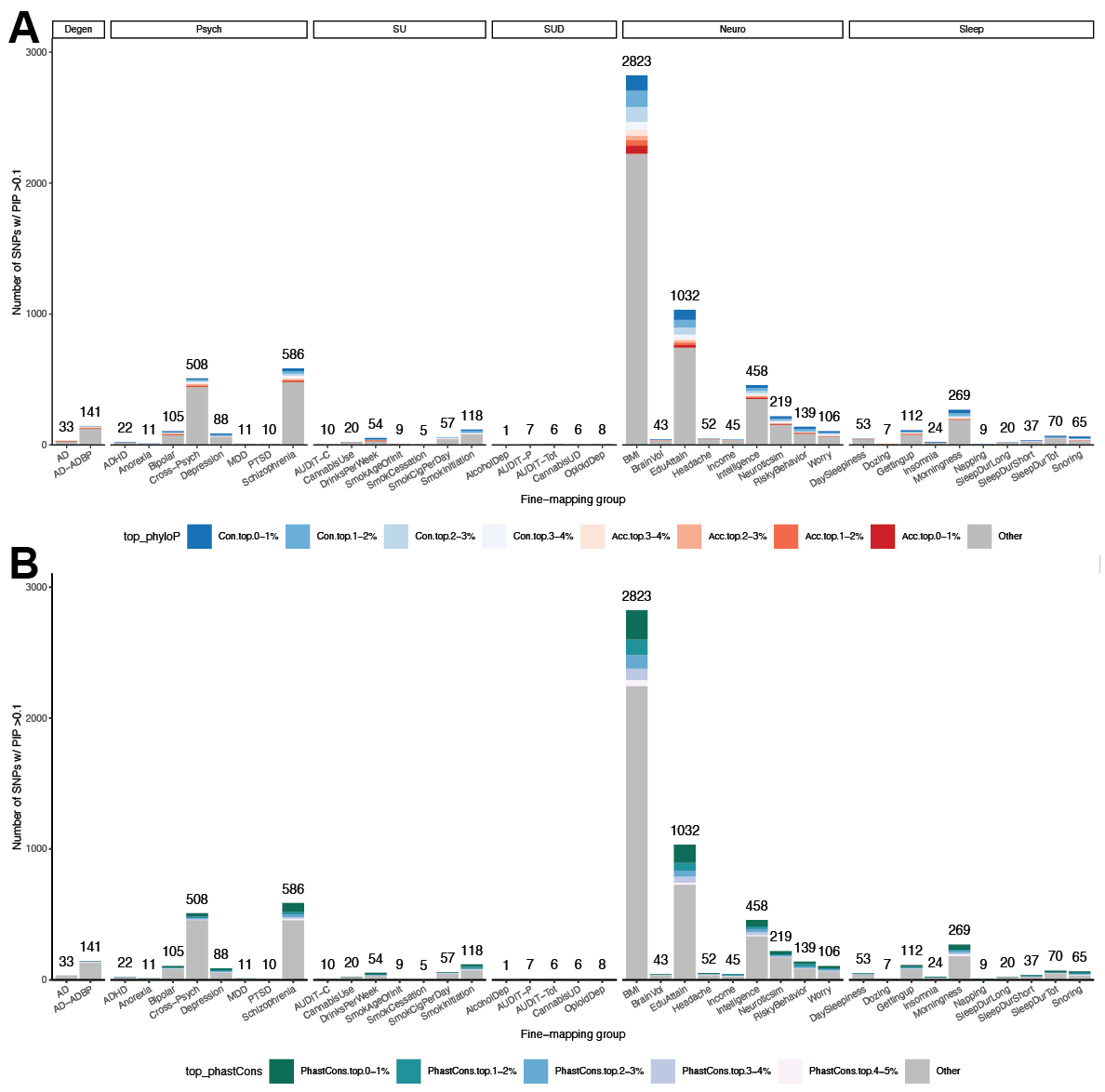

#### Figure S27. Fine-mapping of neuropsychiatric traits show varying number of genetic variants related to power of GWAS to detect genome-wide loci

**(A)** Candidate causal SNPs in neuropsychiatric traits with non-negligible posterior inclusion probabilities (PIP> 0.1). SNPs are annotated by overlap with the top mammalian phyloP accelerated or constrained groups.

**(B)** As in (A) and SNPs are annotated by overlap with top constrained primate Phastcons groups. Neurodegenerative (Degen), psychiatric disorder (Psych), substance use (SU), substance use disorder (SUD), neurological (Neuro), traits as in Data S1.

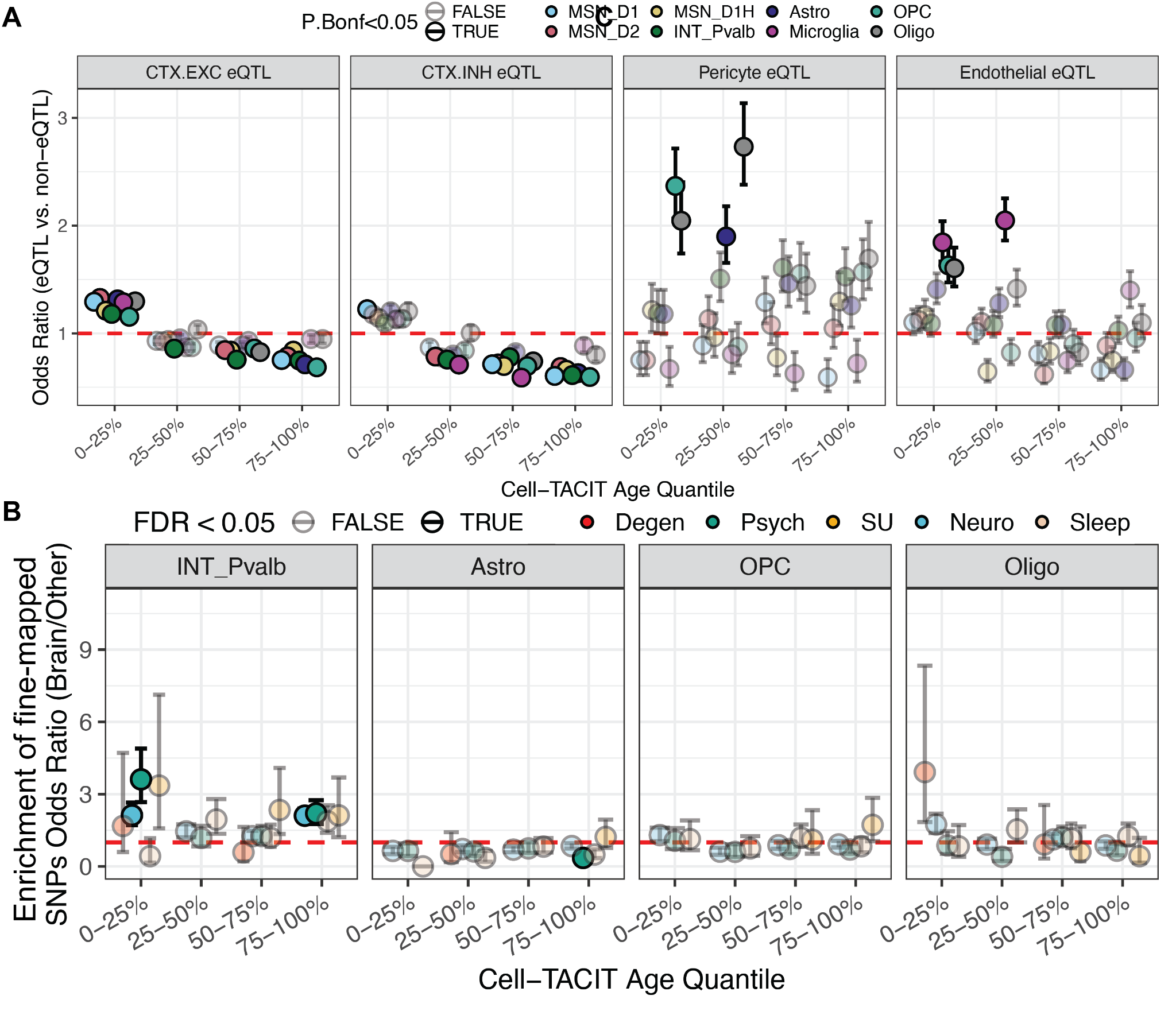

#### Figure S28. Comparison of CTACIT Ages with off-target eQTLs and fine-mapped neuropsychiatric trait genetic variants.

**(A)** Stripchart plotting the odds ratio (±standard error of the mean) that significant eQTLs from single-cell prefrontal cortex cell types with CTACIT Age quartiles. These prefrontal cortex cell types do not have one-to-one correspondence with CTACIT caudate cell types. Cell types with one-to-one correspondence are shown in **Figure 3B.** Each point is colored by the CTACIT model. Solid points are significant enrichment at Bonferroni corrected P-value < 0.05. **(B)** Enrichment stripchart for fine-mapped neuropsychiatric trait SNPs with PIP > 0.10 in CTACIT Age quartiles for the other caudate neural cell types. Each point is colored by enrichment for SNPs fine-mapped in neurodegenerative traits (Degen), psychiatric traits (Psych), substance use traits (SU), neurological traits (Neuro), or sleep traits (Sleep). The top CTACIT cell types are shown in **Figure 3C**. Solid points are significant enrichment at FDR corrected P-value < 0.05.

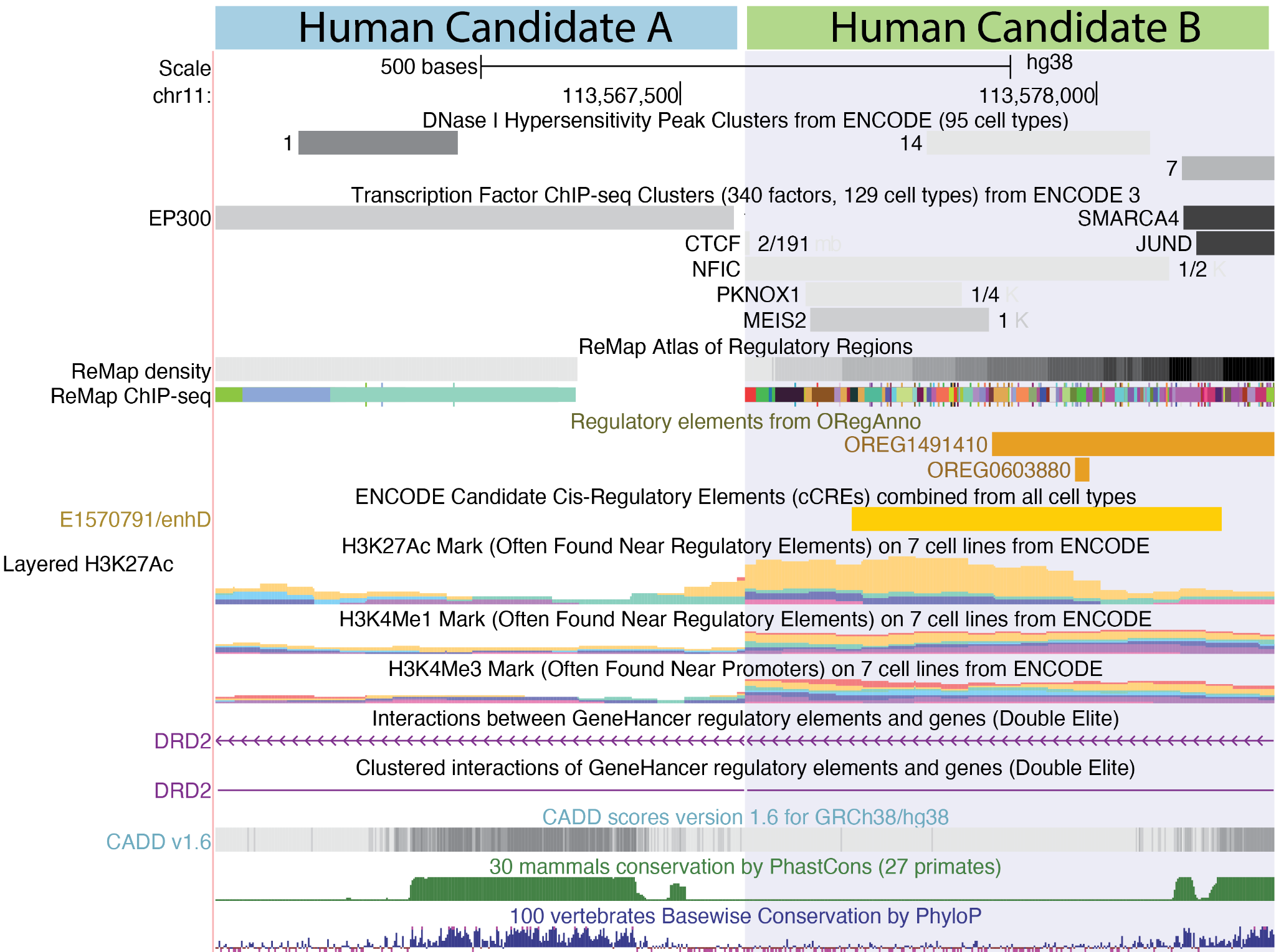

#### Figure S29. UCSC genome tracks for two candidate enhancer sequences upstream of the *DRD2* locus.

Genome tracks of human candidate enhancer A (hg38, chr11:113,567,061-113,567,561) and B (hg38, chr11:113,577,668-113,578,168) with epigenomic tracks available through the UCSC genome browser. The tracks from top to bottom are the ENCODE DNAse I hypersensitivity peaks, ENCODE3 ChIP-seq peaks, ORegAnno database annotations, ENCODE3 cCRE regions, ENCODE histone ChIP-seq multi-layer plots, GeneHancer links, CADD SNP impact predicted scores, and older nucleotide conservation tracks.

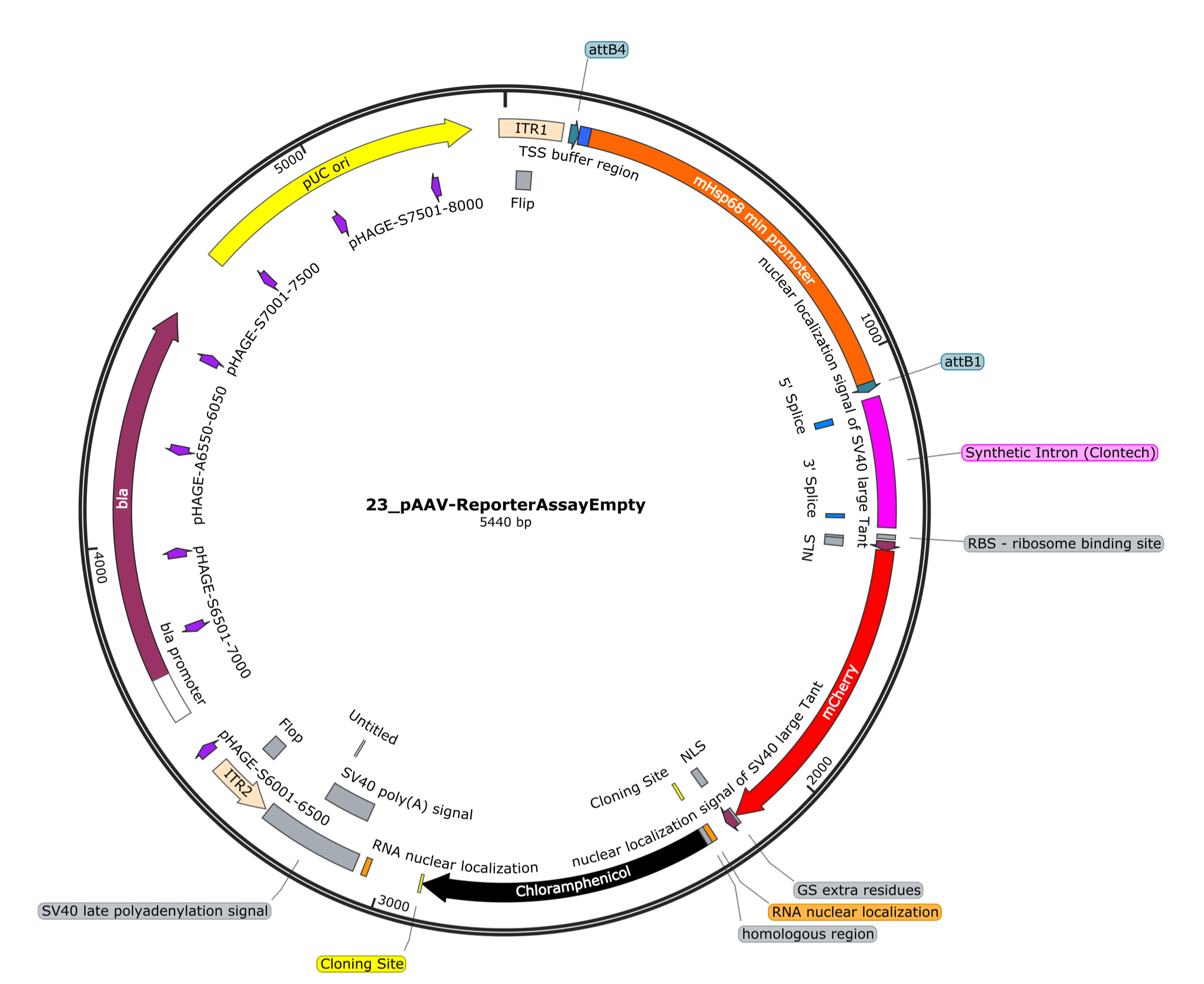

#### Figure S30. Plasmid template for adeno-associated virus-based *in vivo* reporter assays.

Plasmid map for mCherry reporter assay to measure transcriptional activation by candidate enhancer sequences. This plasmid contains inverted terminal repeat (ITR) compatible with adeno-associated virus (AAV) serotype 2-based packaging systems and capsids such as PhP.eB. This empty plasmid has the chloramphenicol resistance gene as the placeholder where candidate enhancers to be tested can be cloned in between multiple cloning sites. The plasmid contains the ampicillin resistance gene *bla*, the Hsp68 minimal promoter, an mRNA stabilizing synthetic intron, a mCherry protein open reading frame, two protein nuclear localization motifs, three mRNA nuclear localization motifs, and a polyA signal.

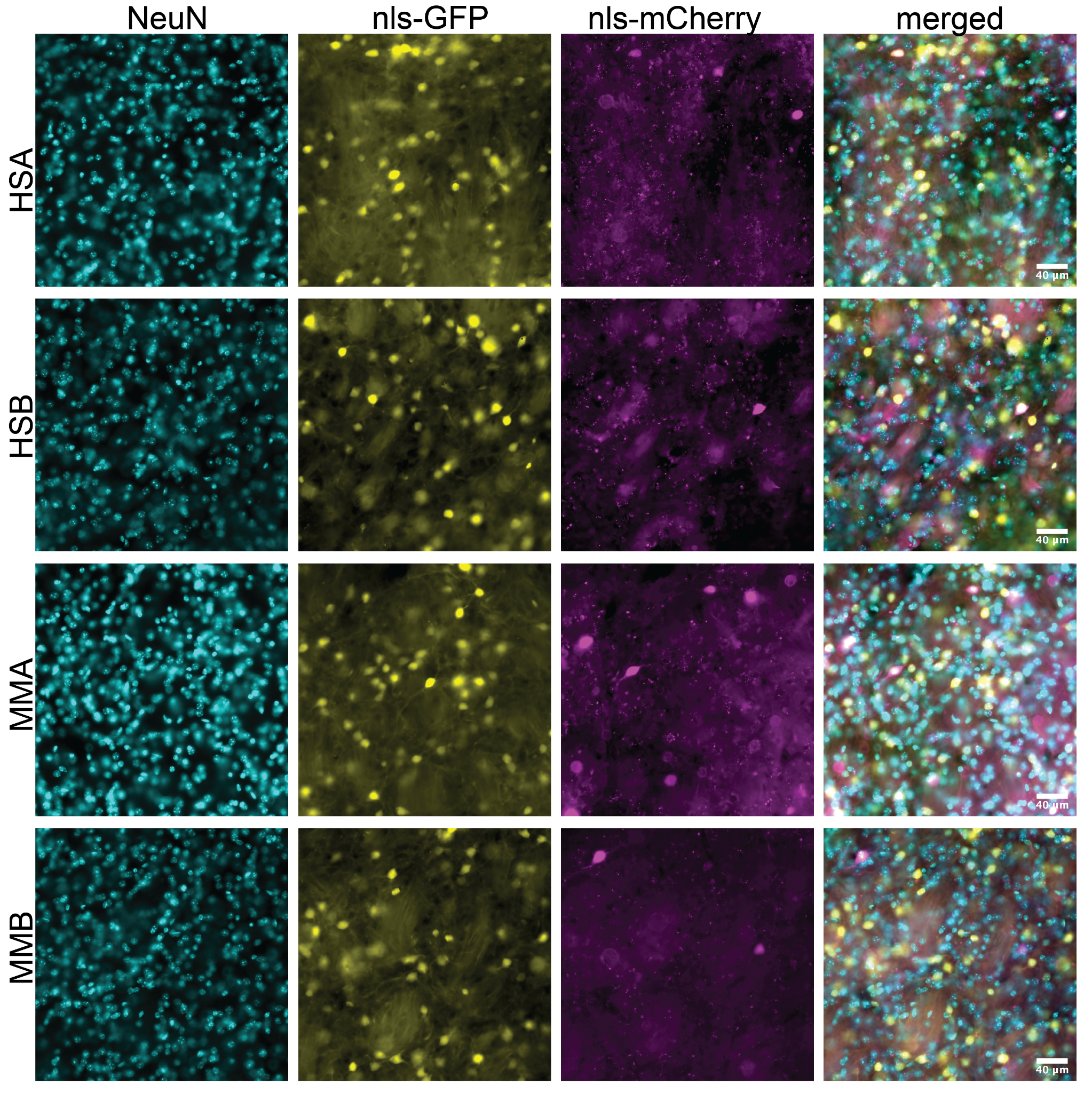

#### Figure S31. An *in vivo* reporter assay to test conserved cell type regulatory activity of candidate enhancers.

Multiplexed fluorescence micrographs acquired at 20X magnification of the mouse caudoputamen with injections of a transduction control virus delivering a nuclear localising (nls) GFP with the reporter assay virus delivering nls-mCherry under the control of a minimal promoter and the candidate enhancer sequence (A or B) from the human (HS) or mouse genomes (MM). The immunofluorescence from left to right label NeuN, GFP, mCherry, and the composite image.

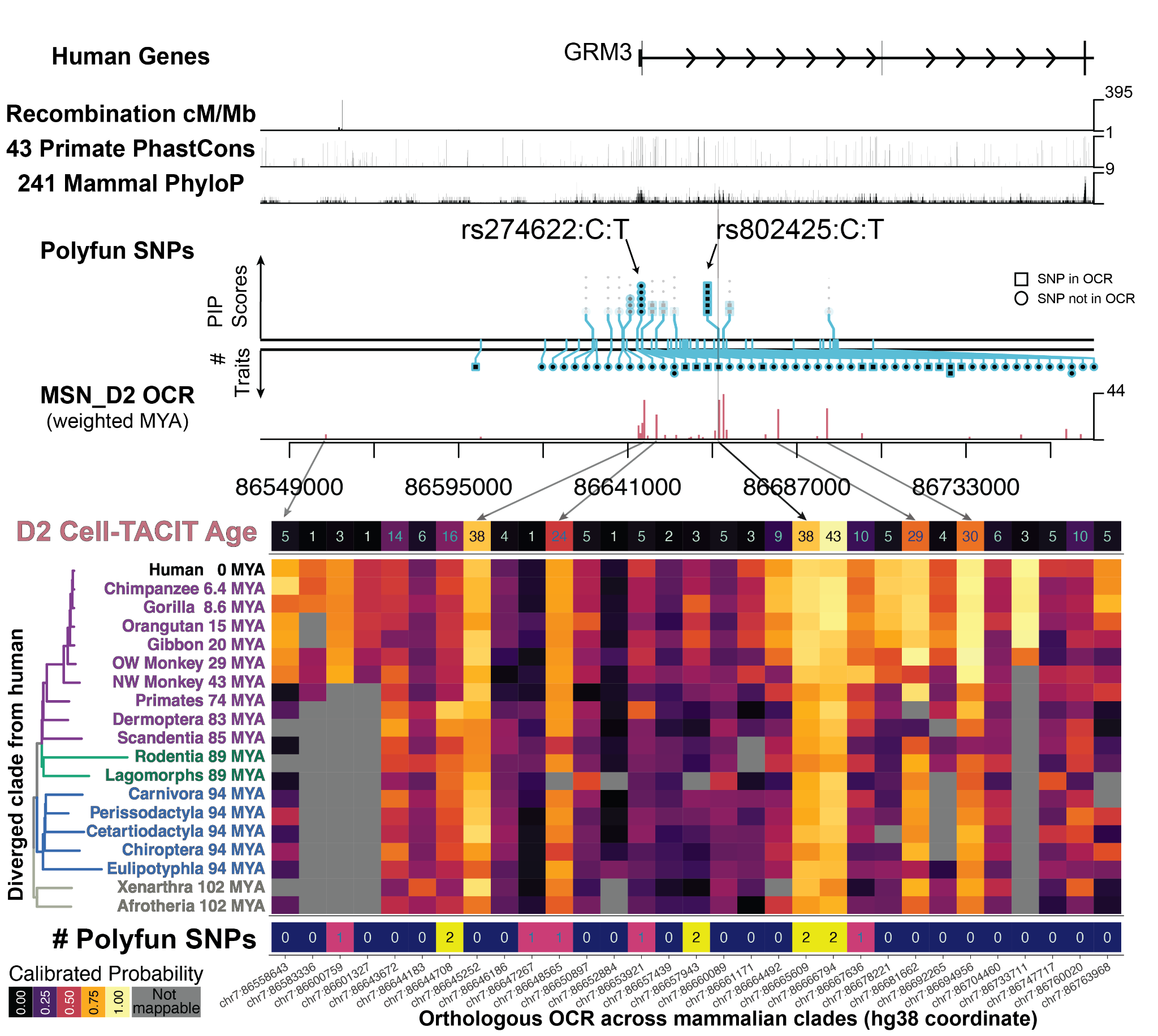

#### Figure S32. CTACIT annotations prioritize fine-mapped SNPs of neuroticism to conserved, active D2 open chromatin regions.

Locus plot at the *GRM3* gene locus with SNPs and CTACIT annotations for D2 MSN models identifies candidate regulatory elements with fine-mapped SNPs from the neuroticism GWAS. CTACIT Age (top row) heatmap for D2 MSNs summarizes the predicted activity of each open chromatin region in the track plot across increasingly distant clades of placental mammals (middle rows). The phylogenetic tree annotates the millions of years ago (MYA) that the common ancestor of each group of placental mammals and humans has diverged. Multiple D2 MSN open chromatin regions contain fine-mapped SNPs in this locus (bottom row). Lollipop plot feature height reflects the maximum PIP of the fine-mapped SNPs, and features are jittered to better visualize clusters of fine-mapped SNPs across a large genomic region. Mb: million bases. Arrows align cell type OCR tracks to corresponding columns in the heatmap. OCRs not mappable to a clade are plotted in gray.

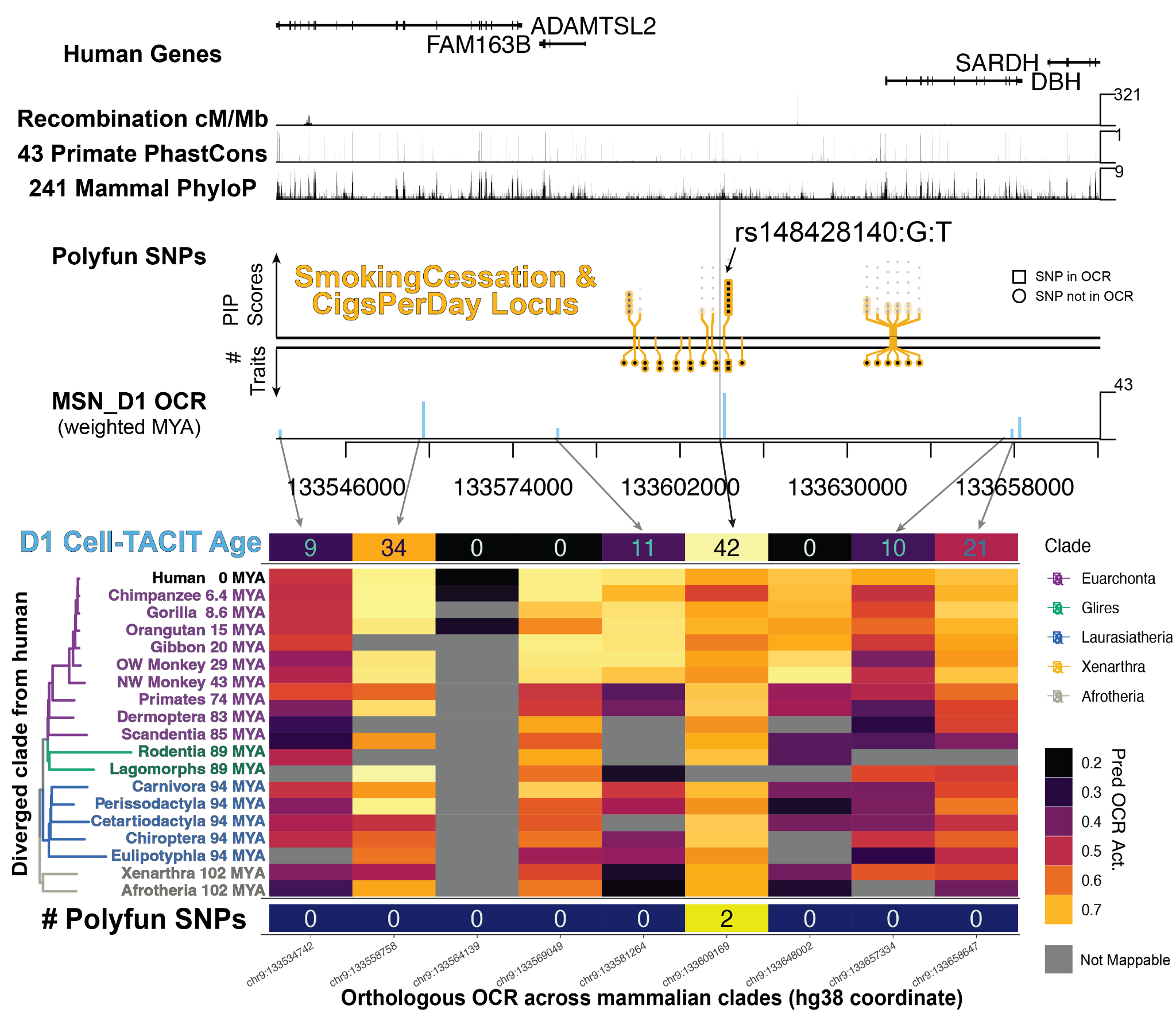

#### Figure S33. CTACIT annotations prioritize fine-mapped SNPs of smoking use traits to conserved, active D1 MSN open chromatin regions.

Locus plot at a dense gene locus including *ADAMTSL2* and other genes with SNPs and CTACIT annotations for D1 MSN models identifies candidate regulatory elements with fine-mapped SNPs from smoking cessation and cigarettes per day GWAS. CTACIT Age (top row) heatmap for D1 MSNs summarizes the predicted activity of each open chromatin region in the track plot across increasingly distant clades of placental mammals (middle rows). The phylogenetic tree annotates the millions of years ago (MYA) that the common ancestor of each group of placental mammals and humans has diverged. One D1 MSN open chromatin region contains fine-mapped SNPs in this locus (bottom row). Lollipop plot feature height reflects the maximum PIP of the fine-mapped SNPs, and features are jittered to better visualize clusters of fine-mapped SNPs across a large genomic region. Mb: million bases. Arrows align cell type OCR tracks to corresponding columns in the heatmap. OCRs not mappable to a clade are plotted in gray.

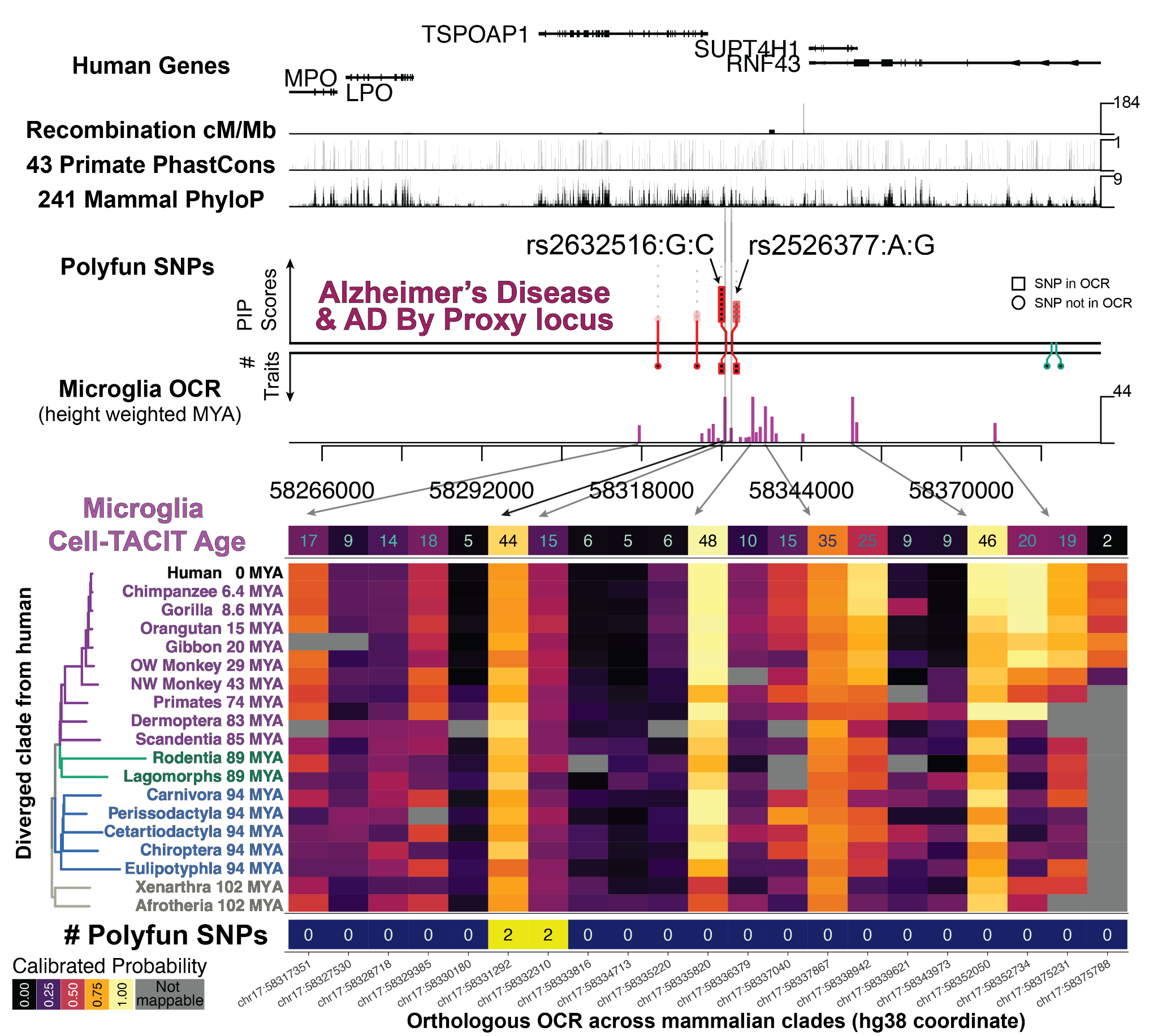

#### Figure S34. CTACIT annotations prioritize fine-mapped SNPs of Alzheimer’s Disease to conserved, active microglia open chromatin regions.

Locus plot at a dense gene locus including *TSPOAP1* and other genes with SNPs and CTACIT annotations for microglia models identifies candidate regulatory elements with fine-mapped SNPs from Alzheimer’s Disease (AD) and AD-by-proxy GWAS. CTACIT Age (top row) heatmap for microglia summarizes the predicted activity of each open chromatin region in the track plot across increasingly distant clades of placental mammals (middle rows). The phylogenetic tree annotates the millions of years ago (MYA) that the common ancestor of each group of placental mammals and humans has diverged. Two microglia open chromatin regions contain fine-mapped SNPs in this locus (bottom row). Lollipop plot feature height reflects the maximum PIP of the fine-mapped SNPs, and features are jittered to better visualize clusters of fine-mapped SNPs across a large genomic region. Mb: million bases. Arrows align cell type OCR tracks to corresponding columns in the heatmap. OCRs not mappable to a clade are plotted in gray.

Supplemental Data (separate files)

##### Data_S1_cross-species_meme-chip_motifs.xlsx

Table of MEME-chip motif enrichments computed in each cell type and source species from cell type OCR sequences before splitting into test, validation, and training sets for CTACIT model training and evaluations.

##### Data_S2_CTACIT_model_validation_performance_values.xlsx

Numerical values for model validation performance**.** Tabs are split by overall model validation set performance for overall positives and negatives, cell type-specific positives and negatives, and species-specific positives and negatives.

##### Data_S3_CTACIT_model_test_performance_values.xlsx

Numerical values for model test performance**.** Tabs are split by overall model test set performance for overall positives and negatives, cell type-specific positives and negatives, and species-specific positives and negatives.

##### Data_S4_list_of_GWAS_traits.xlsx

List of genome-wide association studies (GWAS) analyzed in this work. The references for each GWAS are listed in shorthand in the data column reference and in Methods. The order that these traits appear in this list also dictate the traits that occur in **Figure S11-26**. N_UKB indicates the GWAS contains significant overlap with the UK Biobank and the GWAS has a sample size as indicated for inclusion in polyfun fine-mapping. No entries indicated that the GWAS was not run in polyfun fine-mapping.

##### Data_S5_ldsc_coefficient_caudate_celltype_cross-species.xlsx

Numerical data underlying the linear regression of the points plot in **Figure S10** on the conditional independence effect sizes between OCR sets for each cell type. Term is the linear regression term used Methods. “model_” terms specifies the offset effect based on which genome an OCR set came from, mm10 or rheMac10 relative to hg38. Additional columns are the numerical data underlying **Figure S10** from conditional independent LD score regression analyses across 64 GWAS traits, 8 cell types, and 7 cross-species open chromatin sets. The first tab describes the column names, and each tab is divided by GWAS trait in order as listed in **Data S4**. This orientation was selected for fast filtering and lookup by GWAS.

##### Data_S6_Zoonomia_species.xlsx

The tabs in this data table list the species in the Zoonomia CACTUS, the taxonomic orders used for the 20 clade comparisons.

##### Data_S7_heritability_enrichments_Corces2020_phyloP_CellTACIT-Age.xlsx

Numerical data underlying **Figure S11-26** from heritability enrichment LD score regression analyses across 64 GWAS traits, 8 cell types, and 26 cross-species open chromatin sets. The first tab describes the column names, and each tab is divided up by GWAS trait in order as listed in **Data S4**. This orientation was selected for fast filtering and lookup by GWAS. phyloP: mammalian phyloP constrained (phyloP.cons) or accelerated regions (phyloP.accel), gray bars.

ATAC-seq: cell type OCRs measured in human, human and macaque, or human and mouse.

CTACIT Score: human cell type OCRs mapabble to distant species or mappable and predicted active by CTACIT machine learning models.

CTACIT Age: distal human cell type OCRs binned into the youngest (0-25%) to oldest (75-100%) quartiles of being active across distantly related species.

##### Data_S8_CTACIT-Age_eQTL_and_fine-mapped_SNPs.xlsx

Numerical outputs from logistic regression enrichment analyses of eQTLs or fine-mapped SNPs in CTACIT Age quartiles. These columns are the numerical data for **Figure 3A-C, S28**.

##### Data_S9_polyfun_caudate_finemapped_snps_20210518.xlsx

Output of functionally-informed fine-mapping by polyfun and SuSIE for 43 traits (41 returned SNPs). The first tab describes the column names, and each tab is divided up by GWAS trait in order as listed in **Data S1**. This orientation was selected for fast filtering and lookup by GWAS.

##### Data_S10_CellTACIT_Age_caudate_finemapped_snps_20220718.xlsx

CTACIT Age annotations for polyfun fine-mapped SNPs from **Data S5** if those SNPs overlapped non-exonic human cell type OCRs. Each tab is divided up by the CTACIT age annotated across all traits and fine-mapped SNPs. This orientation was selected for fast filtering and lookup by GWAS. Additional columns here are “cell type” for the source CTACIT Age model, “consFrac” is the proportion of the cell type OCR that has a Zoonomia phyloP score > 2.270, and the relevant “CTACIT Age”.

##### Data_S11_reporter_assay_CellTACIT-predictions.xlsx

CTACIT Age and Score predictions for two candidate enhancers near the *DRD2* locus. These columns are the numerical data for **Figure 4C, D, F**.

##### Data_S12_reporter_assay_stats_and_data.xlsx

Reporter assay segmentation outputs per image and statistical outputs underlying **Figure 4E, F.**

References

1. Davenport AT, Grant KA, Szeliga KT, Friedman DP, Daunais JB. Standardized method for the harvest of nonhuman primate tissue optimized for multiple modes of analyses. Cell Tissue Bank. 2014;15:99–110.

2. Fang R, Preissl S, Li Y, Hou X, Lucero J, Wang X, et al. Comprehensive analysis of single cell ATAC-seq data with SnapATAC. Nat Commun. 2021;12:1337.

3. Martin M. Cutadapt removes adapter sequences from high-throughput sequencing reads [Internet]. EMBnet.journal. 2011. p. 10. Available from: http://dx.doi.org/10.14806/ej.17.1.200

4. Langmead B, Salzberg SL. Fast gapped-read alignment with Bowtie 2. Nat Methods. 2012;9:357–9.

5. Warren WC, Harris RA, Haukness M, Fiddes IT, Murali SC, Fernandes J, et al. Sequence diversity analyses of an improved rhesus macaque genome enhance its biomedical utility. Science [Internet]. 2020;370. Available from: http://dx.doi.org/10.1126/science.abc6617

6. Li H, Handsaker B, Wysoker A, Fennell T, Ruan J, Homer N, et al. The Sequence Alignment/Map format and SAMtools. Bioinformatics. 2009;25:2078–9.

7. Granja JM, Corces MR, Pierce SE, Bagdatli ST, Choudhry H, Chang HY, et al. ArchR is a scalable software package for integrative single-cell chromatin accessibility analysis. Nat Genet. 2021;53:403–11.

8. Shumate A, Salzberg SL. Liftoff: accurate mapping of gene annotations. Bioinformatics [Internet]. 2020; Available from: http://dx.doi.org/10.1093/bioinformatics/btaa1016

9. He J, Kleyman M, Chen J, Alikaya A, Rothenhoefer KM, Ozturk BE, et al. Transcriptional and anatomical diversity of medium spiny neurons in the primate striatum. Curr Biol [Internet]. 2021; Available from: http://dx.doi.org/10.1016/j.cub.2021.10.015

10. Korsunsky I, Millard N, Fan J, Slowikowski K, Zhang F, Wei K, et al. Fast, sensitive and accurate integration of single-cell data with Harmony. Nat Methods. 2019;16:1289–96.

11. Stuart T, Butler A, Hoffman P, Hafemeister C, Papalexi E, Mauck WM, et al. Comprehensive Integration of Single-Cell Data [Internet]. Cell. 2019. p. 1888-1902.e21. Available from: http://dx.doi.org/10.1016/j.cell.2019.05.031

12. Zhang Y, Liu T, Meyer CA, Eeckhoute J, Johnson DS, Bernstein BE, et al. Model-based Analysis of ChIP-Seq (MACS). Genome Biol. 2008;9:R137.

13. Pennacchio LA, Ahituv N, Moses AM, Prabhakar S, Nobrega MA, Shoukry M, et al. In vivo enhancer analysis of human conserved non-coding sequences. Nature [Internet]. 2006 [cited 2022 Mar 20];444. Available from: https://pubmed.ncbi.nlm.nih.gov/17086198/

14. Zanta MA, Belguise-Valladier P, Behr J-P. Gene delivery: A single nuclear localization signal peptide is sufficient to carry DNA to the cell nucleus [Internet]. Proceedings of the National Academy of Sciences. 1999. p. 91–6. Available from: http://dx.doi.org/10.1073/pnas.96.1.91

15. Zhang B, Gunawardane L, Niazi F, Jahanbani F, Chen X, Valadkhan S. A Novel RNA Motif Mediates the Strict Nuclear Localization of a Long Noncoding RNA. Mol Cell Biol. 2014;34:2318.

16. Srinivasan C, Phan BN, Lawler AJ, Ramamurthy E, Kleyman M, Brown AR, et al. Addiction-associated genetic variants implicate brain cell type- and region-specific cis-regulatory elements in addiction neurobiology. J Neurosci [Internet]. 2021; Available from: http://dx.doi.org/10.1523/JNEUROSCI.2534-20.2021

17. Lawler AJ, Brown AR, Bouchard RS, Toong N, Kim Y, Velraj N, et al. Cell Type-Specific Oxidative Stress Genomic Signatures in the Globus Pallidus of Dopamine-Depleted Mice. J Neurosci. 2020;40:9772–83.

18. Chan KY, Jang MJ, Yoo BB, Greenbaum A, Ravi N, Wu W-L, et al. Engineered AAVs for efficient noninvasive gene delivery to the central and peripheral nervous systems. Nat Neurosci. 2017;20:1172–9.

19. Graveland GA, DiFiglia M. The frequency and distribution of medium-sized neurons with indented nuclei in the primate and rodent neostriatum. Brain Res [Internet]. 1985 [cited 2022 July 11];327. Available from: https://pubmed.ncbi.nlm.nih.gov/3986508/

20. Corces MR, Shcherbina A, Kundu S, Gloudemans MJ, Frésard L, Granja JM, et al. Single-cell epigenomic analyses implicate candidate causal variants at inherited risk loci for Alzheimer’s and Parkinson’s diseases. Nat Genet. 2020;52:1158–68.

21. Li YE, Preissl S, Hou X, Zhang Z, Zhang K, Qiu Y, et al. An atlas of gene regulatory elements in adult mouse cerebrum. Nature. 2021;598:129–36.

22. Kaplow IM, Schäffer DE, Wirthlin ME, Lawler AJ, Brown AR, Kleyman M, et al. Inferring mammalian tissue-specific regulatory conservation by predicting tissue-specific differences in open chromatin. BMC Genomics [Internet]. 2022 [cited 2022 July 5];23. Available from: https://pubmed.ncbi.nlm.nih.gov/35410163/

23. Wirthlin, Morgan, Kaplow, Irene M., Lawler, Alyssa J., He, J., Phan., BaDoi, N., Brown, Ashley R., Stauffer, William R., Pfenning, Andreas R. The Regulatory Evolution of the Primate Fine-Motor System. 2020.

24. Kaplow IM, Lawler AJ, Schäffer DE, Srinivasan C, Sestili HH, Wirthlin ME, et al. Relating enhancer genetic variation across mammals to complex phenotypes using machine learning. Science. 2023;Apr 28;380:eabm7993.

25. Wirthlin ME, Zhang X, Kaplow IM, Schäffer DE, Lawler AJ, Schmid T, et al. Vocal learning-associated convergent evolution in mammalian regulatory elements and proteins. 2021;

26. Halstead MM, Kern C, Saelao P, Wang Y, Chanthavixay G, Medrano JF, et al. A comparative analysis of chromatin accessibility in cattle, pig, and mouse tissues. BMC Genomics. 2020;21:698.

27. Armstrong J, Hickey G, Diekhans M, Fiddes IT, Novak AM, Deran A, et al. Progressive Cactus is a multiple-genome aligner for the thousand-genome era. Nature. 2020;587:246–51.

28. Zhang X, Kaplow IM, Wirthlin M, Park TY, Pfenning AR. HALPER facilitates the identification of regulatory element orthologs across species. Bioinformatics. 2020;36:4339–40.

29. Genereux DP, Seres A, Armstrong J, Johnson J, Marinescu VD, Murén E, et al. A comparative genomics multitool for scientific discovery and conservation. Nature. 2020;587:240–5.

30. Hickey G, Paten B, Earl D, Zerbino D, Haussler D. HAL: a hierarchical format for storing and analyzing multiple genome alignments. Bioinformatics. 2013;29:1341–2.

31. Kent WJ, Baertsch R, Hinrichs A, Miller W, Haussler D. Evolution’s cauldron: duplication, deletion, and rearrangement in the mouse and human genomes. Proc Natl Acad Sci U S A. 2003;100:11484–9.

32. Finucane HK, Bulik-Sullivan B, Gusev A, Trynka G, Reshef Y, Loh P-R, et al. Partitioning heritability by functional annotation using genome-wide association summary statistics. Nat Genet. 2015;47:1228.

33. Finucane HK, Reshef YA, Anttila V, Slowikowski K, Gusev A, Byrnes A, et al. Heritability enrichment of specifically expressed genes identifies disease-relevant tissues and cell types. Nat Genet. 2018;50:621–9.

34. Consortium 1000 Genomes Project, Auton A, Brooks LD, Durbin RM, Garrison EP, Kang HM, et al. A global reference for human genetic variation. Nature. 2015;526:68–74.

35. Bulik-Sullivan B, Finucane HK, Anttila V, Gusev A, Day FR, Loh P-R, et al. An atlas of genetic correlations across human diseases and traits. Nat Genet. 2015;47:1236–41.

36. Dey KK, van de Geijn B, Kim SS, Hormozdiari F, Kelley DR, Price AL. Evaluating the informativeness of deep learning annotations for human complex diseases. Nat Commun. 2020;11:4703.

37. Jansen IE, Savage JE, Watanabe K, Bryois J, Williams DM, Steinberg S, et al. Genome-wide meta-analysis identifies new loci and functional pathways influencing Alzheimer’s disease risk. Nat Genet. 2019;51:404–13.

38. van Rheenen W, Shatunov A, Dekker AM, McLaughlin RL, Diekstra FP, Pulit SL, et al. Genome-wide association analyses identify new risk variants and the genetic architecture of amyotrophic lateral sclerosis. Nat Genet. 2016;48:1043–8.

39. Fritsche LG, Igl W, Bailey JNC, Grassmann F, Sengupta S, Bragg-Gresham JL, et al. A large genome-wide association study of age-related macular degeneration highlights contributions of rare and common variants. Nat Genet. 2016;48:134–43.

40. Nalls MA, Blauwendraat C, Vallerga CL, Heilbron K, Bandres-Ciga S, Chang D, et al. Identification of novel risk loci, causal insights, and heritable risk for Parkinson’s disease: a meta-analysis of genome-wide association studies. Lancet Neurol. 2019;18:1091–102.

41. Kunkle BW, Grenier-Boley B, Sims R, Bis JC, Damotte V, Naj AC, et al. Genetic meta-analysis of diagnosed Alzheimer’s disease identifies new risk loci and implicates Aβ, tau, immunity and lipid processing. Nat Genet. 2019;51:414–30.

42. Demontis D, Walters RK, Martin J, Mattheisen M, Als TD, Agerbo E, et al. Discovery of the first genome-wide significant risk loci for attention deficit/hyperactivity disorder. Nat Genet. 2019;51:63–75.

43. Watson HJ, Yilmaz Z, Thornton LM, Hübel C, Coleman JRI, Gaspar HA, et al. Genome-wide association study identifies eight risk loci and implicates metabo-psychiatric origins for anorexia nervosa. Nat Genet. 2019;51:1207–14.

44. Mullins N, Forstner AJ, O’Connell KS, Coombes B, Coleman JRI, Qiao Z, et al. Genome-wide association study of more than 40,000 bipolar disorder cases provides new insights into the underlying biology. Nat Genet. 2021;53:817–29.

45. Howard DM, Adams MJ, Clarke T-K, Hafferty JD, Gibson J, Shirali M, et al. Genome-wide meta-analysis of depression identifies 102 independent variants and highlights the importance of the prefrontal brain regions. Nat Neurosci. 2019;22:343–52.

46. Wray NR, Ripke S, Mattheisen M, Trzaskowski M, Byrne EM, Abdellaoui A, et al. Genome-wide association analyses identify 44 risk variants and refine the genetic architecture of major depression. Nat Genet. 2018;50:668–81.

47. (iocdf-Gc) IOCDFGC, (ocgas) OCGAS. Revealing the complex genetic architecture of obsessive-compulsive disorder using meta-analysis. Mol Psychiatry. 2018;23:1181–8.

48. Nievergelt CM, Maihofer AX, Klengel T, Atkinson EG, Chen C-Y, Choi KW, et al. International meta-analysis of PTSD genome-wide association studies identifies sex- and ancestry-specific genetic risk loci. Nat Commun. 2019;10:4558.

49. Cross-Disorder Group of the Psychiatric Genomics Consortium. Electronic address:, Cross-Disorder Group of the Psychiatric Genomics Consortium. Genomic Relationships, Novel Loci, and Pleiotropic Mechanisms across Eight Psychiatric Disorders. Cell. 2019;179:1469-1482.e11.

50. Consortium TSWG of TPG, The Schizophrenia Working Group of the Psychiatric Genomics Consortium, Ripke S, Walters JTR, O’Donovan MC. Mapping genomic loci prioritises genes and implicates synaptic biology in schizophrenia. Nature [Internet]. 2022; Available from: http://dx.doi.org/10.1038/s41586-022-04434-5

51. Sanchez-Roige S, Palmer AA, Fontanillas P, Elson SL, 23andMe Research Team, the Substance Use Disorder Working Group of the Psychiatric Genomics Consortium, Adams MJ, et al. Genome-Wide Association Study Meta-Analysis of the Alcohol Use Disorders Identification Test (AUDIT) in Two Population-Based Cohorts. Am J Psychiatry. 2019;176:107–18.

52. Liu M, Jiang Y, Wedow R, Li Y, Brazel DM, Chen F, et al. Association studies of up to 1.2 million individuals yield new insights into the genetic etiology of tobacco and alcohol use. Nat Genet. 2019;51:237–44.

53. Pasman JA, Verweij KJH, Gerring Z, Stringer S, Sanchez-Roige S, Treur JL, et al. GWAS of lifetime cannabis use reveals new risk loci, genetic overlap with psychiatric traits, and a causal influence of schizophrenia. Nat Neurosci. 2018;21:1161–70.

54. Coffee, Consortium CG, Cornelis MC, Byrne EM, Esko T, Nalls MA, et al. Genome-wide meta-analysis identifies six novel loci associated with habitual coffee consumption. Mol Psychiatry. 2015;20:647–56.

55. Polimanti R, Walters RK, Johnson EC, McClintick JN, Adkins AE, Adkins DE, et al. Leveraging genome-wide data to investigate differences between opioid use vs. opioid dependence in 41,176 individuals from the Psychiatric Genomics Consortium [Internet]. Available from: http://dx.doi.org/10.1101/765065

56. Cabana-Domínguez J, Shivalikanjli A, Fernàndez-Castillo N, Cormand B. Genome-wide association meta-analysis of cocaine dependence: Shared genetics with comorbid conditions [Internet]. Available from: http://dx.doi.org/10.1101/374553

57. Walters RK, Polimanti R, Johnson EC, McClintick JN, Adams MJ, Adkins AE, et al. Transancestral GWAS of alcohol dependence reveals common genetic underpinnings with psychiatric disorders. Nat Neurosci. 2018;21:1656–69.

58. Johnson EC, Demontis D, Thorgeirsson TE, Walters RK, Polimanti R, Hatoum AS, et al. A large-scale genome-wide association study meta-analysis of cannabis use disorder. Lancet Psychiatry. 2020;7:1032–45.

59. Lee JJ, Wedow R, Okbay A, Kong E, Maghzian O, Zacher M, et al. Gene discovery and polygenic prediction from a genome-wide association study of educational attainment in 1.1 million individuals. Nat Genet. 2018;50:1112–21.

60. Hill WD, Davies NM, Ritchie SJ, Skene NG, Bryois J, Bell S, et al. Genome-wide analysis identifies molecular systems and 149 genetic loci associated with income. Nat Commun. 2019;10:5741.

61. Pulit SL, Stoneman C, Morris AP, Wood AR, Glastonbury CA, Tyrrell J, et al. Meta-analysis of genome-wide association studies for body fat distribution in 694 649 individuals of European ancestry. Hum Mol Genet. 2019;28:166–74.

62. Jansen PR, Nagel M, Watanabe K, Wei Y, Savage JE, de Leeuw CA, et al. Genome-wide meta-analysis of brain volume identifies genomic loci and genes shared with intelligence. Nat Commun. 2020;11:5606.

63. Nagel M, Jansen PR, Stringer S, Watanabe K, de Leeuw CA, Bryois J, et al. Meta-analysis of genome-wide association studies for neuroticism in 449,484 individuals identifies novel genetic loci and pathways. Nat Genet. 2018;50:920–7.

64. Savage JE, Jansen PR, Stringer S, Watanabe K, Bryois J, de Leeuw CA, et al. Genome-wide association meta-analysis in 269,867 individuals identifies new genetic and functional links to intelligence. Nat Genet. 2018;50:912–9.

65. Karlsson Linnér R, Biroli P, Kong E, Meddens SFW, Wedow R, Fontana MA, et al. Genome-wide association analyses of risk tolerance and risky behaviors in over 1 million individuals identify hundreds of loci and shared genetic influences. Nat Genet. 2019;51:245–57.

66. Johnston KJA, Adams MJ, Nicholl BI, Ward J, Strawbridge RJ, Ferguson A, et al. Genome-wide association study of multisite chronic pain in UK Biobank. PLoS Genet. 2019;15:e1008164.

67. Meng W, Adams MJ, Hebert HL, Deary IJ, McIntosh AM, Smith BH. A Genome-Wide Association Study Finds Genetic Associations with Broadly-Defined Headache in UK Biobank (N=223,773). EBioMedicine. 2018;28:180–6.

68. Wang H, Lane JM, Jones SE, Dashti HS, Ollila HM, Wood AR, et al. Genome-wide association analysis of self-reported daytime sleepiness identifies 42 loci that suggest biological subtypes. Nat Commun. 2019;10:3503.

69. Jansen PR, Watanabe K, Stringer S, Skene N, Bryois J, Hammerschlag AR, et al. Genome-wide analysis of insomnia in 1,331,010 individuals identifies new risk loci and functional pathways. Nat Genet. 2019;51:394–403.

70. Dashti HS, Jones SE, Wood AR, Lane JM, van Hees VT, Wang H, et al. Genome-wide association study identifies genetic loci for self-reported habitual sleep duration supported by accelerometer-derived estimates. Nat Commun. 2019;10:1100.

71. Wuttke M, Li Y, Li M, Sieber KB, Feitosa MF, Gorski M, et al. A catalog of genetic loci associated with kidney function from analyses of a million individuals. Nat Genet. 2019;51:957–72.

72. Willer CJ, Schmidt EM, Sengupta S, Peloso GM, Gustafsson S, Kanoni S, et al. Discovery and refinement of loci associated with lipid levels. Nat Genet. 2013;45:1274–83.

73. Xue A, Wu Y, Zhu Z, Zhang F, Kemper KE, Zheng Z, et al. Genome-wide association analyses identify 143 risk variants and putative regulatory mechanisms for type 2 diabetes. Nat Commun. 2018;9:2941.

74. Kemp JP, Morris JA, Medina-Gomez C, Forgetta V, Warrington NM, Youlten SE, et al. Identification of 153 new loci associated with heel bone mineral density and functional involvement of GPC6 in osteoporosis. Nat Genet. 2017;49:1468–75.

75. Howson JMM, Zhao W, Barnes DR, Ho W-K, Young R, Paul DS, et al. Fifteen new risk loci for coronary artery disease highlight arterial-wall-specific mechanisms. Nat Genet. 2017;49:1113–9.

76. Wood AR, Esko T, Yang J, Vedantam S, Pers TH, Gustafsson S, et al. Defining the role of common variation in the genomic and biological architecture of adult human height. Nat Genet. 2014;46:1173–86.

77. Zillikens MC, Demissie S, Hsu Y-H, Yerges-Armstrong LM, Chou W-C, Stolk L, et al. Large meta-analysis of genome-wide association studies identifies five loci for lean body mass. Nat Commun. 2017;8:80.

78. Timmers PR, Mounier N, Lall K, Fischer K, Ning Z, Feng X, et al. Genomics of 1 million parent lifespans implicates novel pathways and common diseases and distinguishes survival chances. Elife [Internet]. 2019;8. Available from: http://dx.doi.org/10.7554/eLife.39856

79. Chen L, Fish AE, Capra JA. Prediction of gene regulatory enhancers across species reveals evolutionarily conserved sequence properties. PLoS Comput Biol. 2018;14:e1006484.

80. Kelley DR. Cross-species regulatory sequence activity prediction. PLoS Comput Biol. 2020;16:e1008050.

81. Grossman SR, Engreitz J, Ray JP, Nguyen TH, Hacohen N, Lander ES. Positional specificity of different transcription factor classes within enhancers. Proc Natl Acad Sci U S A. 2018;115:E7222–30.

82. Vandel J, Cassan O, Lèbre S, Lecellier C-H, Bréhélin L. Probing transcription factor combinatorics in different promoter classes and in enhancers. BMC Genomics. 2019;20:103.

83. Architecture of the human regulatory network derived from ENCODE data [Internet]. Nature 2012 p. 91–100. Available from: https://www.nature.com/articles/nature11245.pdf

84. Khan A, Riudavets Puig R, Boddie P, Mathelier A. BiasAway: command-line and web server to generate nucleotide composition-matched DNA background sequences. Bioinformatics. 2021;37:1607–9.

85. Blair Hedges S, Kumar S. The Timetree of Life. OUP Oxford; 2009.

86. Kumar S, Stecher G, Suleski M, Hedges SB. TimeTree: A Resource for Timelines, Timetrees, and Divergence Times. Mol Biol Evol. 2017;34:1812–9.

87. Foley NM, Mason VC, Harris AJ, Breder KR, Damas J, Lewin HA, et al. A genomic timescale for placental mammal evolution. 2021;

88. Yu G, Lam TT-Y, Zhu H, Guan Y. Two Methods for Mapping and Visualizing Associated Data on Phylogeny Using Ggtree. Mol Biol Evol. 2018;35:3041–3.

89. Wang L-G, Lam TT-Y, Xu S, Dai Z, Zhou L, Feng T, et al. Treeio: An R Package for Phylogenetic Tree Input and Output with Richly Annotated and Associated Data. Mol Biol Evol. 2020;37:599–603.

90. Consortium TG, The GTEx Consortium. The GTEx Consortium atlas of genetic regulatory effects across human tissues [Internet]. Science. 2020. p. 1318–30. Available from: http://dx.doi.org/10.1126/science.aaz1776

91. Bryois J, Calini D, Macnair W, Foo L, Urich E, Ortmann W, et al. Cell-type specific cis-eQTLs in eight brain cell-types identifies novel risk genes for human brain disorders [Internet]. Available from: http://dx.doi.org/10.1101/2021.10.09.21264604

92. Benjamini Y, Hochberg Y. Controlling the false discovery rate: a practical and powerful approach to multiple testing. J R Stat Soc [Internet]. 1995; Available from: https://rss.onlinelibrary.wiley.com/doi/abs/10.1111/j.2517-6161.1995.tb02031.x

93. Enea M. Fitting Linear and Generalized Linear Models to Large Data Sets [R package speedglm version 0.3-4]. 2022 [cited 2022 July 16]; Available from: https://CRAN.R-project.org/package=speedglm

94. Weissbrod O, Hormozdiari F, Benner C, Cui R, Ulirsch J, Gazal S, et al. Functionally informed fine-mapping and polygenic localization of complex trait heritability. Nat Genet. 2020;52:1355–63.

95. Sullivan PF, Meadows JRS, Gazal S, Phan BN, Li X, Genereux DP, et al. Leveraging base-pair mammalian constraint to understand genetic variation and human disease. Science. 2023;380:eabn2937.

96. ENCODE Project Consortium, Moore JE, Purcaro MJ, Pratt HE, Epstein CB, Shoresh N, et al. Expanded encyclopaedias of DNA elements in the human and mouse genomes. Nature. 2020;583:699–710.

97. Wang G, Sarkar A, Carbonetto P, Stephens M. A simple new approach to variable selection in regression, with application to genetic fine mapping [Internet]. Journal of the Royal Statistical Society: Series B (Statistical Methodology). 2020. p. 1273–300. Available from: http://dx.doi.org/10.1111/rssb.12388

98. Nasser J, Bergman DT, Fulco CP, Guckelberger P, Doughty BR, Patwardhan TA, et al. Genome-wide enhancer maps link risk variants to disease genes. Nature. 2021;593:238–43.

99. Lambert JT, Su-Feher L, Cichewicz K, Warren TL, Zdilar I, Wang Y, et al. Parallel functional testing identifies enhancers active in early postnatal mouse brain. Elife [Internet]. 2021;10. Available from: http://dx.doi.org/10.7554/eLife.69479

100. Greenwald NF, Miller G, Moen E, Kong A, Kagel A, Dougherty T, et al. Whole-cell segmentation of tissue images with human-level performance using large-scale data annotation and deep learning. Nat Biotechnol. 2021;40:555–65.

101. van der Walt S, Schönberger JL, Nunez-Iglesias J, Boulogne F, Warner JD, Yager N, et al. scikit-image: image processing in Python. PeerJ. 2014;2:e453.

102. Virtanen P, Gommers R, Oliphant TE, Haberland M, Reddy T, Cournapeau D, et al. SciPy 1.0: fundamental algorithms for scientific computing in Python. Nat Methods. 2020;17:261–72.

103. Bates D, Mächler M, Bolker B, Walker S. Fitting Linear Mixed-Effects Models Using**lme4** [Internet]. Journal of Statistical Software. 2015. Available from: http://dx.doi.org/10.18637/jss.v067.i01

104. Roller M, Stamper E, Villar D, Izuogu O, Martin F, Redmond AM, et al. LINE retrotransposons characterize mammalian tissue-specific and evolutionarily dynamic regulatory regions. Genome Biol. 2021;22:62.

105. Sullivan PF, Agrawal A, Bulik CM, Andreassen OA, Børglum AD, Breen G, et al. Psychiatric Genomics: An Update and an Agenda. Am J Psychiatry. 2018;175:15.

106. Gjoneska E, Pfenning AR, Mathys H, Quon G, Kundaje A, Tsai L-H, et al. Conserved epigenomic signals in mice and humans reveal immune basis of Alzheimer’s disease. Nature. 2015;518:365–9.

107. Latini A, Ciccacci C, Benedittis GD, Novelli L, Ceccarelli F, Conti F, et al. Altered expression of miR-142, miR-155, miR-499a and of their putative common target in systemic lupus erythematosus. Epigenomics. 2021;13:5–13.

108. Ghanbari M, Munshi ST, Ma B, Lendemeijer B, Bansal S, Adams HH, et al. A functional variant in the miR-142 promoter modulating its expression and conferring risk of Alzheimer disease. Hum Mutat. 2019;40:2131–45.

109. Kim T-K, Hemberg M, Gray JM, Costa AM, Bear DM, Wu J, et al. Widespread transcription at neuronal activity-regulated enhancers. Nature. 2010;465:182–7.

110. Mo A, Mukamel EA, Davis FP, Luo C, Henry GL, Picard S, et al. Epigenomic Signatures of Neuronal Diversity in the Mammalian Brain. Neuron. 2015;86:1369–84.

111. Buenrostro JD, Wu B, Litzenburger UM, Ruff D, Gonzales ML, Snyder MP, et al. Single-cell chromatin accessibility reveals principles of regulatory variation. Nature. 2015;523:486–90.

112. Klein JC, Agarwal V, Inoue F, Keith A, Martin B, Kircher M, et al. A systematic evaluation of the design and context dependencies of massively parallel reporter assays. Nat Methods [Internet]. 2020 [cited 2022 July 17];17. Available from: https://pubmed.ncbi.nlm.nih.gov/33046894/

113. Lawler AJ, Ramamurthy E, Brown AR, Shin N, Kim Y, Toong N, et al. Machine learning sequence prioritization for cell type-specific enhancer design. Elife [Internet]. 2022;11. Available from: http://dx.doi.org/10.7554/eLife.69571

114. Gemberling MP, Siklenka K, Rodriguez E, Tonn-Eisinger KR, Barrera A, Liu F, et al. Transgenic mice for in vivo epigenome editing with CRISPR-based systems. Nat Methods. 2021;18:965–74.

115. Stojanovic T, Orlova M, Sialana FJ, Höger H, Stuchlik S, Milenkovic I, et al. Validation of dopamine receptor DRD1 and DRD2 antibodies using receptor deficient mice. Amino Acids. 2017;49:1101–9.

116. Ren K, Guo B, Dai C, Yao H, Sun T, Liu X, et al. Striatal Distribution and Cytoarchitecture of Dopamine Receptor Subtype 1 and 2: Evidence from Double-Labeling Transgenic Mice. Front Neural Circuits. 2017;11:57.

117. Gerfen CR, Paletzki R, Heintz N. GENSAT BAC Cre-Recombinase Driver Lines to Study the Functional Organization of Cerebral Cortical and Basal Ganglia Circuits [Internet]. Neuron. 2013. p. 1368–83. Available from: http://dx.doi.org/10.1016/j.neuron.2013.10.016

118. Öztürk BE, Johnson ME, Kleyman M, Turunç S, He J, Jabalameli S, et al. scAAVengr, a transcriptome-based pipeline for quantitative ranking of engineered AAVs with single-cell resolution. Elife [Internet]. 2021;10. Available from: http://dx.doi.org/10.7554/eLife.64175

119. Hrvatin S, Tzeng CP, Nagy MA, Stroud H, Koutsioumpa C, Wilcox OF, et al. A scalable platform for the development of cell-type-specific viral drivers. Elife [Internet]. 2019;8. Available from: http://dx.doi.org/10.7554/eLife.48089

120. Phan BN, Bohlen JF, Davis BA, Ye Z, Chen H-Y, Mayfield B, et al. A myelin-related transcriptomic profile is shared by Pitt-Hopkins syndrome models and human autism spectrum disorder. Nat Neurosci. 2020;23:375–85.

121. Halldorsson BV, Palsson G, Stefansson OA, Jonsson H, Hardarson MT, Eggertsson HP, et al. Characterizing mutagenic effects of recombination through a sequence-level genetic map. Science [Internet]. 2019;363. Available from: http://dx.doi.org/10.1126/science.aau1043

122. Christmas MJ, Kaplow IM, Genereux DP, Dong MX, Hughes GM, Li X, et al. Evolutionary constraint and innovation across hundreds of placental mammals. Science. 2023;380:eabn3943.

123. Ou J, Zhu LJ. trackViewer: a Bioconductor package for interactive and integrative visualization of multi-omics data. Nat Methods. 2019;16:453–4.

124. The Schizophrenia Working Group of the Psychiatric Genomics Consortium, Ripke S, Walters JTR, O’Donovan MC. Mapping genomic loci prioritises genes and implicates synaptic biology in schizophrenia. medRxiv. 2020;2020.09.12.20192922.
